# The SARS-CoV-2 Spike S1 Protein Induces Global Proteomic Changes in ATII-Like Rat L2 Cells that are Attenuated by Hyaluronan

**DOI:** 10.1101/2022.08.31.506023

**Authors:** James A. Mobley, Adam Molyvdas, Kyoko Kojima, Tamas Jilling, Jian-Liang Li, Stavros Garantziotis, Sadis Matalon

## Abstract

The COVID-19 pandemic continues to impose a major impact on global health and economy since its identification in early 2020, causing significant morbidity and mortality worldwide. Caused by the SARS-CoV-2 virus, along with a growing number of variants that have been characterized to date, COVID-19 has led to 571,198,904 confirmed cases, and 6,387,863 deaths worldwide (as of July 15^th^, 2022). Despite tremendous advances in our understanding of COVID19 pathogenesis, the precise mechanism by which SARS-CoV2 causes epithelial injury is incompletely understood. In this current study, robust application of global-discovery proteomics applications combined with systems biology analysis identified highly significant induced changes by the Spike S1 protein of SARS-CoV-2 in an ATII-like Rat L2 cells that include three significant network hubs: E2F1, CREB1/ RelA, and ROCK2/ RhoA. Separately, we found that pre-treatment with High Molecular Weight Hyaluronan (HMW-HA), greatly attenuated the S1 effects. Immuno-targeted studies carried out on E2F1 and Rock2/ RhoA induction and kinase-mediated activation, in addition to cell cycle measurements, validated these observations. Taken as a whole, our discovery proteomics and systems analysis workflow, combined with standard immuno-targeted and cell cycle measurements revealed profound and novel biological changes that contribute to our current understanding of both Spike S1 and Hyaluronan biology. This data shows that the Spike S1 protein may contribute to epithelial injury induced by SARS-CoV-2. In addition, our work supports the potential benefit of HMW-HA in ameliorating SARS CoV2 induced cell injury.

## Introduction

Coronavirus disease 2019 (COVID-19) is caused by a new beta-coronavirus, SARS-CoV-2; its epi-center was in Wuhan China. COVID-19 was identified as a pandemic by the World Health Organization (WHO) on March 11, 2020. As of 8/11/2022, there have been 592,703,091 million cases worldwide resulting in 6,447,162 deaths (https://www.worldometers.info/coronavirus/). This far exceeds the total deaths caused by both SARS and Middle Eastern respiratory syndrome (MERS). A large fraction of COVID-19 patients develop ARDS within 24-48 hours after onset of symptoms with more than a 50% mortality rate (67).

COVID-19 is caused by an enveloped, non-segmented, positive sense RNA new beta-coronavirus, SARS-CoV-2; it belongs to the sarbecovirus, *orthornaviridae* subfamily which is broadly distributed in humans and other mammals (26). It contains four main structural proteins including three glycoproteins (spike (S), envelope (E) and a membrane (M)), a nucleocapsid (N) protein, and several accessory proteins (33). The S glycoprotein is a transmembrane protein with a molecular weight of about 150 kDa, found in the outer portion of the virus. It forms homotrimers protruding in the viral surface and facilitates binding of SARS CoV-2 to human ACE2 and CD147 receptors of host cells (38). The SARS CoV-2 S protein has significant homology with the SARS-CoV-1 (51), but binds with higher affinity to ACE2 receptors (66) due to a furin-like cleavage site (^682^RRAR/S^686^) inserted in the S1/S2 protease cleavage site (15). The S1 region of the Spike protein is responsible for binding to the host cell ACE2 receptor, whereas the S2 region is responsible for fusion of the viral RNA and cellular membrane. SARS-CoV-2 can infect several different animal species like rats, ferrets, hamsters, non-human primates, minks, tree shrews, raccoon dogs, fruit bats, and rabbits (8).

Recently there have been reports that the SARS-CoV-2 spike protein interacts with other receptors as well, like CD147(63). Despite the introduction of several effective vaccines, the mutagenicity and transmissibility of SARS-CoV-2, as evidenced by the emergence of several variants, makes the understanding of the pathophysiological mechanism involved and the identification of therapeutic agents an extremely high priority (55). COVID-19 is mainly a respiratory disease but it can lead to other systemic effects and eventually to a systemic disorder (29). SARS-CoV-2 exhibits a very broad tissue tropism as it can infect and affect a variety of different organs and systems. Among the systems affected are the musculoskeletal system(46), nervous system(28), kidneys(49), and the cardiovascular system (29)

The pathophysiology and immune response to SARS-CoV-2 infection is very broad and involves signaling pathways from several cell and tissue types (54). In particular, while it is understood that SARS-CoV2 utilized S-protein binding to ACE2 in order to enter cells, it is now increasingly reported that other pathways, involving innate immunity, may contribute to the inflammatory response. Viral genetic material activates TLR7, and there have been several reports linking TLR7 deficiency with severe outcomes in COVID19 (16; 69; 70). Less well understood is the role that S-protein-induced inflammation and cell injury may play in this disease. There is increasing evidence that s-protein can activate innate immunity, including toll-like receptors TLR2 and TLR4 (2; 36; 58; 72) Thus, it is important to understand the inflammatory effect of S1 on cells, independent of viral replication, if we are to grasp the full spectrum of SARSCoV2 effects on mammalian biology.

Hyaluronan or hyaluronic acid (HA) is a linear polymer formed by a repeating disaccharide structure of glucuronic acid and N-acetyl-glucosamine consisting of up to 25,000 disaccharide units (MW=1-10 million Daltons). Dysregulation of HA expression is relevant in disease and injury, (32). High Molecular Weight Hyaluronan (HMW-HA) is a major structural component of the extracellular matrix; it promotes cell survival, has anti-angiogenic properties and anti-inflammatory effects on immune cells, some of these mediated via binding to its receptors CD44 TLR2, and TLR4 (17; 20; 31; 56). LMW-HA (L-HA ∼300 kDa) fragments, act as endogenous innate immune ligands, and promote inflammatory responses, angiogenesis, and epithelial to mesenchymal transition (20; 31; 32; 42). LMW-HA increases vascular permeability by activating RhoA and ROCK (its downstream kinase), inducing cytoskeletal reorganization, and inhibiting cell-cell contacts (39; 41). LMW-HA is increased in severe COVID19 (24; 37; 53). On the contrary, administration of HMW-HA protects from the development of lung injury in ozone exposure (19), bleomycin administration (30), smoke inhalation, and sepsis (27; 45) and has been used in the treatment of COVID19 (clinicaltrials.gov, NCT04830020). Thus, HMWHA can be a useful anti-inflammatory agent in SARS-CoV2-induced cell injury and inflammation. A pharmaceutical preparation of HMW-HA (Yabro^®^; ∼1,000 kDa) has been shown to prevent exercise-induced bronchoconstriction in patients with asthma (52). No protective effects were seen following administration of LMW-HA (61). Several studies indicate that the relevant factor for HMW-HA protection is their size and not the source from which HMW-HA is derived (27; 31; 45). Thus, HMWHA can be a useful anti-inflammatory agent in SARS-CoV2-induced cell injury and inflammation.

To better understand both organ specific, and global effects of COVID-19, in addition to better drug targets, a growing number of bioinformatics studies include the mining of public data repositories containing –omics data sets derived from COVID-19 patients and translational models (28; 49).

Herein we used an unsupervised global proteomics analysis followed by systems biology analysis to identify changes in the proteome of an ATII-like Rat lung epithelial cell line (L2) following incubation with SARS-Cov-2 Spike S1 recombinant protein independent of the S-protein-ACE2 interaction, since rat ACE2 has very low affinity to S1. Our findings showed the S1 treatment alone resulted in significant modification to the Rat L2 cellular proteome. Network analysis indicated increased activation of E2F1, CREB1 and RhoA/ROCK2 networks, which were confirmed by Western blot studies. Conversely, pre-incubation of L2 cells with HMW-HA prior to addition of S1 prevented these changes. Previous protein-protein interaction studies along with analysis of differentially expressed genes identified within RNA sequencing datasets from COVID-19 patients highlighted the transcription factor E2F1 as an important player(57).

## MATERIALS AND METHODS

### Materials

SARS-Cov-2 (2019-nCoV) Spike S1-His recombinant protein (Cat. No 40591-V08H) was purchased from Sino Biological Inc. Yabro [pure, pharmaceutical-grade HMWHA, MW, 800,000 –1,200,000, at a concentration of 3 mg/ml was provided by IBSA Farmaceutici Italia srl, Italy. The chemical reagents dithiothreitol (cat. no. D9779) and iodoacetamide (cat. no. I1149) were purchased from Sigma-Aldrich (St. Louis, MO). Acetonitrile (ACN) was purchased from Thermo Fisher Scientific (cat. no. A996SK; St. Louis, MO). All other disposables are referenced within the manuscript.

### Cell culture and exposure to spike S1 protein

Rat lung type II-like epithelial cells (L2) were provided to us by Dr. Judith Creighton, previously at the University of Alabama at Birmingham. Their origin was characterized and verified using the presence of human and rat genes using PCR, immunohistochemistry, and Western blots of appropriate markers (reference (5); supplementary figure S1). Cells were cultured in DMEM media supplemented with 10% FBS, and 1% antibiotics in a humidified incubator with 95% air and 5% CO2 at 37°C until they became confluent as determined by light microscopy examination. For the experiment, cells were seeded in six-well plates at 5×104 cells·cm^2^ and incubated at 37°C until they became confluent in a humidified atmosphere of 95% air and 5% CO2. On the day of the experiment, some wells were pretreated with HMW-HA (300 µg/ml) for 1 h, and after that cells were treated with S1 protein (100 ng total) for either 24 h, at which time they were used for various exeriments.

### Proteomics Analysis

Proteomics analysis was carried out as previously described (3). The protein fractions were quantified using an 8 point Pierce BCA Protein Assay Kit (Thermo Fisher Scientific, Cat.# PI23225), and 20 µg of protein per sample diluted to 35µL using NuPAGE LDS sample buffer (1x final conc., Invitrogen, Cat.#NP0007). Proteins were then reduced with Dichlorodiphenyltrichloroethane (DTT) and denatured at 70°C for 10 min prior to loading onto Novex NuPAGE 10% Bis-Tris Protein gels (Invitrogen, Cat.# NP0315BOX) and separated half way (15min at 200 constant voltage). The gels were stained overnight with Novex Colloidal Blue Staining kit (Invitrogen, Cat.# LC6025). Following de-staining, each lane was partitioned into three separate MW fractions and equilibrated in 100mM ammonium bicarbonate (Millipore SIGMA; CAS # 1066-33-7). Each gel plug was then digested overnight with Trypsin Gold, Mass Spectrometry Grade (Promega, Cat.# V5280) following manufacturer’s instruction. Peptide extracts were reconstituted in 0.1% Formic Acid/ and double distilled H_2_O at 0.1µg/µL.

### nLC-ESI-MS2 Analysis & Database Searches

Peptide digests (8µL each) were injected onto a 1260 Infinity nHPLC stack (Agilent Technologies), and separated using a 100 micron I.D. x 13.5 cm pulled tip C-18 column (Jupiter C-18 300 Å, 5 micron, Phenomenex). This system runs in-line with a Thermo Orbitrap Velos Pro hybrid mass spectrometer, equipped with a nano-electrospray source (Thermo Fisher Scientific), and all data were collected in collision-induced dissociation (CID) mode. The nHPLC was configured with binary mobile phases that included solvent A (0.1%FA in ddH2O), and solvent B (0.1%FA in 15% ddH2O / 85% ACN), programmed as follows; 10min @ 5%B (2µL/ min, load), 90min @ 5%-40%B (linear: 0.5nL/ min, analyze), 5min @ 70%B (2µL/ min, wash), 10min @ 0%B (2µL/ min, equilibrate). Following each parent ion scan (300-1200m/z @ 60k resolution), fragmentation data (MS2) was collected on the top most intense 15 ions. For data dependent scans, charge state screening and dynamic exclusion were enabled with a repeat count of 2, repeat duration of 30s, and exclusion duration of 90s.

The XCalibur RAW files were collected in profile mode, centroided and converted to MzXML using ReAdW v. 3.5.1. The data was searched using SEQUEST, which was set for two maximum missed cleavages, a precursor mass window of 20ppm, trypsin digestion, variable modification C @ 57.0293, and M @ 15.9949. Searches were performed separately with both Human and Rat species-specific subsets of the UniProtKB databases. Following this analysis, we utilized the high-confidence *rattus norvegicus* to human tryptic peptide homolog data to avoid confusion for down-stream Systems Analysis.

### Peptide Filtering, Grouping, and Quantification

The list of peptide IDs generated based on SEQUEST (Thermo Fisher Scientific) search results were filtered using Scaffold (Protein Sciences, Portland Oregon). Scaffold filters and groups all peptides to generate and retain only high confidence IDs while also generating normalized spectral counts (N-SC’s) across all samples for the purpose of relative quantification. The filter cut-off values were set with minimum peptide length of >five amino acids (AA), with no (*m/z*+1) charge states, with peptide probabilities of >80% confidence intervals (C.I.), and with the number of peptides per protein ≥2. The protein probabilities were then set to a >99.0% C.I., and a Dlaw False Discovery Rate (FDR)<1.0. Scaffold incorporates the two most common methods for statistical validation of large proteome datasets, the false discovery rate (FDR) and protein probability (35; 50; 64). Relative quantification across experiments were then performed via spectral counting(44), and when relevant, spectral count abundances were then normalized between samples (10).

### Statistical & Systems Biology Analysis

For the proteomic data generated, two separate non-parametric statistical analyses were performed between each pair-wise comparison. These non-parametric analyses include 1) the calculation of weight values by significance analysis of microarray (SAM; cut off >|0.6|combined with 2) t-test (single tail, unequal variance, cut off of p < 0.05), which then were sorted according to the highest statistical relevance in each comparison. For SAM (22) whereby the weight value (W) is a statistically derived function that approaches significance as the distance between the means (μ1-μ2) for each group increases, and the SD (δ1-δ2) decreases using the formula, W=(μ1-μ2)/(δ1-δ2). For protein abundance ratios determined with N-SC’s, we set a 1.5-2.0 fold change as the threshold for significance, determined empirically by analyzing the inner-quartile data from the control experiment indicated above using ln-ln plots, where the Pierson’s correlation coefficient (R) was 0.98, and >99% of the normalized intensities fell between +/-1.5 fold. In each case, any two of the three tests (SAM, t-test, or fold change) had to pass.

Gene ontology assignments and pathway analysis were carried out using MetaCore (GeneGO Inc., St. Joseph, MI). Interactions identified within MetaCore are manually correlated using full text articles. Detailed algorithms have been described previously(11)

### Measurement of RhoA activity

The Rho activity was measured and quantified by using the RhoA Activation Assay Biochem Kit (cat. no. BK036; Cytoskeleton) based on the Rhotekin pull-down assay as per the manufacturer’s instructions as described previously(39). In brief, L2 cells (*n* =10) were cultured in 6 well culture plates and few wells were pretreated with HMW-HA 300 µg/ml for 1 h and then treated with or without spike S1 (200 ng/ml) protein for 30 min at 37 °C. The cell lysate was prepared in radioimmunoprecipitation assay (RIPA) buffer containing protease inhibitors (cat. no. 78425; Thermo Fisher Scientific) immediately after treatment and protein estimation was done, and equal amounts of protein were incubated with Rhotekin-RBD beads (cat. no. RT02) for 1 h at 4°C. After the beads were washed with wash buffer, proteins were removed from the beads with Laemmli buffer and then subjected to Western blotting.

### Measurement of E2F1 and ROCK2 phosphorylation

In brief, L2 cells (*n* =10) were cultured in six well culture plates and few wells were pretreated with HMW-HA (300 µg/ml) for 1 h and then treated with or without spike S1 (200 ng/ml) protein for 24 h at 37 °C. The cell lysates were prepared in RIPA buffer containing protease inhibitors (Thermo Fisher Scientific, Rockford, IL) immediately after treatment. Protein estimation was carried out using the BCA assay. Equal amounts of protein in Laemmli buffer were loaded in 4-20% gradient Tris·HCl Criterion precast gels, and proteins were transferred to PVDF membranes. Membranes were probed with p-E2F1 (1:1,000 dilution; cat. no. MABE1782; MilliporeSigma) or E2F1 (1:1,000 dilution; cat. no. 3742S; Cell Signaling Technology, Beverly, MA) and ROCK2 (1:1,000 dilution; cat. no. 8236; Cell Signaling Technology, Beverly, MA) or pTyr722-ROCK2 antibody (cat. no. SAB4301564; MilliporeSigma). Bands were detected by the chemiluminescent horseradish peroxidase substrate. Protein loading was normalized by reprobing the membranes with an antibody specific to β-actin.

### Cell Cycle Analysis by Flow Cytometry

Analysis of Cell Cycle: Cell cycle was analyzed using staining with DAPI nucleic acid stain (Thermofisher, Waltham, MA) in cells following dissociation and fixation in single cell suspension. DAPi fluorescence was detected using flow cytometry in the UAB Comprehensive Flow Cytomery Core facility using an LSR II Flow Cytometer (Beckton Dickinson, Franklin lakes, NJ) and cell cycle was determined based nucleic acid content/cell. Cells associated with the peak at the lower fluorescence intensity were deemed to be G0/G1 cells, cells in the peak at the higher fluorescence were deemed G2M phase, and cells with intermediate fluorescence were determined to be S phase.

### Statistical Analysis for Non-Proteomics Studies

Figures were generated and statistics performed using GraphPad Prism version 8 for Windows (GraphPad Software, San Diego, CA). The mean ± SEM was calculated in all experiments, and statistical significance was determined by either the one-way or the two-way ANOVA. For one-way ANOVA analyses, Tukey’s multiple comparisons test was employed, while for two-way ANOVA analyses, Bonferroni posttests were used. Overall survival was analyzed by the Kaplan–Meier method. Differences in survival were tested for statistical significance by the log-rank test. A value of p < 0.05 was considered significant.

### Pathway Associations with Human Cell Lines Exposed to SARS-CoV-19

To determine whether the associated functions observed in the S1-Tx proteomics L2 cells are correlated with the transcriptomic data, the RNA-seq transcriptomic data of five human lung cell lines that were exposed to SARS-CoV-2 were downloaded from public domain (**Supplementary Table 6**). The Gene Set Enrichment Analysis (GSEA v4.0.3; https://www.gsea-msigdb.org/gsea/index.jsp) was conducted against the Hallmark and C2 curated canonical pathways datasets in the Molecular Signatures Database (MSigDB v7.4) with 1000 permutations (min size 15, max size 500). Gene sets with adjusted FDR q-value less than 0.25 were considered as significant.

## RESULTS

### Spike S1 Effects on the ATII Like Rat L2 Proteome

We tested whether treatment (Tx) of ATII-like Rat L2 cells with SARS-Cov-2 Spike S1-recombinant protein for 24hrs causes significant changes to its proteome (Tx-S1 vs. vehicle control/ C, n=4). The proteomics analysis revealed 1678 altered aproteins, seven hundred thirty of which had <1% FDR with at least 3 out of 4 replicates measuring non-zero values across all groups tested (**Supplementary Table 1**). For the pairwise comparison between Tx-S1 vs. C, there were 99 proteins found to be significantly changed in abundance, fifty-one ow which increased (**Table 1**), and fourty eight decreased (**Table 2**). This data set was best visualized using a Volcano Plot (**Figure 1**), in addition to a two dimensional (2D) hierarchical heat map (HCA-HM, **Figure 2a**) followed by principal component analysis (PCA, **Figure 2b**). All statistical analysis and visualizations were carried out using the data matrix found in **Supplementary Table 1**.

**Figure 1.**
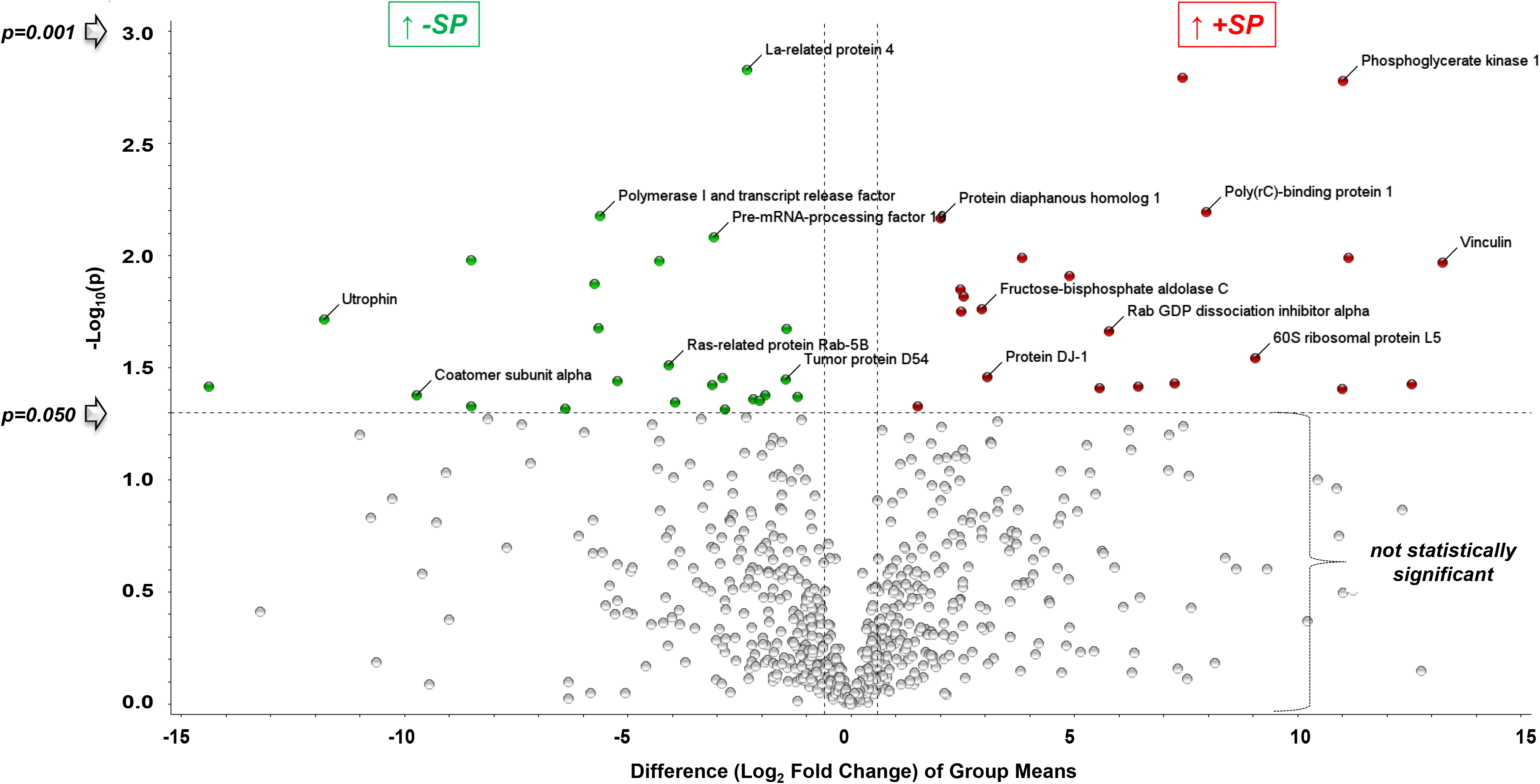
Volcano Plot of Proteins Increased or Decreased in Abundance following addition of S1 vs. C (Rat L2 Cells; S1-Tx vs. C, 24hrs). In this plot, the y axis is the negative log(base 10) of the p value. This results in highly significant points with low p values appearing toward the top of the plot. The log of the fold change (FC) between the two conditions is plotted on the x axis in base 2 (ex: Log2 Fold 35=5). The log fold change is adapted so that changes in both directions become equidistant from the center. In this way, the points in the plot form two regions of interest with significant changes: those points that are found toward the top of the plot that are either on the left- or right-hand sides. These represent values that display large magnitude fold changes as well as high statistical significance. The log2(fold-change) is the log-ratio of a protein’s expression values under pairwise conditions. While comparing two conditions each feature analyzed gets (normalized) expression values.

**Figure 2.**
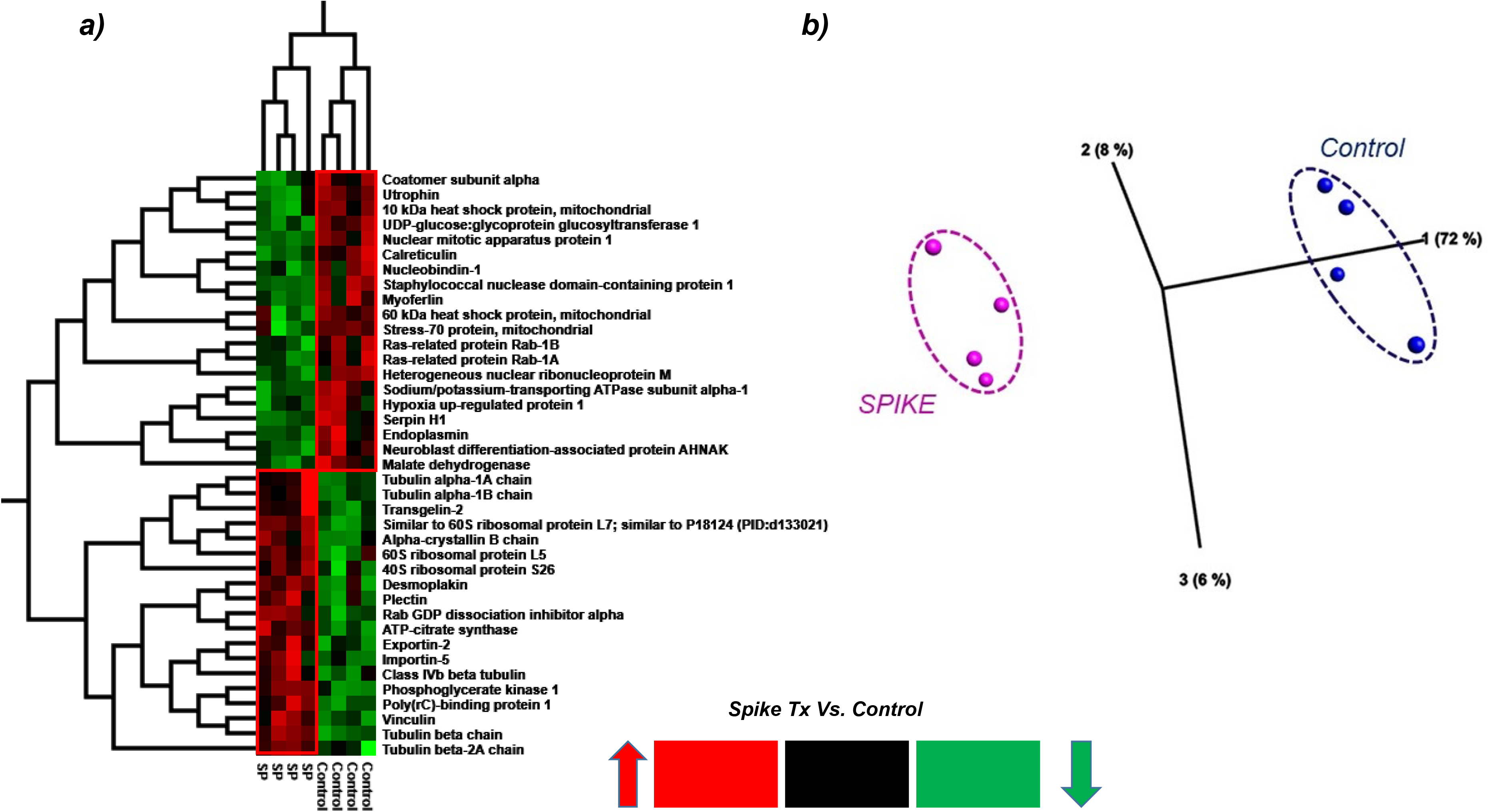
a) 2D HCA Heat Map and b) PCA Plot of the 36 most significantly changed proteins from Rat L2 Cells S1-Tx vs C for 24hrs. Of note, the following 8 proteins are all highly significant, stemming from SupTable1, but do not quite meet the +/− 1.5 fold change required for Tables 1-2: Myoferlin↓, Stress-70 Protein↓, Sodium/ Potassium transprting atp↓, Tubulin alpha 1A↑, Tubulin alpha 1B↑, Importan-5↑, Class Ivb beta tubulin↑, Tubulin beta chain↑. Each column in the HCA Heat Map and each circle in the PCA plot represent different experiments.

**Table 1.**
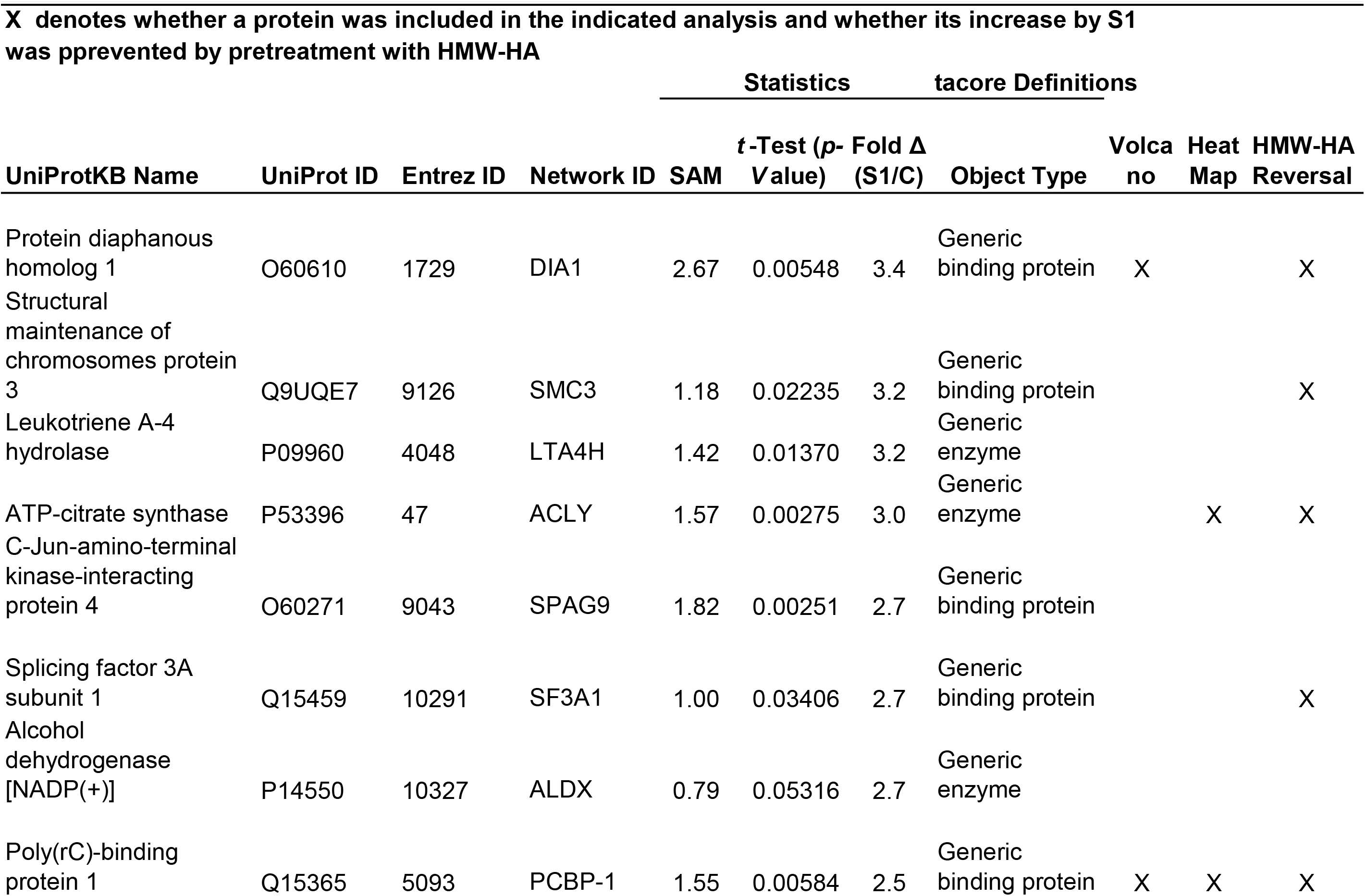
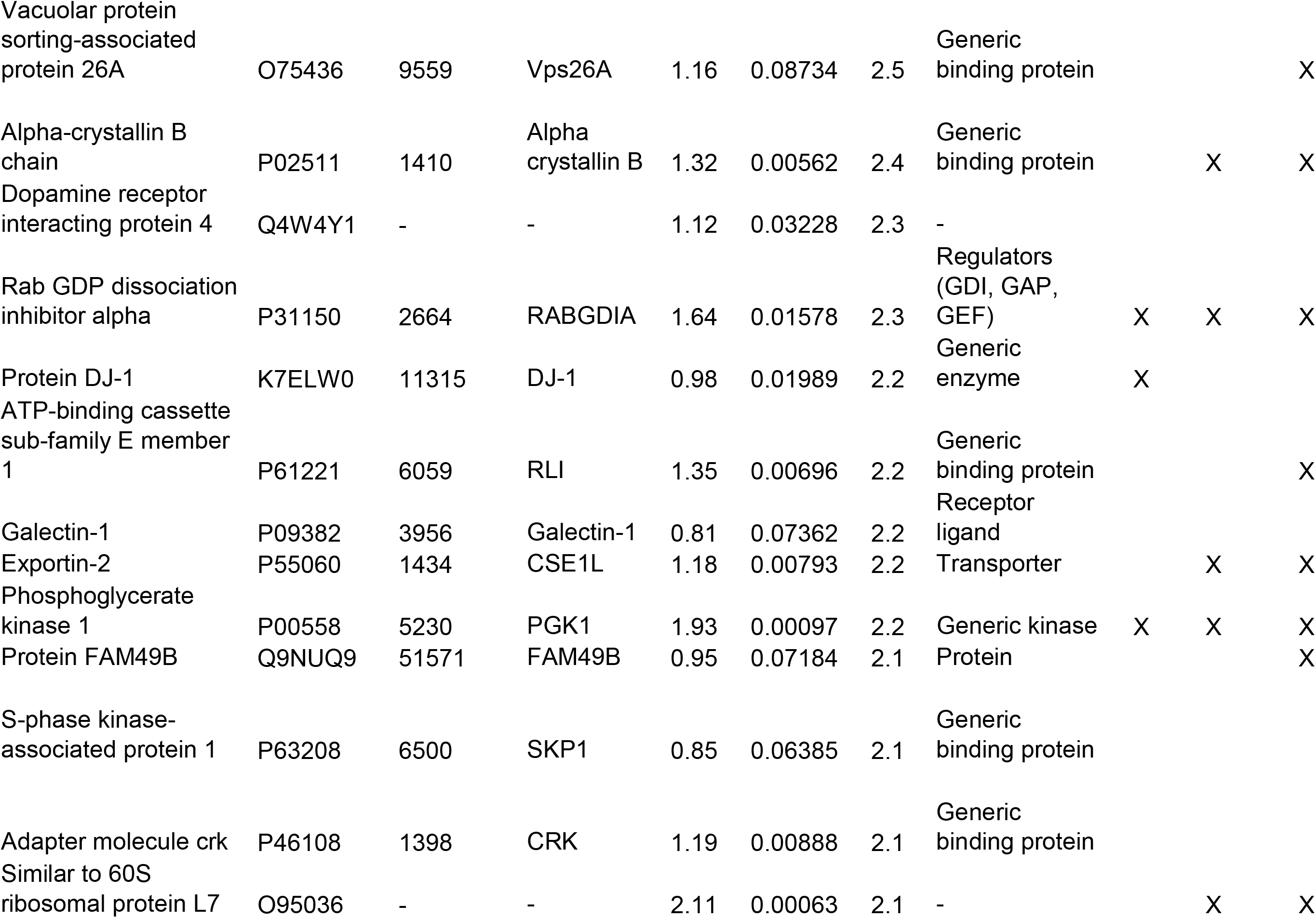
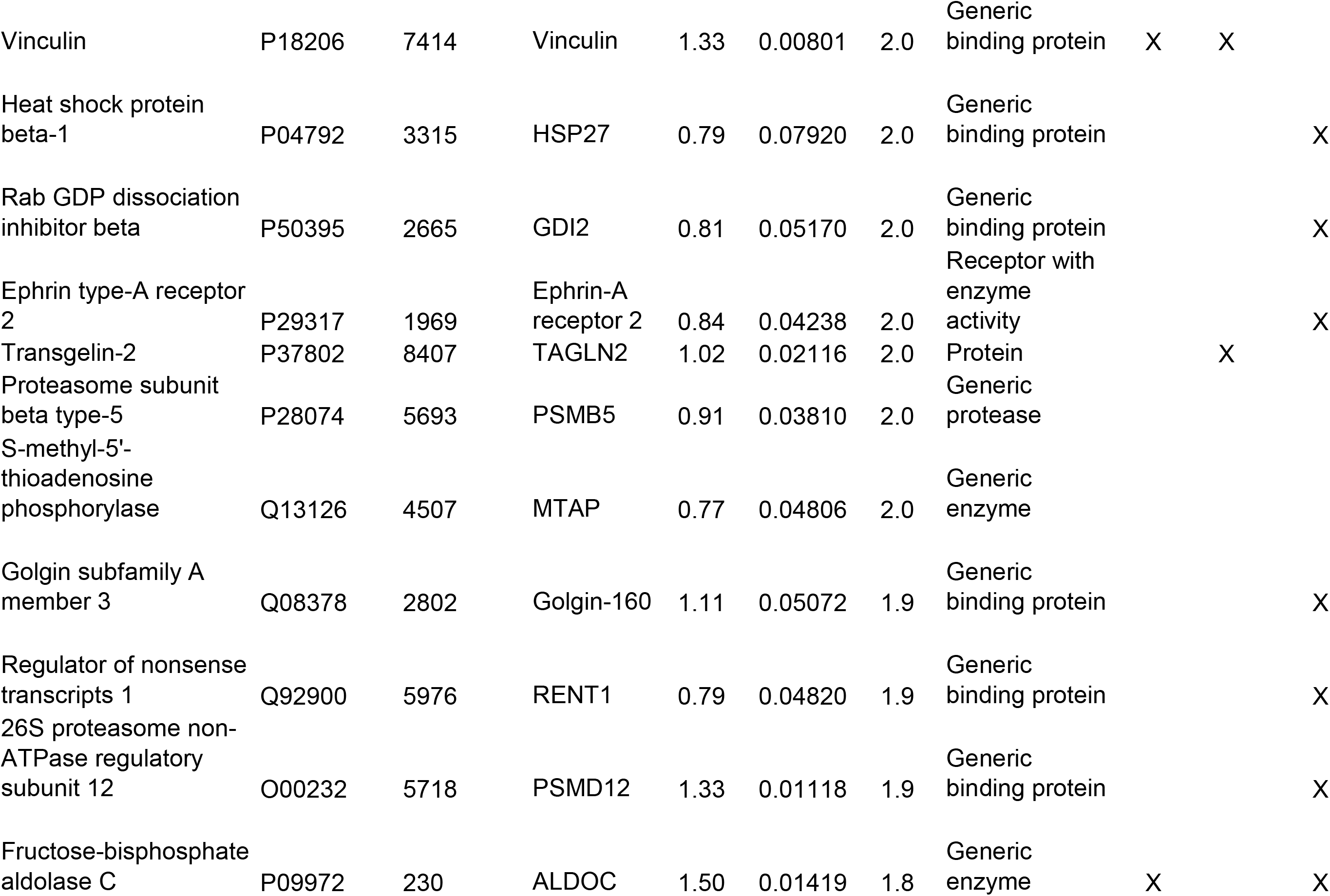
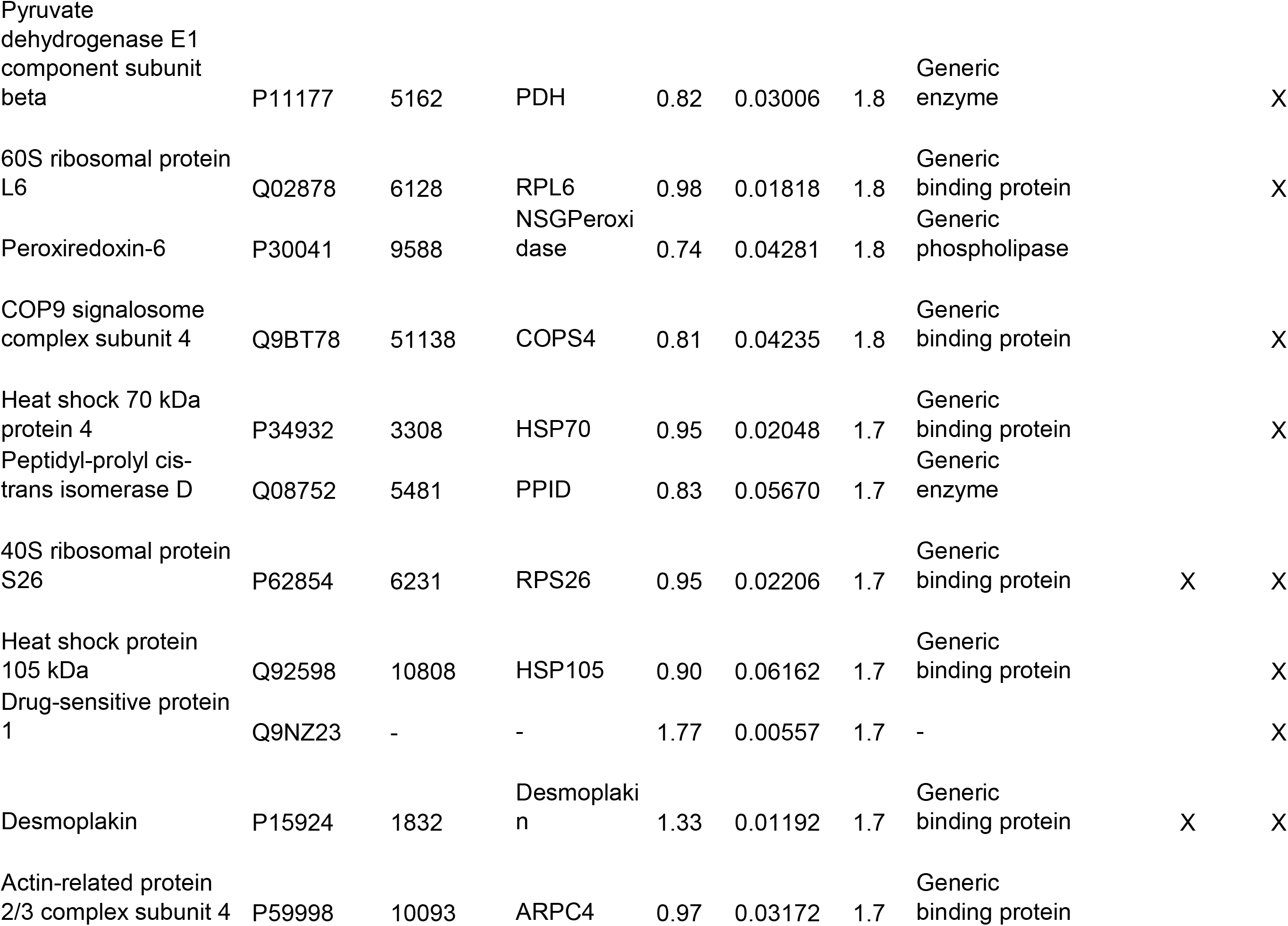
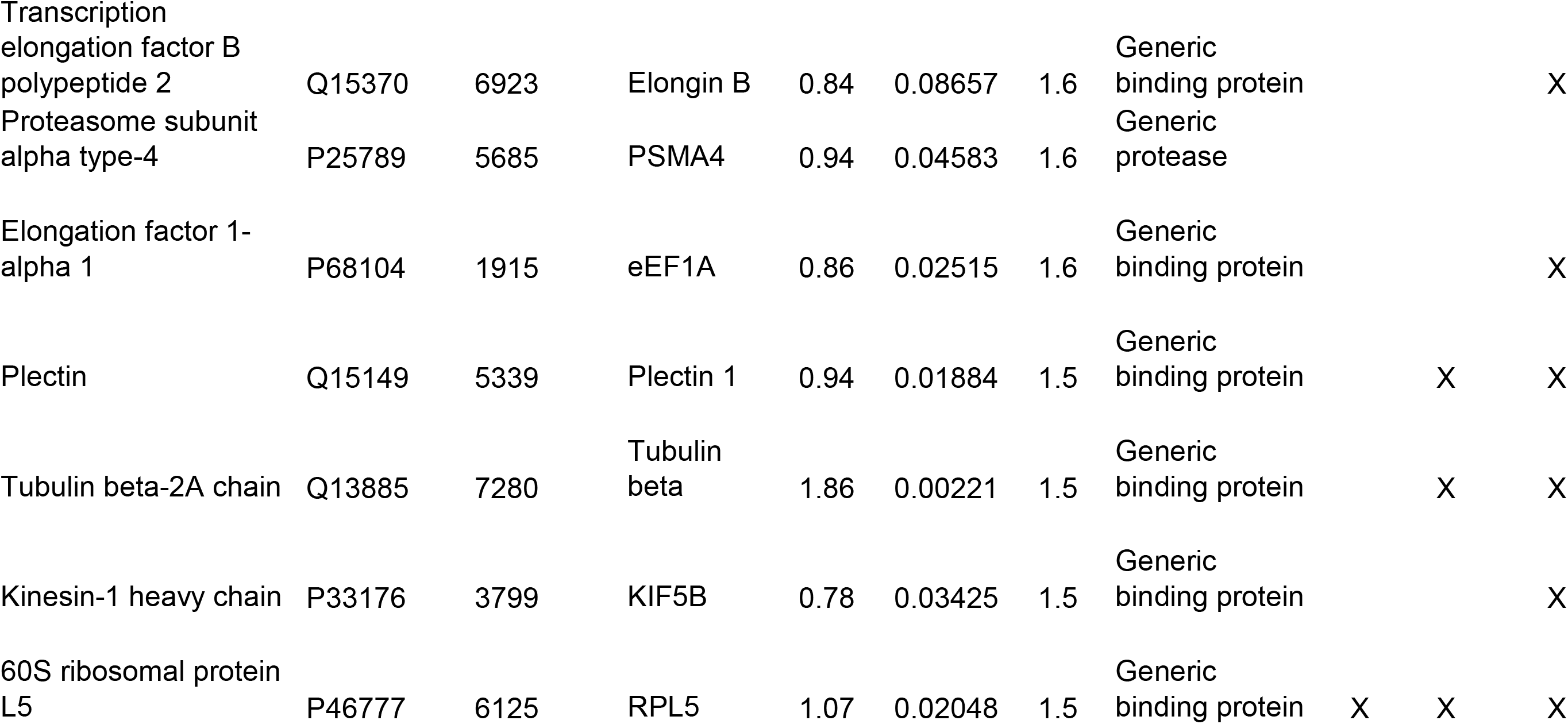
List of fifty-one proteins of rat L2 cells that were increased signficantly twenty-four h post S1.

**Table 2.**
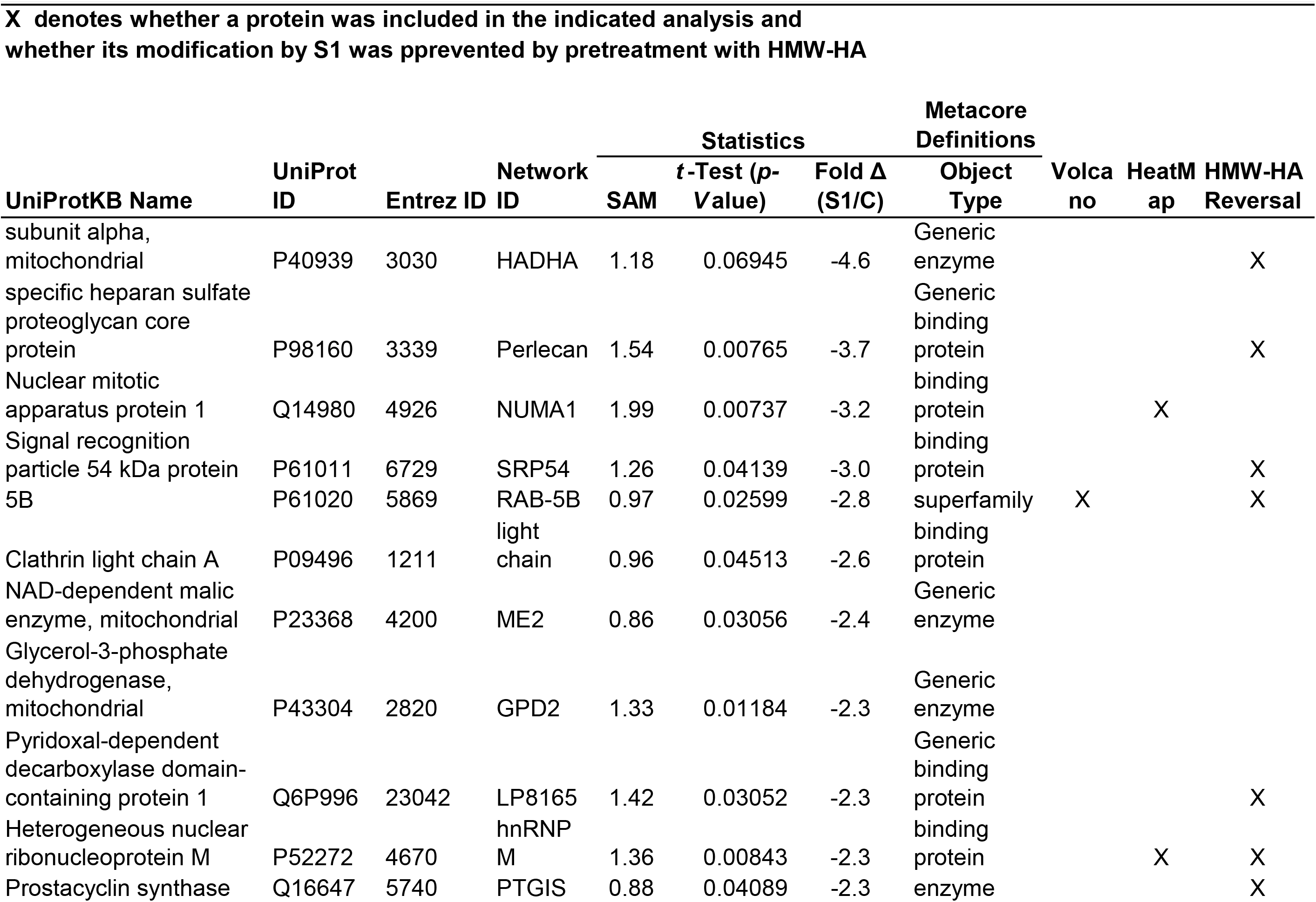
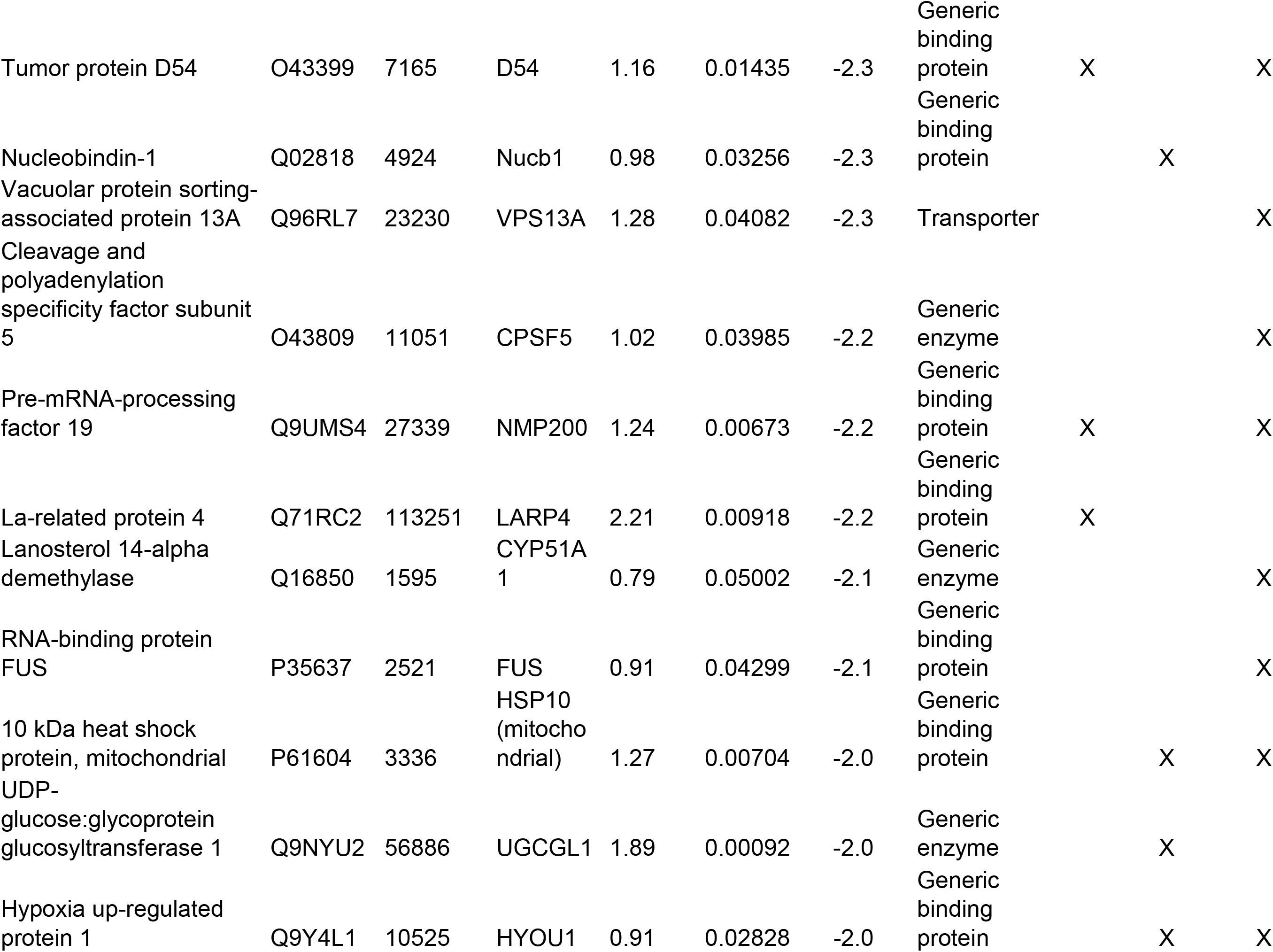
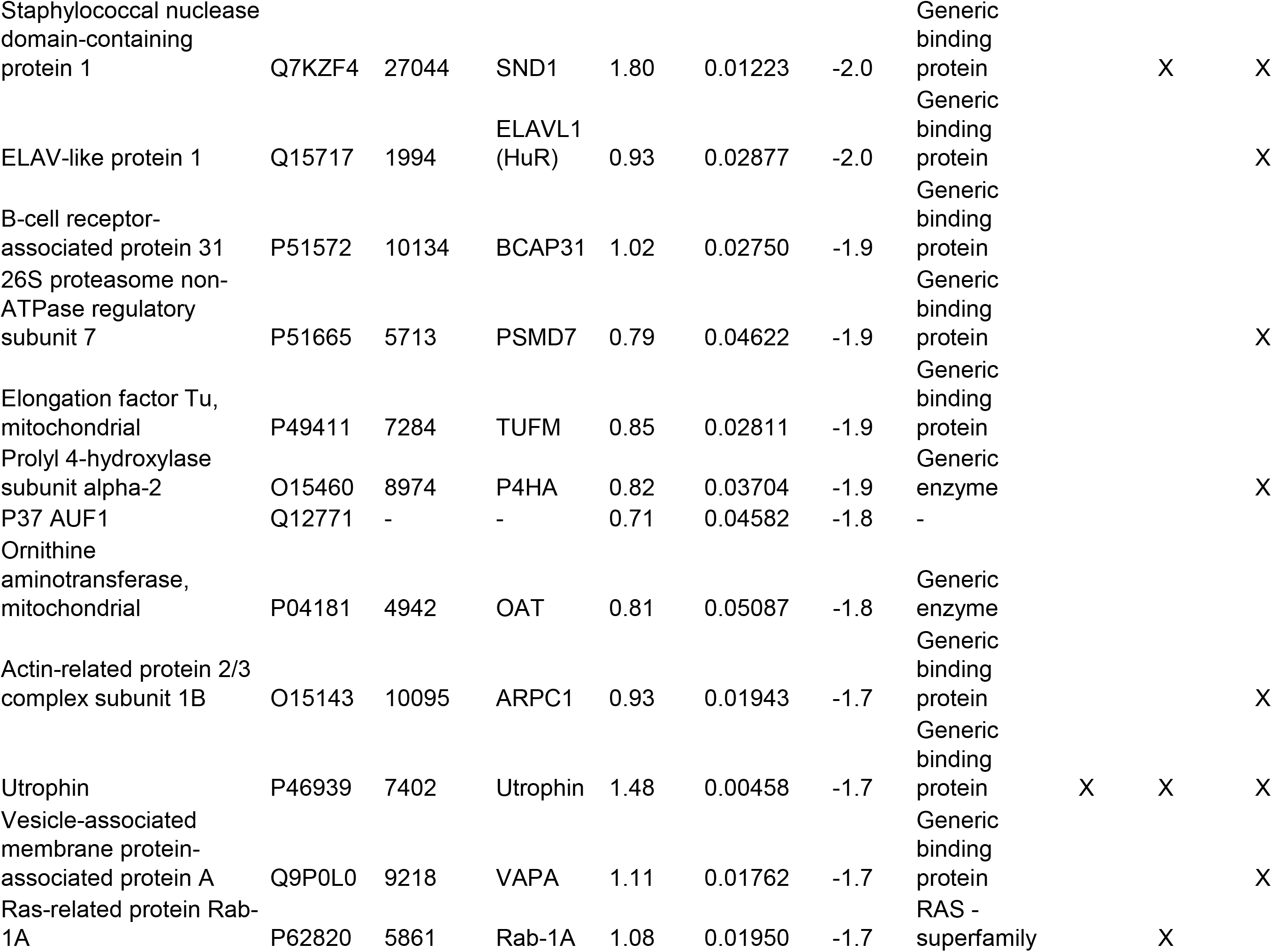
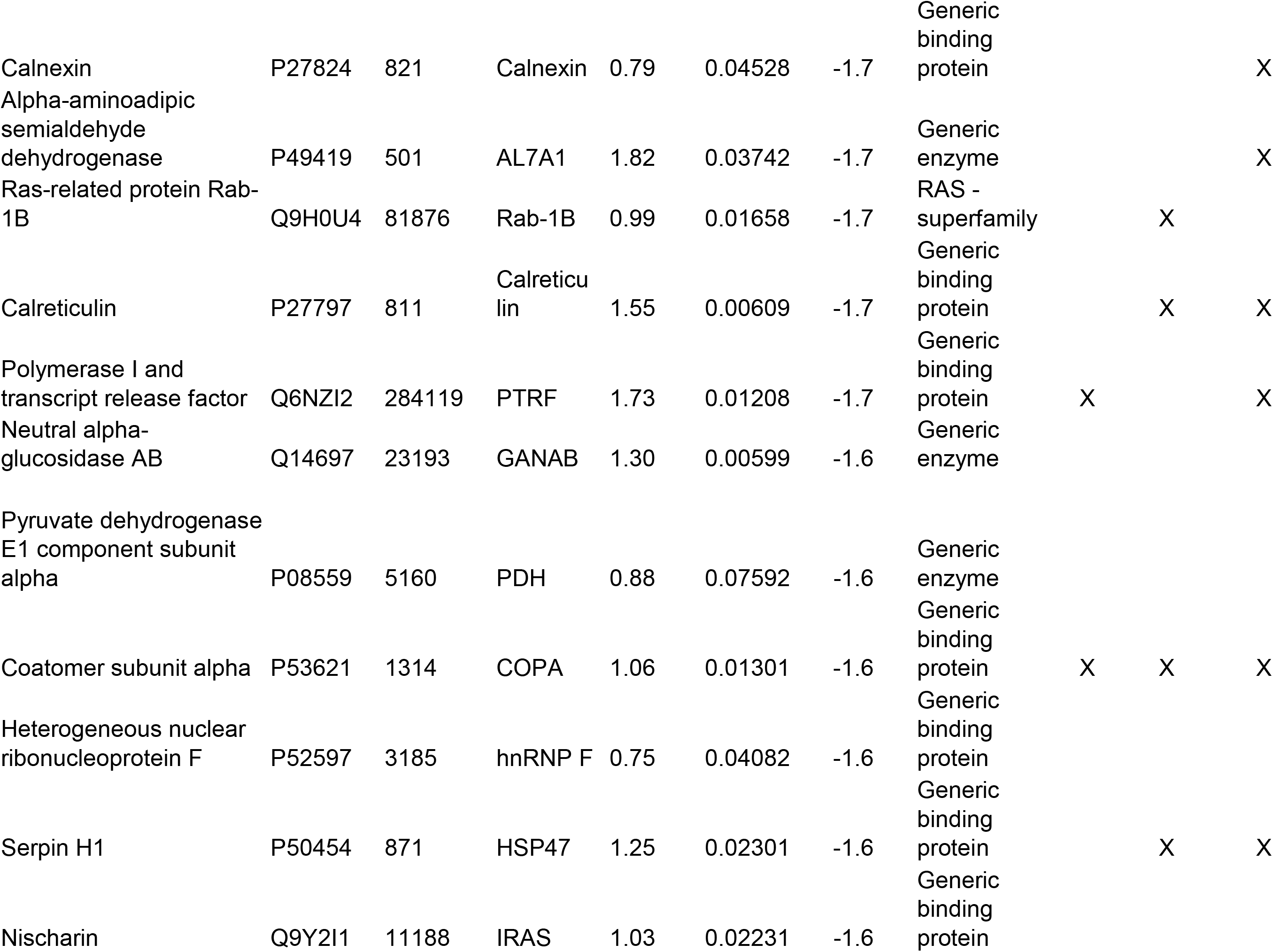
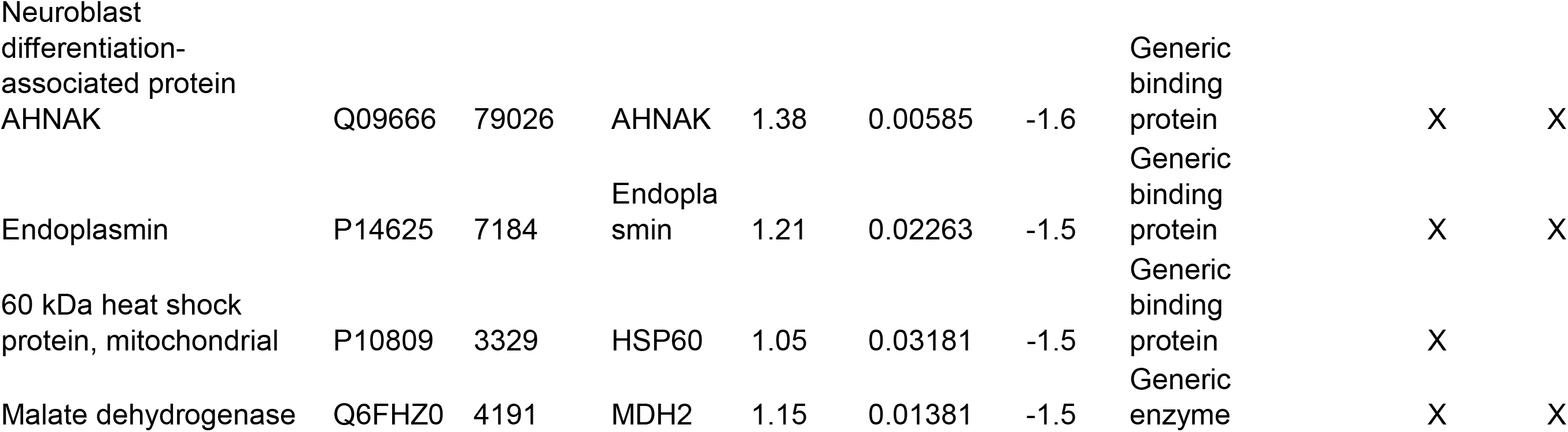
List of fourty eight proteins of Rat L2 cells that were signficantly decreased twenty four h post S1.

### Spike S1 Associated Biological Processes and Pathways via Systems Biology

Proteins that significantly changed, as listed in **Tables 1** and **2** for Tx-S1 of L2 cells, were combined and utilized for in-depth systems biology analysis, which allowed for the identification of key biological pathways associated solely with S1treatment. Following this analysis, we created a pie chart to better visualize the more common gene ontology (GO) based Process Networks that the most changed proteins are associated with (**Figure 3**). This analysis suggests that Tx-S1 has an effect on many cellular functions, including: catabolic and glucose metabolism, ER-stress, cell migration, and signal transduction. We then performed unbiased network analysis using Metacore’s Systems Biology software https://www.nihlibrary.nih.gov/about-us/news/metacore-enabling-systems-biology-research-through-pathway-analysis. This analysis identified three important network hubs that cross correlate with the E2F1 pathway (**Figure 4**), ROCK2/ RhoA (**Supplementary Table 2**), and CREB/ RelA (**Supplementary Figure 2**, and **Table 3**). To better understand the Network and Pathway figures, we have included the figure legend (GenegoMetacore, **Supplementary Figure 1**). As illustrated in **Figure 4** (red circles increased, blue circles decreased in S1-Tx vs. C L2 cell) nearly the entire list of significantly changed proteins after S1 addition (listed as network IDs in **Tables 1** and **2**) are associated with E2F1 signaling. Similarly in **Supplementary Figure 2**, both CREB1 and RelA transcription factors are highlighted as central network hubs. Further analysis illustrated in **Supplementary Table 2** highlights networks such as cell adhesion, cytoskeleton associated (actin filaments, rearrangement, spindle and cytoplasmic tubules), in addition to cell cycle mitosis and meiosis. Also of interest, there was an apparent pronounced effect on cellular MHC class I Immune response as the top pathway (**Supplementary Figure 3**).

**Figure 3.**
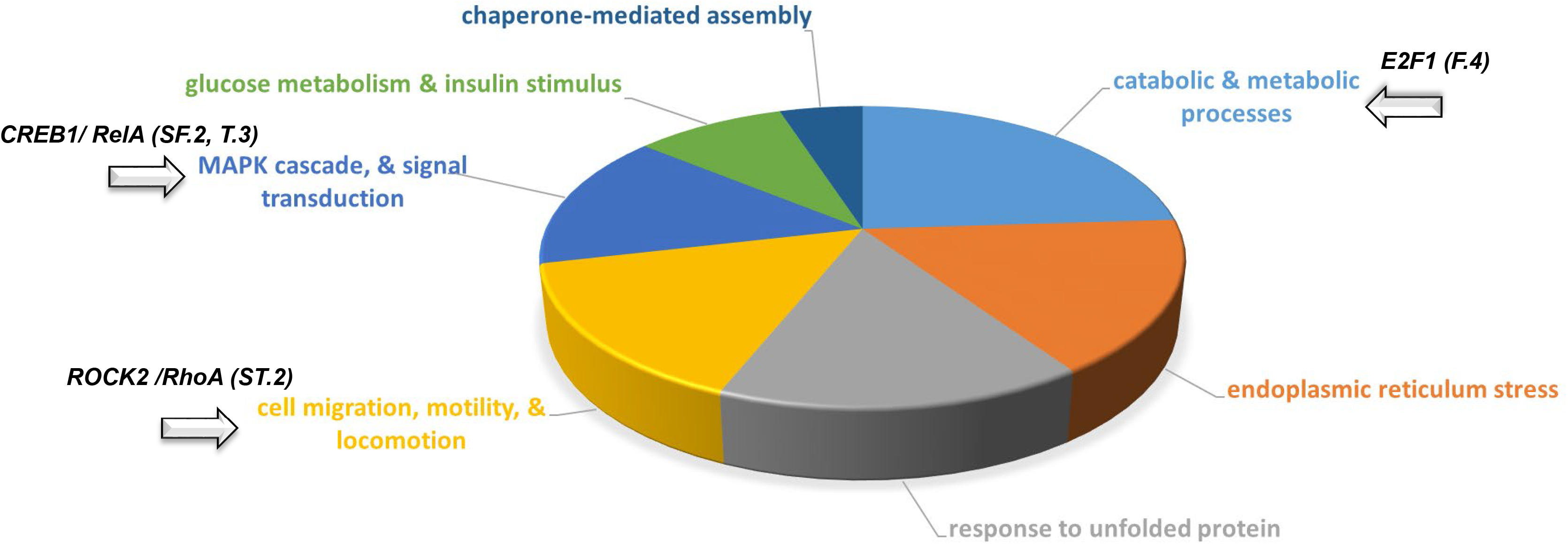
Top Process Networks Pie Chart from Systems Biology Analysis carried out on the most significantly changed proteins (Tables 1-2), from Rat L2 Cells were treated with SARS CoV-2 S1 (S1-Tx) or vehicle © for 24hrs. Highlighted segments contain proteins associated with (1) ROCK22/ RhoA (cell migration, motility, and locomotion; SuppTable. 2), CREB1/ RelA (MAPK cascade, and signal transduction; SuppFigure. 2 and Table. 3), and E2F1 (catabolic and metabolic processes; Figure. 4).

**Figure 4.**
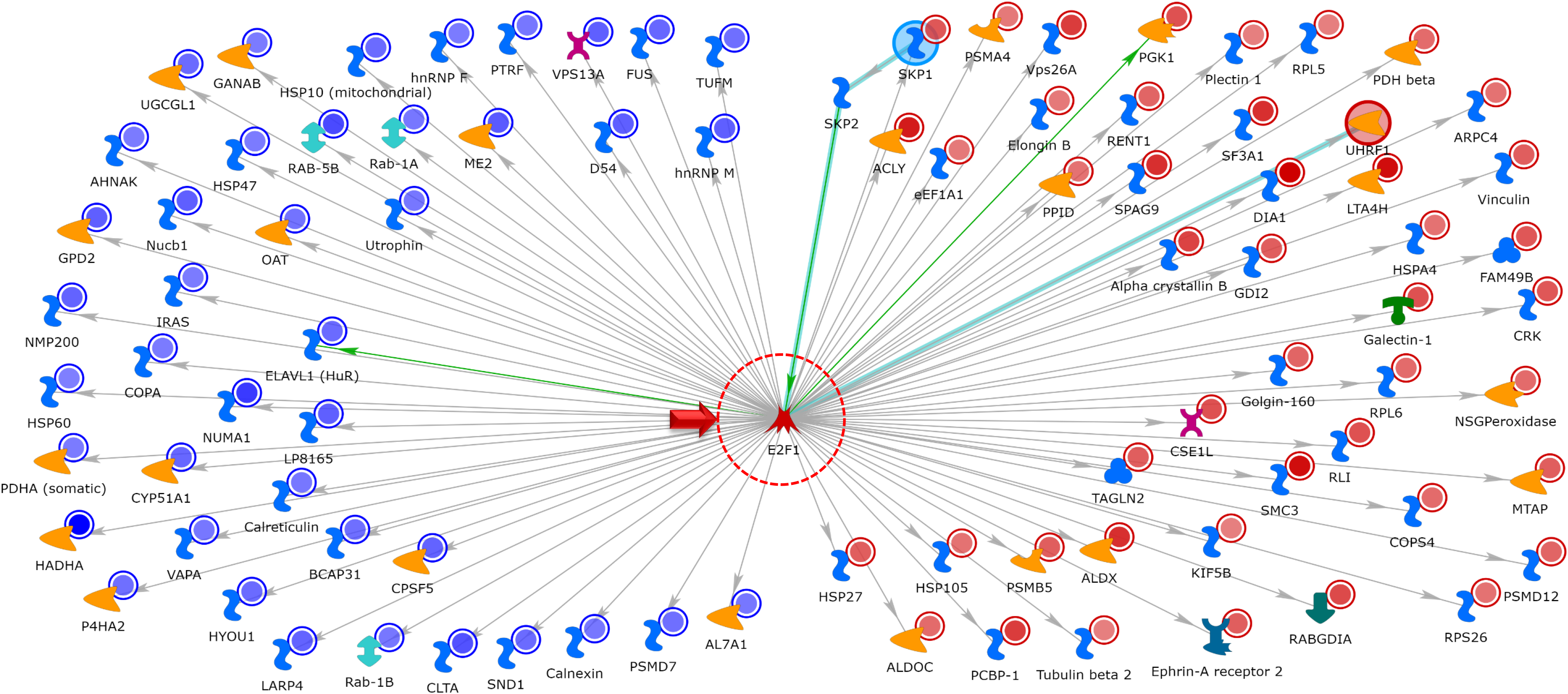
Top Network Map (S1-Tx vs C); with the use of the 99 most significant proteins identified in the Tables 1-2, using Metacore’s Systems Biology software to carry out Network Analysis. E2F1 was identified as a major centralized network hub. Blue circles indicate a decrease, while red circles indicate an increase in protein abundance (Rat L2 Cells; S1-Tx vs. C, 24hrs). Nearly the entire list of proteins identified were shown to interact/ associate in some way to E2F1 as a direct results of treatment of L2 cells with SARS CoV-2 S1 protein for 24 h (S1-Tx).

**Table 3.** Rat L2 Proteomic S1-Tx Cross-Correlated with Pathway Associations using Trancriptomic Data From Five Human Cell Lines Exposed to SARS-CoV-2 Virus.

| Pathway Accociations with Five Human Lung Derived Celll Lines Exposed to SARS-CoV-2 |  |  |  |  |  |
| --- | --- | --- | --- | --- | --- |
| GENESET | Pathway | #Targets | NES | p-value | q-value |
| **1. NHBE (Normal Human Bronchial Epithelial)** |  |  |  |  |  |
| HALLMARK | E2F | 195 | -1.4 | 0.0091 | 0.2044 |
| **2. A549 (adenocarcinoma human alveolar basal epithelial cells)** |  |  |  |  |  |
| HALLMARK | E2F | 197 | -3.1 | 0.0000 | 0.0000 |
| HALLMARK | G2M Checkpoint | 194 | -2.7 | 0.0000 | 0.0000 |
| HALLMARK | Fatty Acid Metabolism | 150 | -1.5 | 0.0105 | 0.0320 |
| **3. A549-ACE2** |  |  |  |  |  |
| HALLMARK | Fatty Acid Metabolism | 150 | -2.3 | 0.0000 | 0.0000 |
| HALLMARK | E2F | 196 | -1.8 | 0.0000 | 0.0002 |
| REACTOME | E2F Mediated DNA Replication | 22 | -1.9 | 0.0056 | 0.0051 |
| BIOCARTA | CREB | 22 | -1.6 | 0.0295 | 0/0661 |
| **4. HAE (Human Airway Epithelial)** |  |  |  |  |  |
| HALLMARK | Fatty Acid Metabolism | 151 | -1.6 | 0.0000 | 0.0177 |
| BIOCARTA | CREB | 21 | -1.7 | 0.0037 | 0.1270 |
| **5. Calu (non-small-cell lung cancer)** |  |  |  |  |  |
| HALLMARK | E2F | 197 | -3 | 0.0000 | 0.0000 |
| HALLMARK | G2M Checkpoint | 194 | -2.3 | 0.0000 | 0.0000 |
| HALLMARK | Fatty Acid Metabolism | 147 | -2.0 | 0.0000 | 0.0000 |
| REACTOME | E2F Mediated DNA Replication | 22 | -1.9 | 0.0030 | 0.0119 |
| KEGG | Cell Cycle | 124 | -1.6 | 0.0081 | 0.0510 |

### HMW-HA Pretreatment Attenuates Spike S1 Induced Changes to the Proteome of rat L2 cells

As shown in **Supplementary Figure 1**, and in **Supplementary Table 3**, incubation of L2 cells with HMW-HA had significant effects on their proteome. PCA analysis of the data from L2 cells treated with vehicle, S1 and preincubated with HMWA-HA prior to S1 (shown in **Supplementary Table 1)** showed tight clustereing and clear separation among groups. **Supplementary Figure 4**).

### HMW-HA Effects on the Rat L2 Proteome & Associated Pathways

In a separate follow up proteomics experiment, we found that the addition of HMW-HA to Rat L2 cells for 24 h caused significant changes in seventy-five proteins (thirty seven increased and thirty eight decreased, HMW-HA vs. C; **Supplementary Table 4**). A 2D-HCA Heat Map of the top most significantly changed 37 proteins (**Supplementary Figure 5a**) is illustrated along with a separate PCA plot (**Supplementary Figure 5b**).

Together these data indicate that HMW-HA induced a profound effect on the Rat L2 Proteome. The top 75 significantly differential protein list was then utilized for systems biology analysis. This analysis led to the identification of a number of significant Network Hubs and GO Biological Processes that include strong associations with E2F1, E2F5, and MCM-complex associated cellular response to stress, DNA Replication, and Cell Cycle Regulation (**Supplementary Table 5**). In addition, p53, NF-kB, and RAD54B associated protein SUMOylation and DNA Repair were also significantly affected by HMW-HA.

### S1 increased E2F1, RhoA and ROCK2 ROCK2 phosphorylation which were prevented by pre-incubation with HMW-HA

Western Blot Analysis carried out on L2 cells treated with S1 protein for 24 h resulted in a significant increase of E2F1 phosphorylation as compared to controls (**Figure 5a**). Pretreatment with HMW-HA prior to S1 addition prevented E2F1 phosphorylation when normalized to either actin or total E2F1 (**Figure 5a**). There were no changes in total E2F1 as compared to actin for any condition (data not shown). Similarly, and in agreement with the proteomics outcome, as illustrated in **Figure 5a**, incubation of L2 cells with the S1 protein induced phosphorylation (activation) of both RhoA (**Figure 5b**) and ROCK2 (**Figure 5c**) within 30min following incubation, whereas pretreatment with HMW-HA significantly inhibited these S1-associated effects.

**Figure 5.**
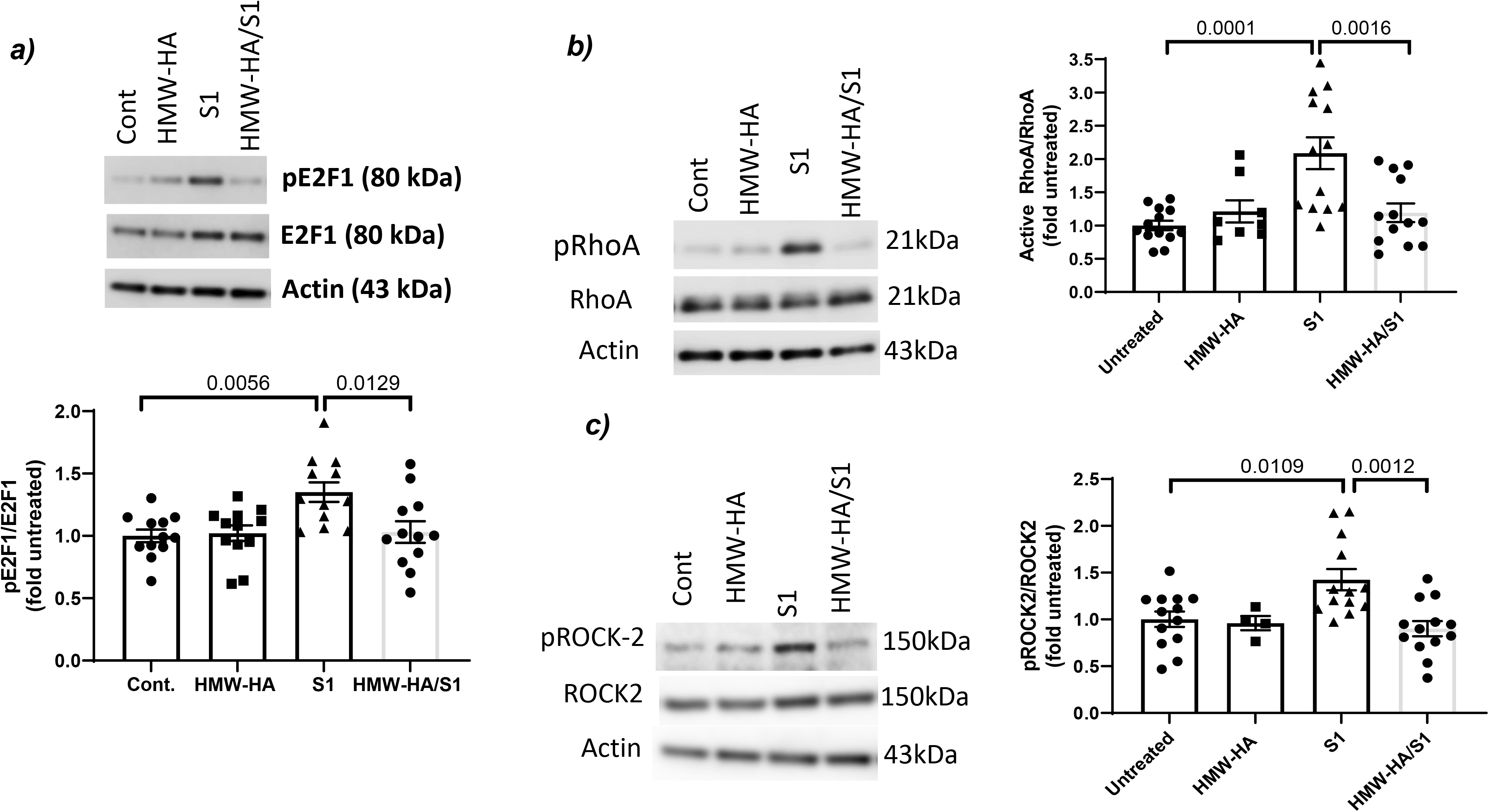
Western Blots of confluent Rat L2 cells, treated with PBS (Contr.), HMW-HA (300 µg/ml) for 24 h, S1 (100 ng) for 24 h or HMW-HA for 1 h followed by S1 for 24 h. **a**) Cells were immunostained with antiobodies against E2F1 or its phosphorylated for and actin (upper panel). The intensities of the shown bands was then quantified and the ratios of pE2F1/E2F1 is shown in the lower panel. (**b**,**c**) Cell were immunostained with antibodies against phosphRhoA (active RhoA) and total RhoA, phosphoROCK2 or total ROCK2 (left panels). The intensities of the various pands were quantified and the rations of the phosphorylated to total forms are shown in the right panels. Incubation of rat L2 cells with S1 upregulated the activities of the phosphorylated forms of E2F1, RhoA and its downstream kinase, ROCK2. Preincubation with HMW-HA prevented these effects. Values are means with Standard Error of the Means and individual points from three different batches of cells. One way analysis of variance followed by the Tukey’s t-test for multiple comparisons.

Since E2F1 plays an important role in cell cycle regulation, we assessed changes in cell cycle by S1 and its reversal by HMW-HA. As shown in **Figure 6**, treatment of L2 cells with the S1 protein for 24h resulted in a significant increase in S phase accumulation with a concomitant decrease in the G2 phase as compared to the non-treated control, and both of these changes were either reversed or blocked with 1h pretreatment with HMW-HA.

**Figure 6.**
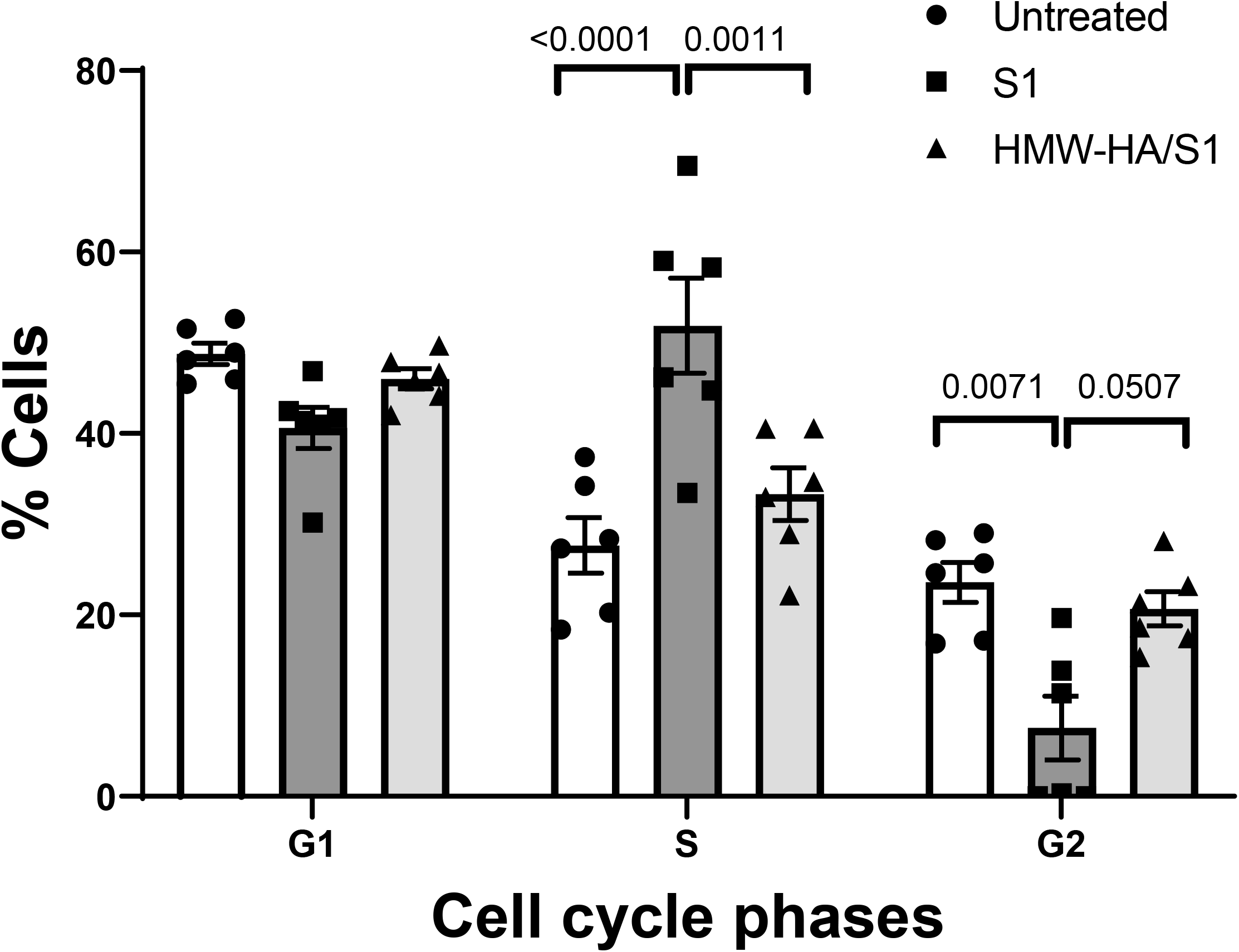
Cell cycle analysis of confluent Rat L2 cells incubated with S1 protein (100 ng) for 24 h or pretreated with HMW-HA (300 µg/ml) for 1h prior to incubation with S1 protein. Cell cycle analysis was conducted as described in the Materials and Methods. Values are means with Standard Error of the Means and individual points from two different batches of cells. One way analysis of variance followed by the Tukey’s t-test for multiple comparisons. Data from two different batches of cells.

### Bioinformatic Networks Correlation of Spike S1 Exposed Rat L2 Proteome Data to SARS-CoV-2 Exposed Human Cells Transcriptome

The list of 99 proteins stemming from statistically significant changes induced by 24h Tx-S1 in Rat L2 cells at the beginning of these studies (**Tables 1 & 2**) were strongly correlated with a number of pathwasys, including E2F1, Cell Cycle, Rock2/ RhoA, and CREB1. Following our validation, we were interested to see if there would be further correlation of these pathways at transcriptomic level using RNA-seq data from human lung cell lines exposed to live SARS-CoV-2. The Geneset Enrichment Analysis (GSEA) results revealed that E2F signaling, E2F Mediated DNA Metabolism, G2M Checkpoint, and/or Cell Cycle were significantly correlated in three of the cell lines, and CREB was also correlated in two of these lung derived human cell lines (**Table 3**)

## DISCUSSION

In this study we addressed two key questions; 1) can human SARS-CoV-2 Spike S1 protein elicit a global proteomic effect on a Rat lung derived ATII-like L2 cell line that is expected to be primarily rat-ACE2 independent when exposed to S1 protein(14), and 2) can HWW-HA attenuate these effects? Our findings support a novel proposed model of S-protein-induced cell injury (**Figure 7**): beyond the well-described interaction of the S-protein with host alveolar epithelia via the receptor human angiotensin-converting enzyme 2 (hACE2) (25; 47; 51), the S-protein also exerts pathological effects (changes in Proteome, E2F1, CREB, p65 activation) independent of ACE2. Furthermore, HMW-HA, which is naturally found in mammalian airways (1; 17), and can also be given pharmacologically (18), attenuates the deleterious effects of S-protein (**Figure 7**). Thus, our findings suggest a novel injury pathway and a novel treatment agent for COVID-19 associated lung injury.

**Figure 7.**
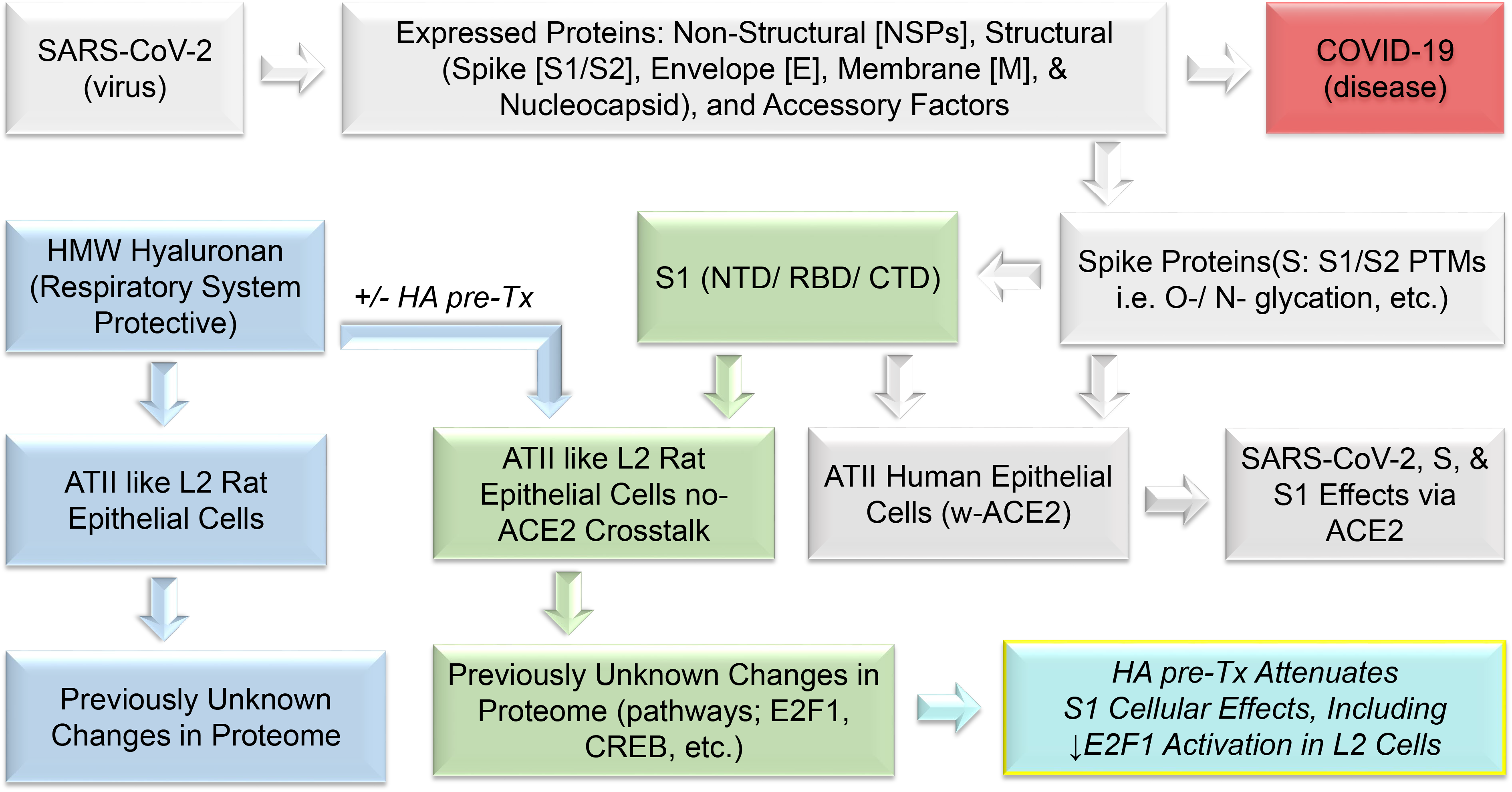
Experimental Workflow Similar to other coronaviruses, the makeup of SARS-CoV-2 is composed of four structural proteins: spike (S), envelope (E), membrane (M), and nucleocapsid (N) proteins. S, E, and M proteins are glycosylated, and the N protein is phosphorylated. The S protein is involved in the interaction with the host receptor human angiotensin-converting enzyme 2 (hACE2), which is also heavily glycosylated. The S protein makes up two parts the S1 and S2 spike proteins which are heavily glycosylated, and S1 contains the receptor binding domain (RBD). S1 is known to present with biological activities that are presumed largely to be due to interactions with hACEII in human ATII cells (grey background). However, previously unreported, S1 also induces significant changes in ATII-like Rat L2 cells that appear independent of hACEII as measured via proteomics analysis (green background). Similarly, and also previously unreported, HA induces significant changes in L2 cells (blue background) that greatly attenuate the deleterious effects of S1-Tx (turquois; bottom right). We have reported these observations here.

It is becoming increasingly evident that S-protein may impact cellular biology independent of interactions with the ACE2 receptor. Recent reports showed that the S-protein induces innate immune activation via toll-like receptors TLR2 and TLR4(2; 36; 71; 72) and *in silico* modeling suggests that it may direct interact with TLRs(13). TLR expression levels correlate with markers of organ injury in COVID19 (59) while genetic polymorphisms in TLR2 and TLR4 are associated with COVID19 outcomes (48; 60). Thus, the TLR axis may contribute significantly to the inflammation observed with SARS-CoV2 infection. Our result support this conclusion. Our unbiased analysis suggested p65, a central downstream mediator of TLR activation (23) as a major hub of our ptoteomic changes. Furthermore, other major hubs like CREB1 and E2F are modulators of TLR signaling (43; 65).

E2F are mostly described for their role in regulation of the cell cycle, with additional roles in metabolism and cell fate determination. However, there is an emerging pattern of E2F playing a role in inflammation as well (43; 68) In particular, recent papers suggest the E2F activation may play a role in COVID19 inflammatory cascade and cytokine storm (6). Our unbiased approach supports this conclusion and further suggests that this effect may be impacting the innate immune pathway, at least partly downstream of the S1-protein. It is also interesting that we identified ROCK/RhoA as a top enriched gene pathway. Recent papers highlight a role for these kinases in cellular injury and disruption of the cell barrier in COVID19 (7; 9), thus supporting that our findings have biological relevance. Importantly, we have shown in several papers that RhoA/ROCK mediate non-infectious lung injury(4; 21; 34; 39; 40; 73), a process which can be inhibited by HMWHA, a striking parallel to the finding in this work.

In aggregate, our proteomic results support that S-protein indices a robust innate immune activation cascade that can be a target of therapeutic interventions. HMWHA is such a potential intervention. It is naturally present in mammalian airways (1; 17) and has been extensively studies in its ability to modulate TLR signaling (62). Importantly, therapeutic application of HMW-HA has been shown to have beneficial effects, including in infectious and inflammatory lung disease(18; 21; 34; 73). Given that it is a cheap, natural, biological compound with essentially no adverse effects, HMWHA is therefore attractive as a first-line agent in emerging infectious diseases like COVID-19.

The addition of HMWHA to L2 cells, even in the absence of S-protein, resulted in significant changes to the proteome. Systems biology analysis led to the identification of a number of significant Network Hubs and GO Biological Processes that include strong associations with E2F1, E2F5, DNA Replication, Cell Cycle Regulation, p53, NF-kB, and DNA Repair. Given that a number of reports have presented beneficial effects of HMW-HA on pulmonary pathologies (19; 39; 41; 73), it is possible that some of the effects of HMWHA in our model are mediated by a pro-homeostatic effect of HMWHA that is not specific to S-protein. Indeed, HMW-HA is known to ameliorate inflammation downstream of LPS/TLR4 signaling(12), thus again supporting a global activity pattern as an anti-inflammatory mediator.

In conclusion, this work demonstrates that S-protein exposure in epithelial cells leads to a robust proteomic response, even in the absence of ACE2 engagement. We identify several novel signaling hubs, including E2F1, CREB, and ROCK/RhoA, and demonstrate that HMWHA, a natural component of airway lining, can ameliorate these changes and promote cellular homeostasis. Our findings shed new light into the pathogenesis of COVID19 and suggest novel angles of treatment that will ameliorate cell injury and lung dysfunction is this devastating disease.

## DATA SHARING

The mass spectrometry proteomics data will be deposited to the ProteomeXchange Consortium via the PRIDE partner repository upon acceptance.

## Acknowledgments

The authors are grateful to Dr. Israr Ahmad, Zhihong Yu for technical assistance with these studies.

---

**Supplemental Table 1.**
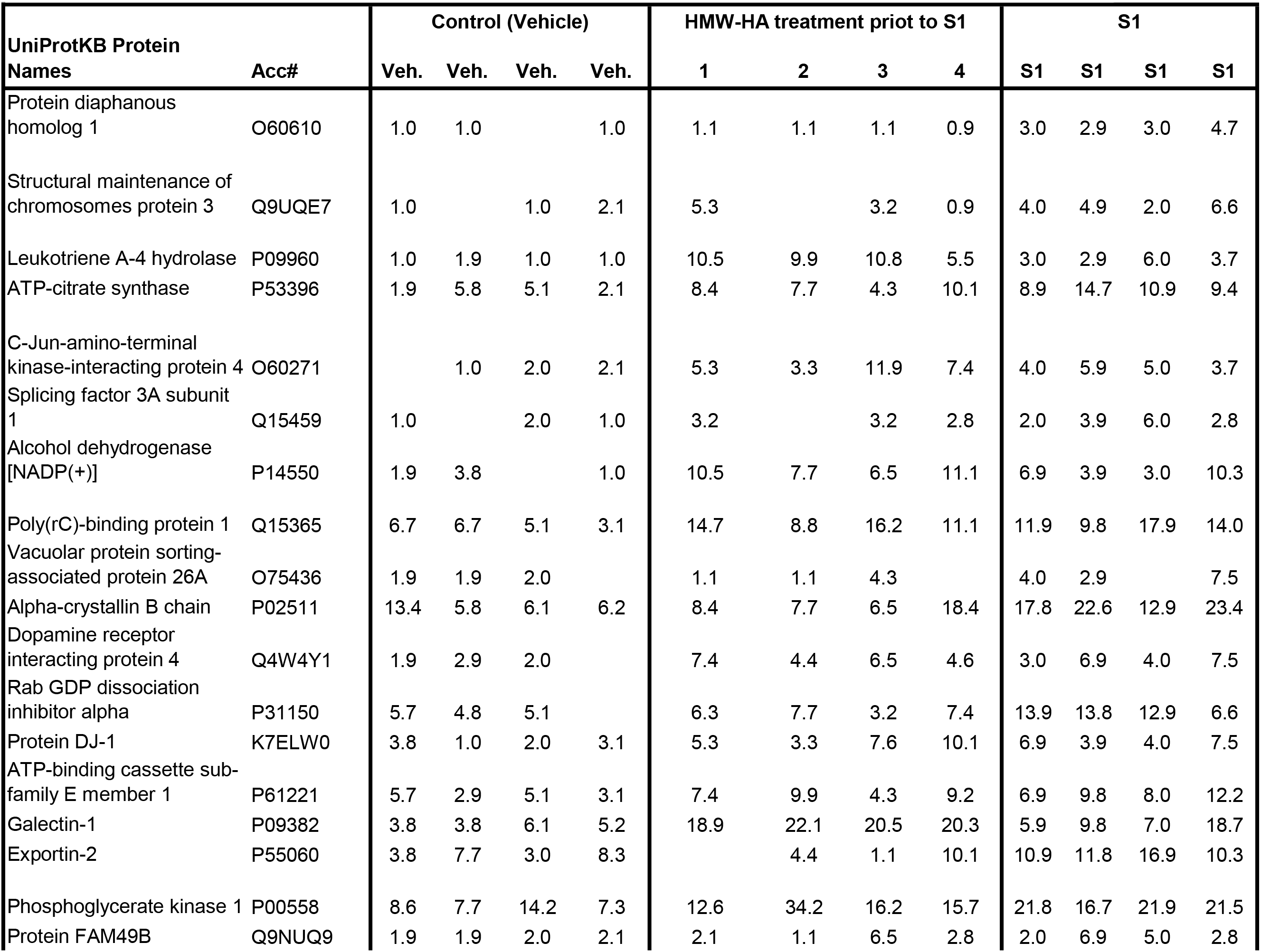
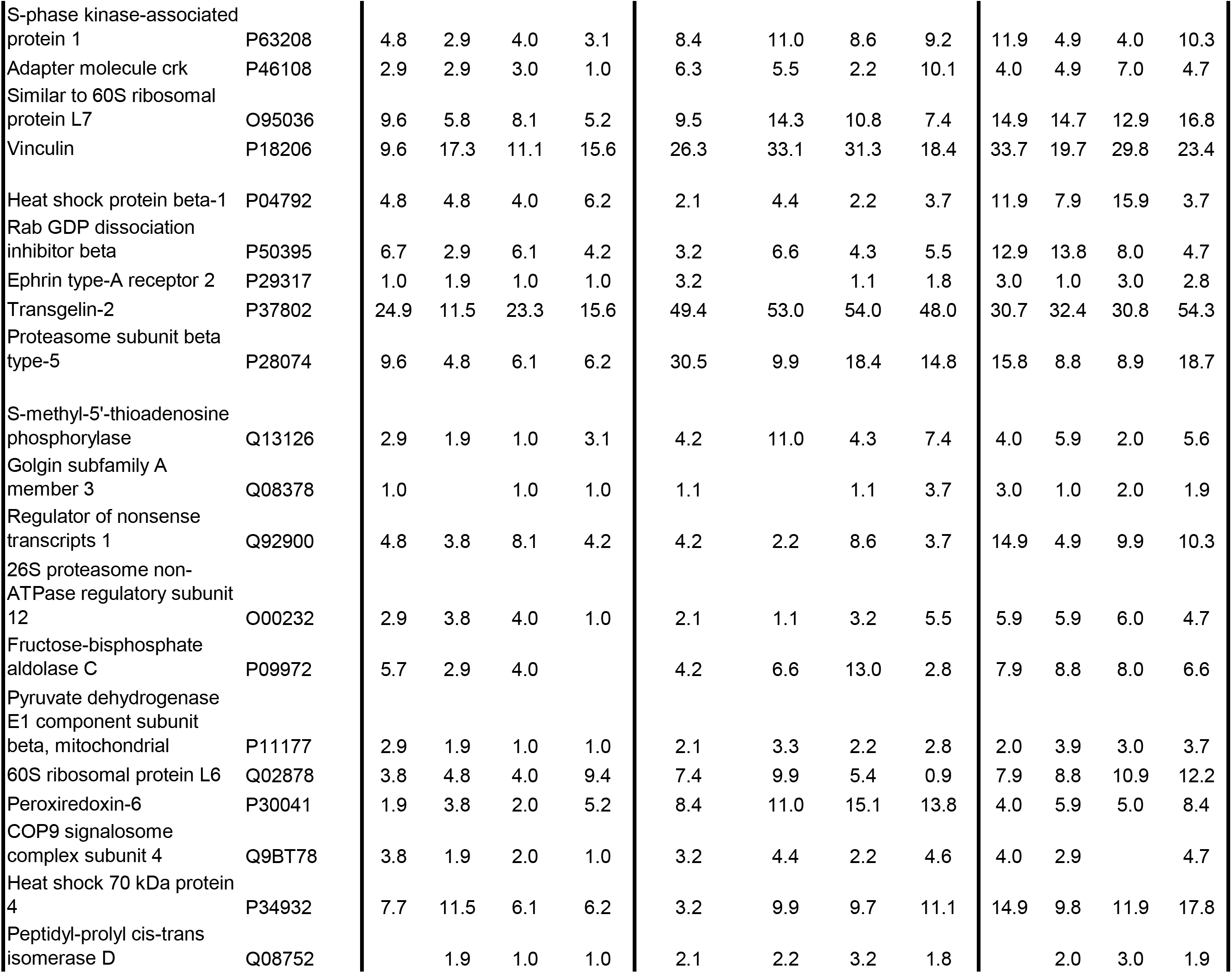
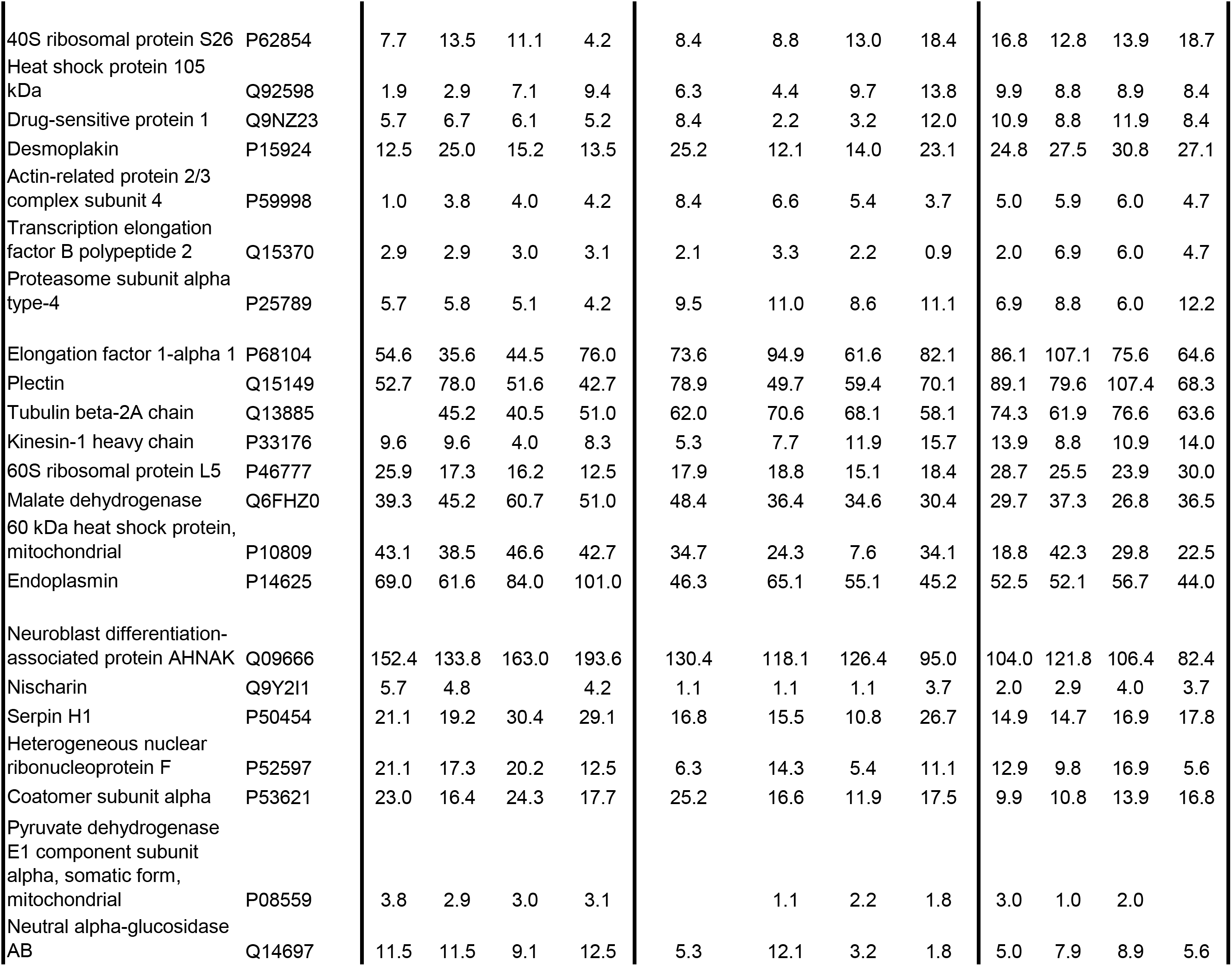
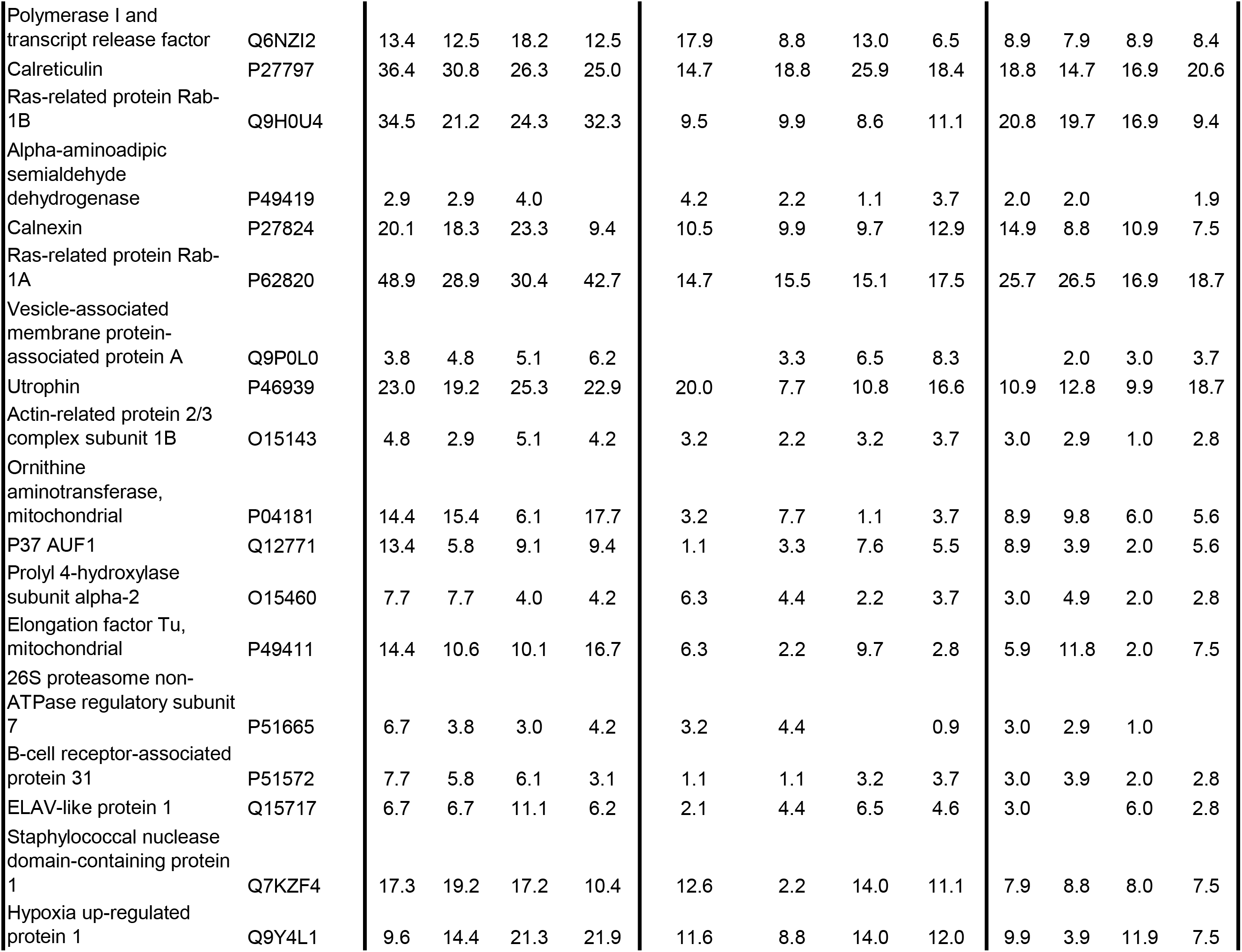
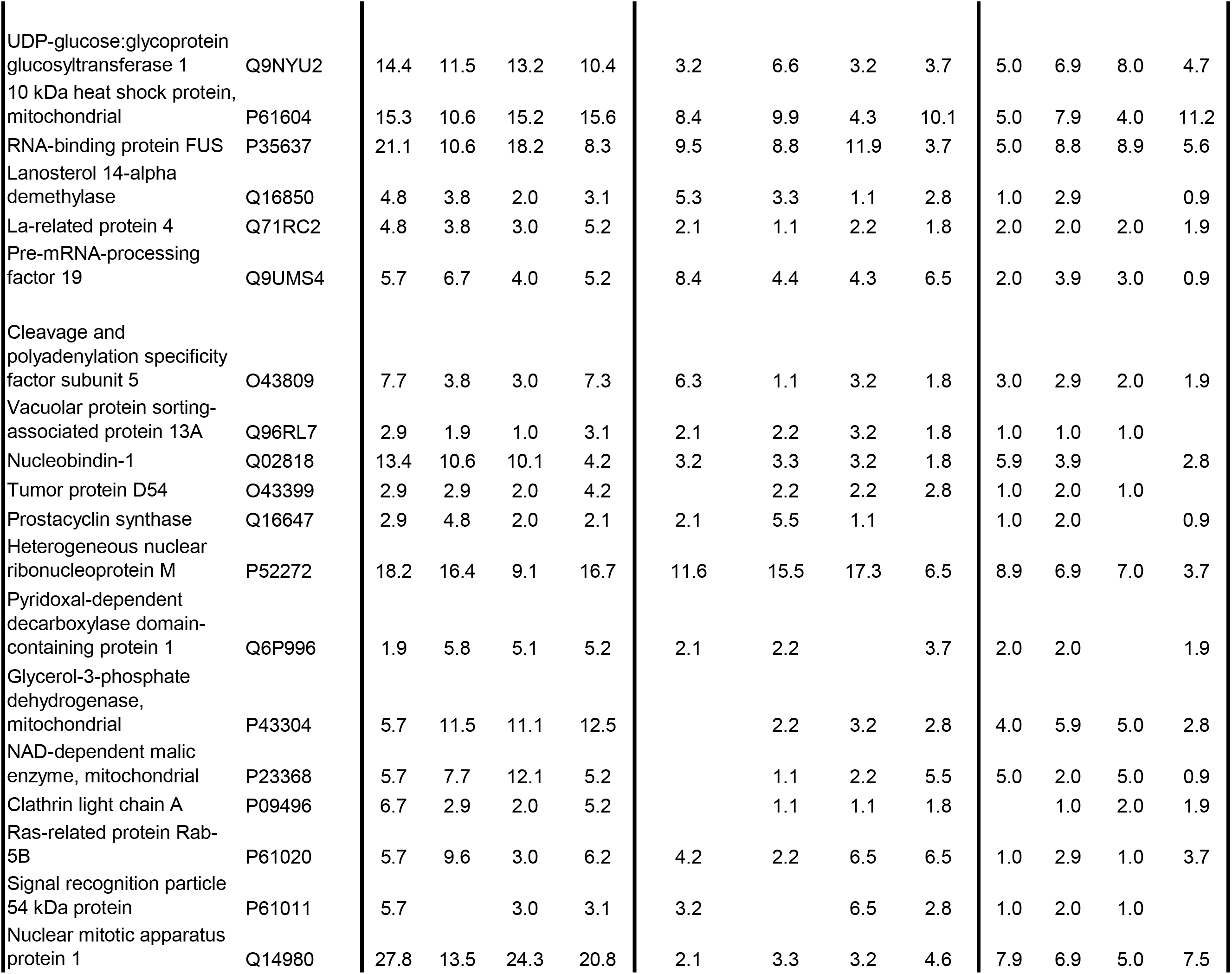
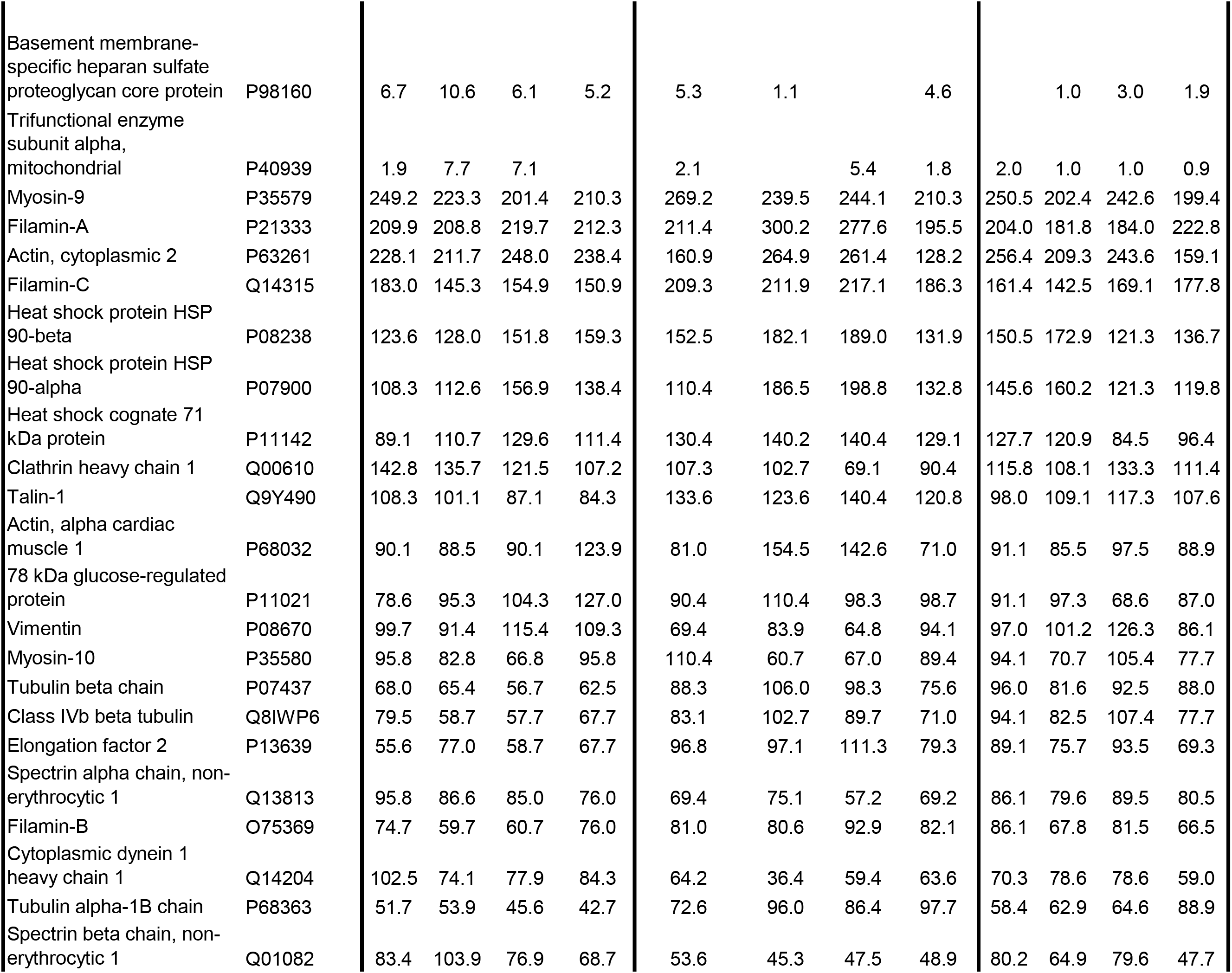
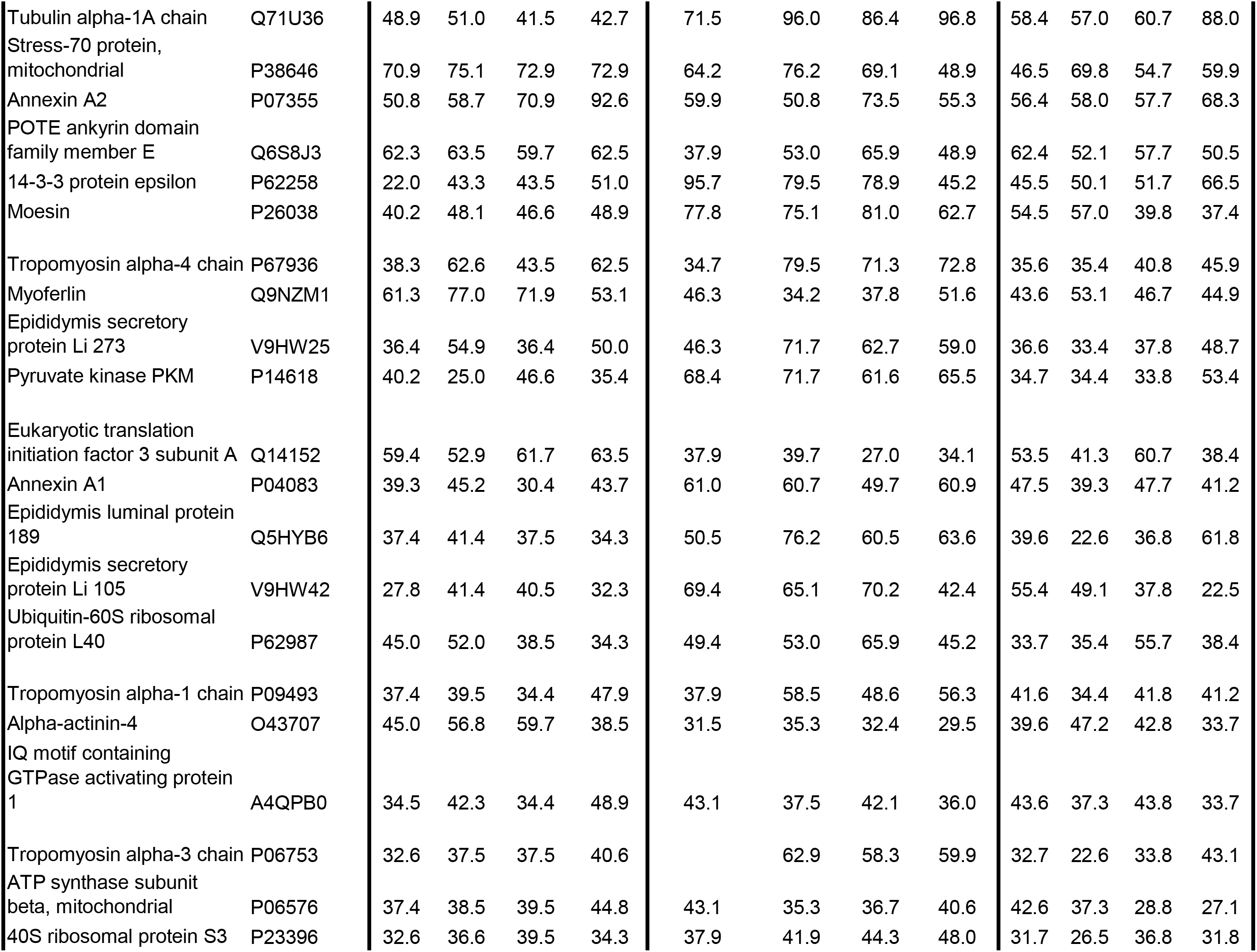
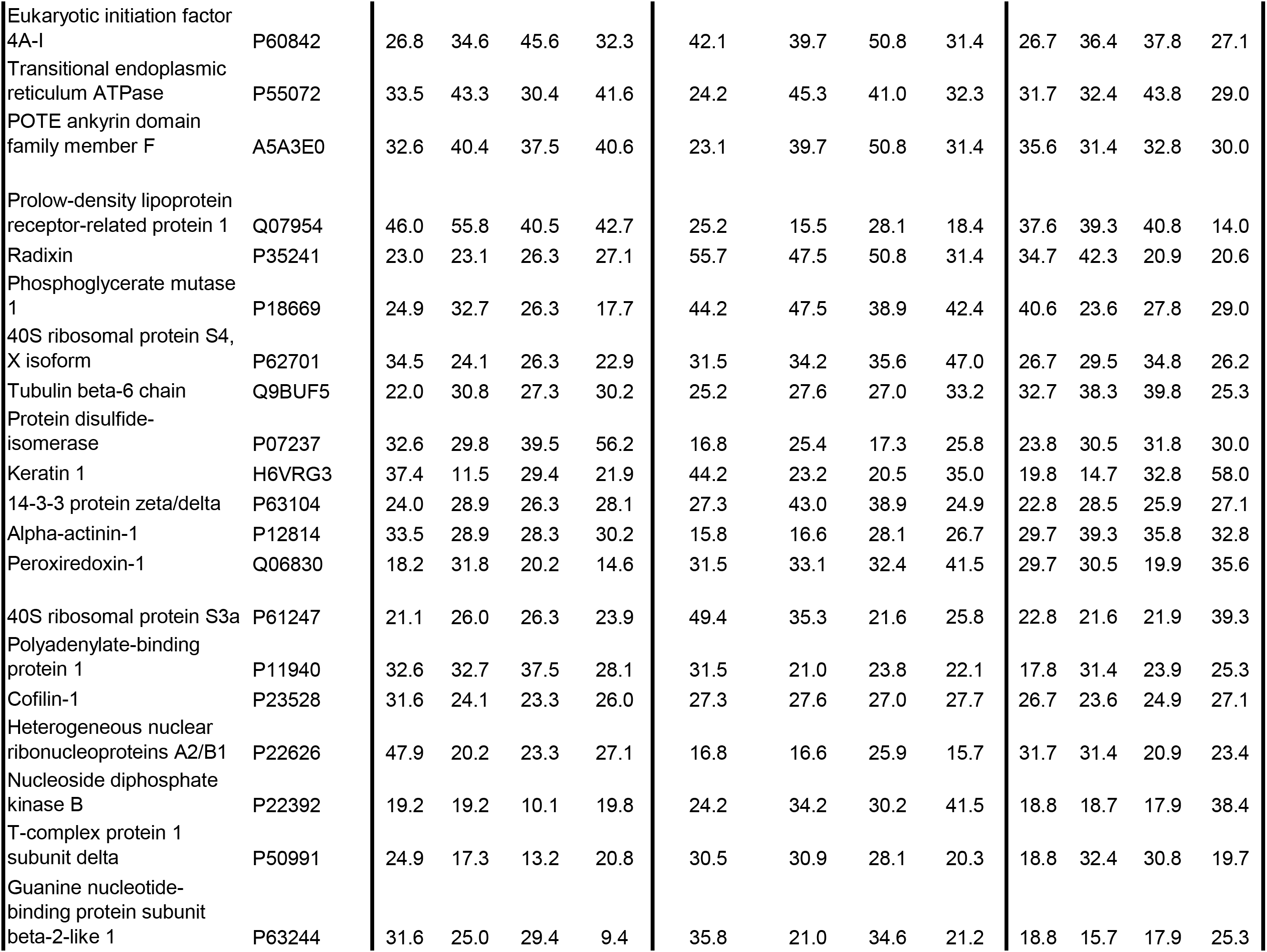
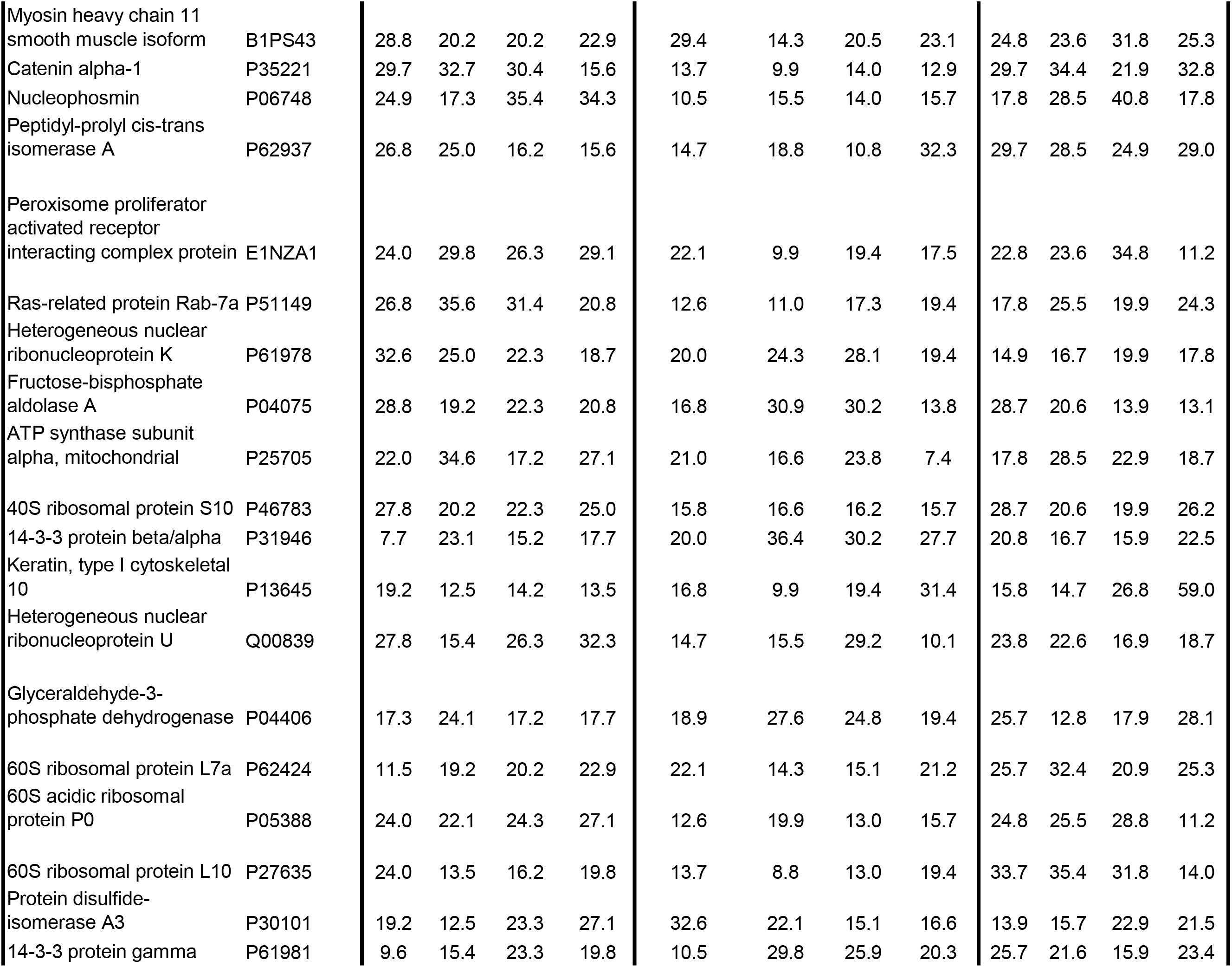
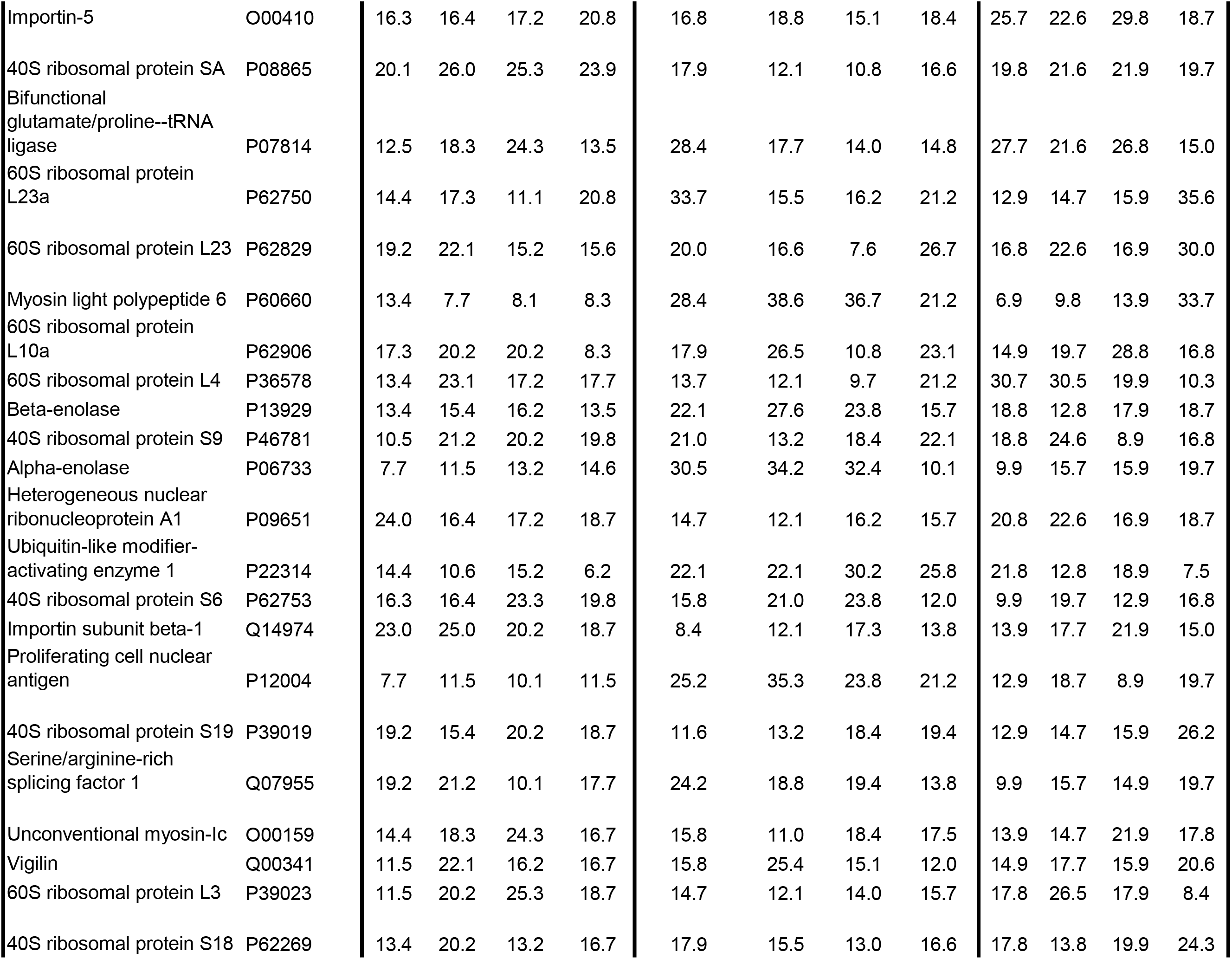
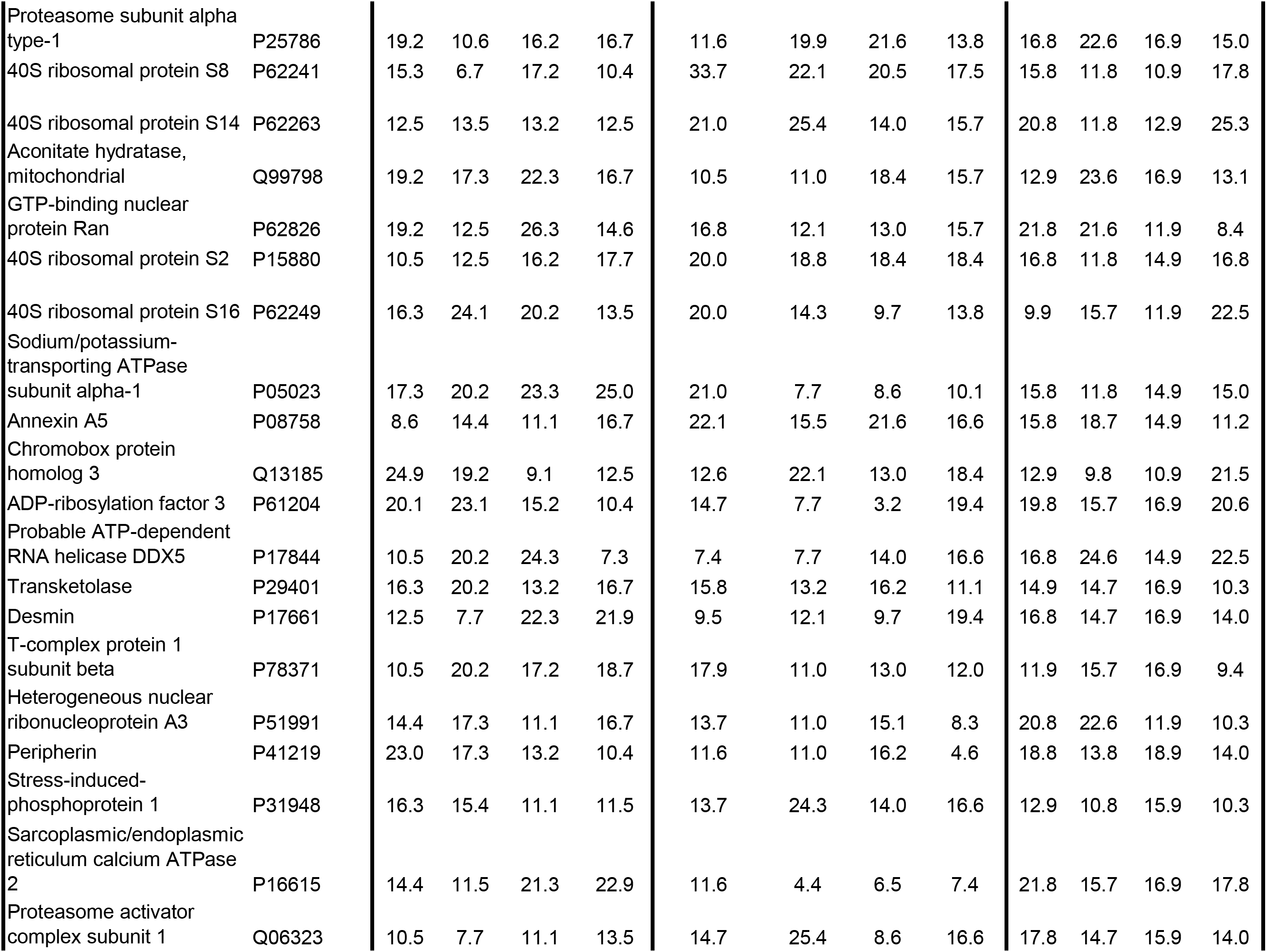
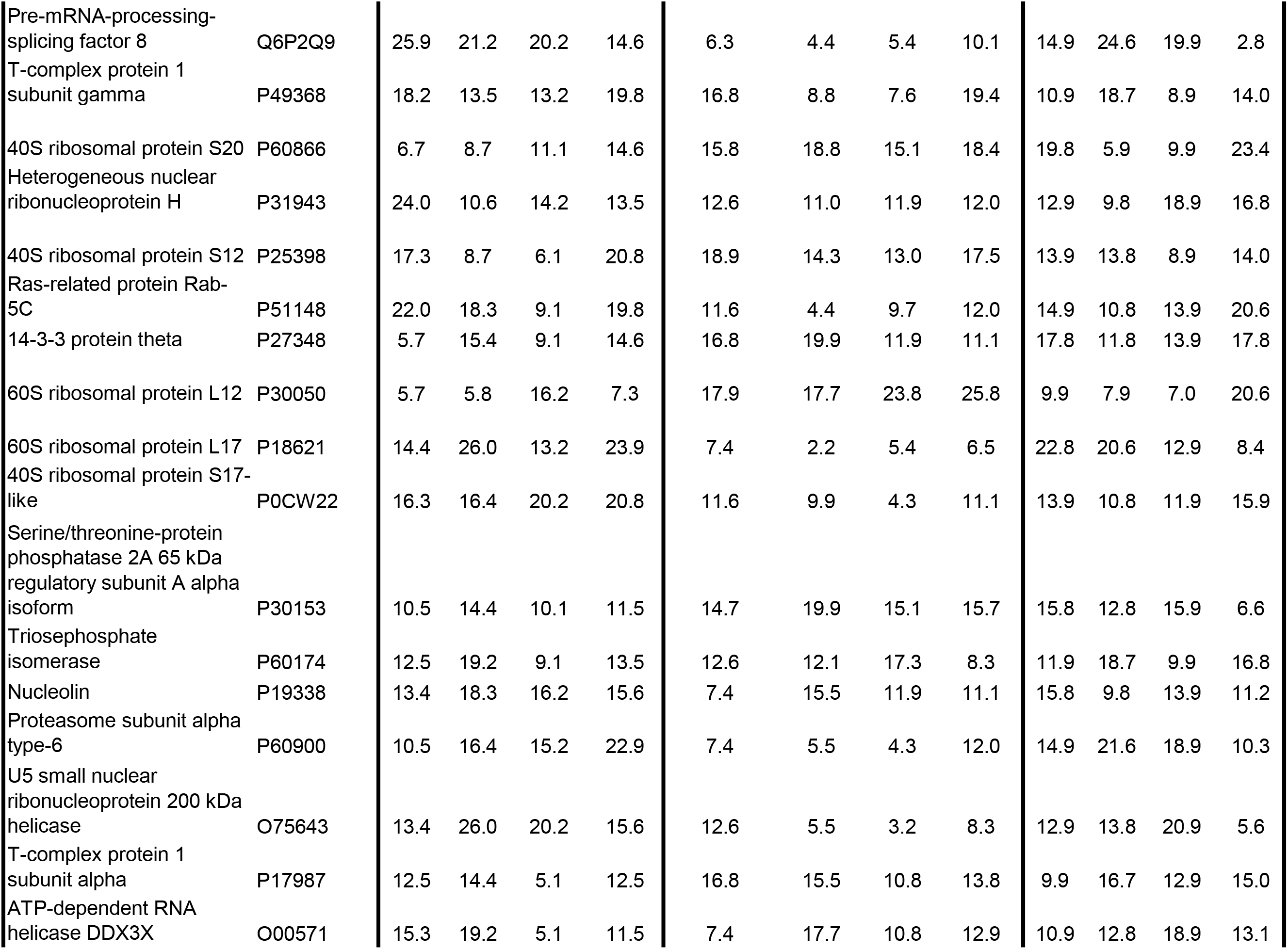
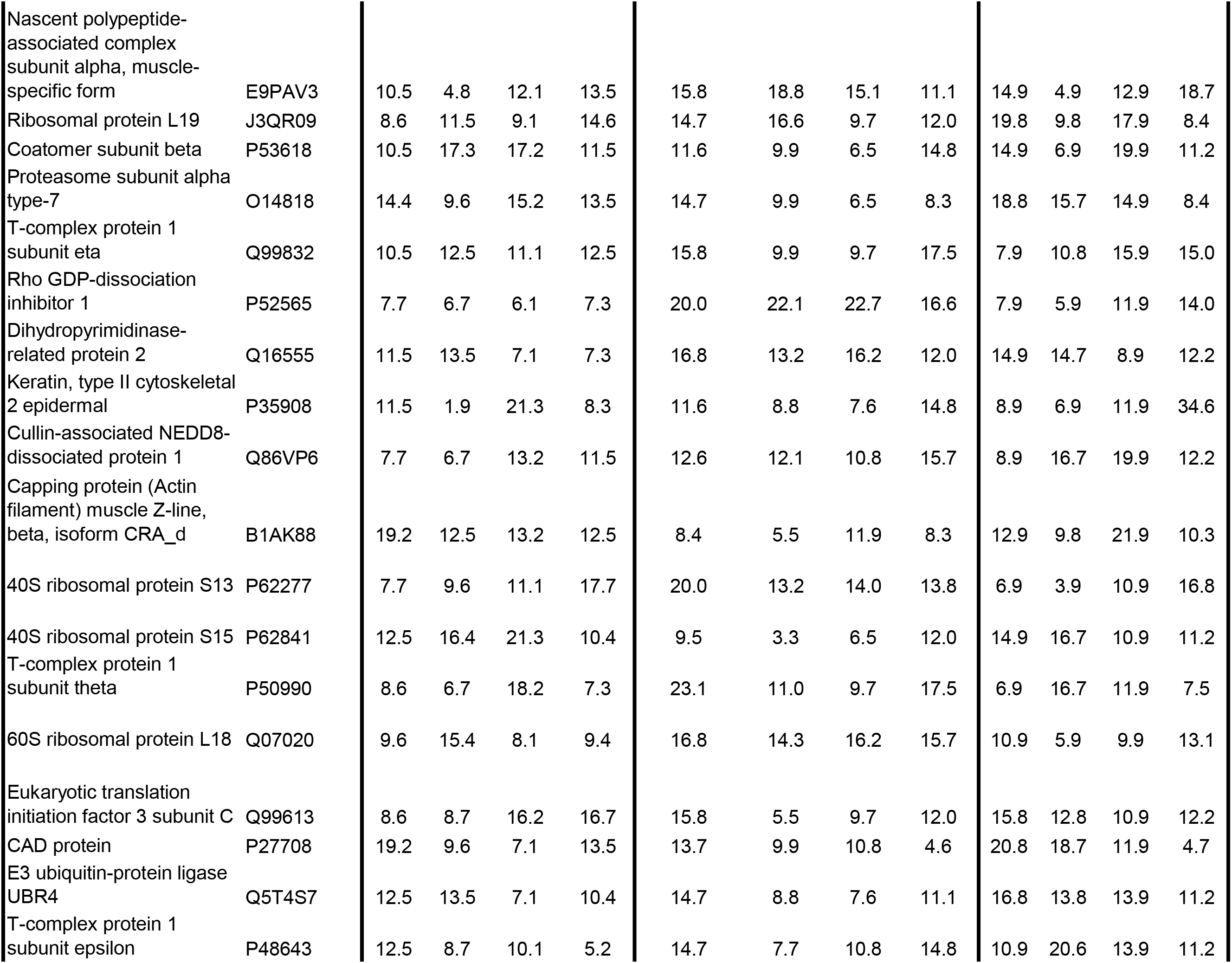
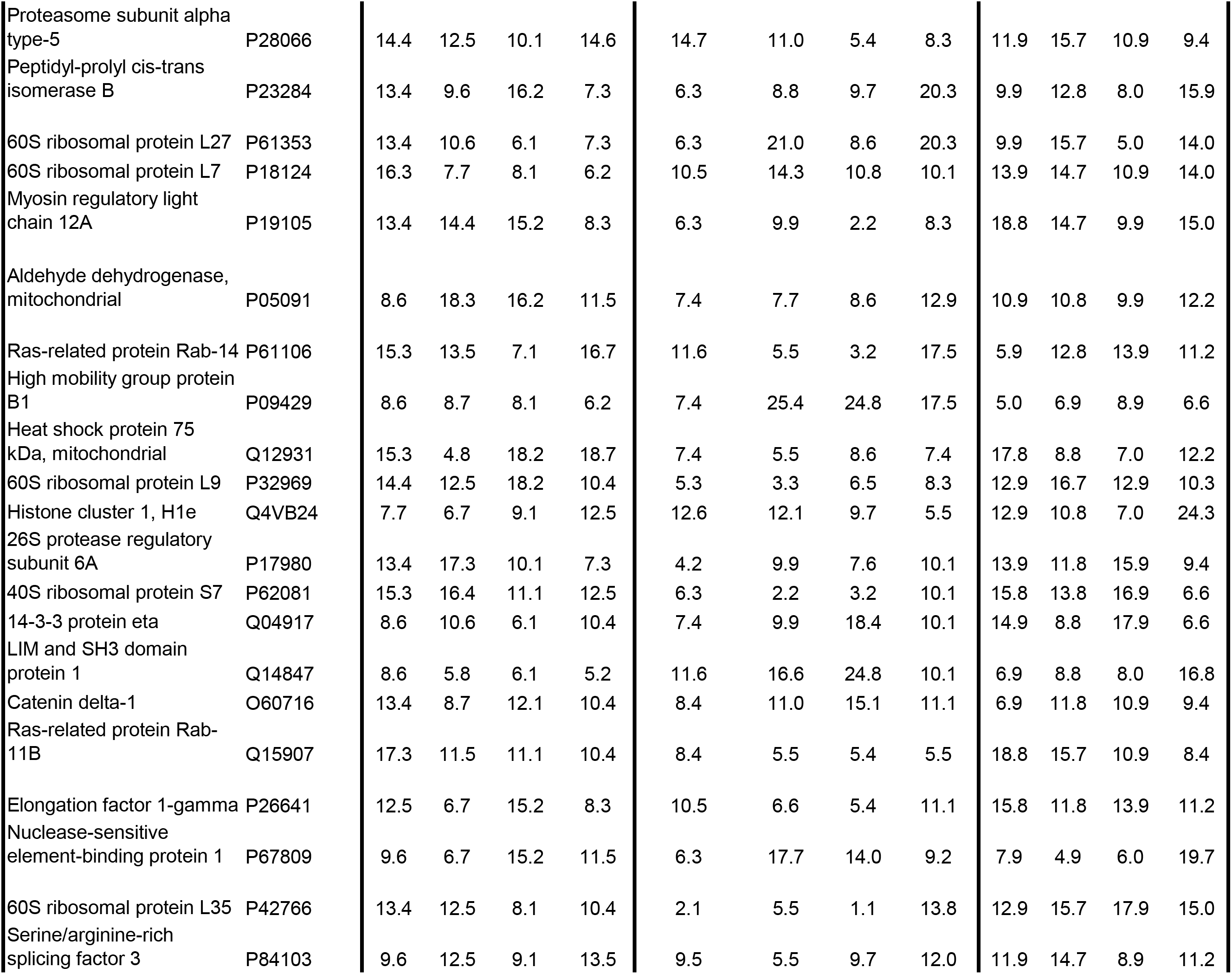
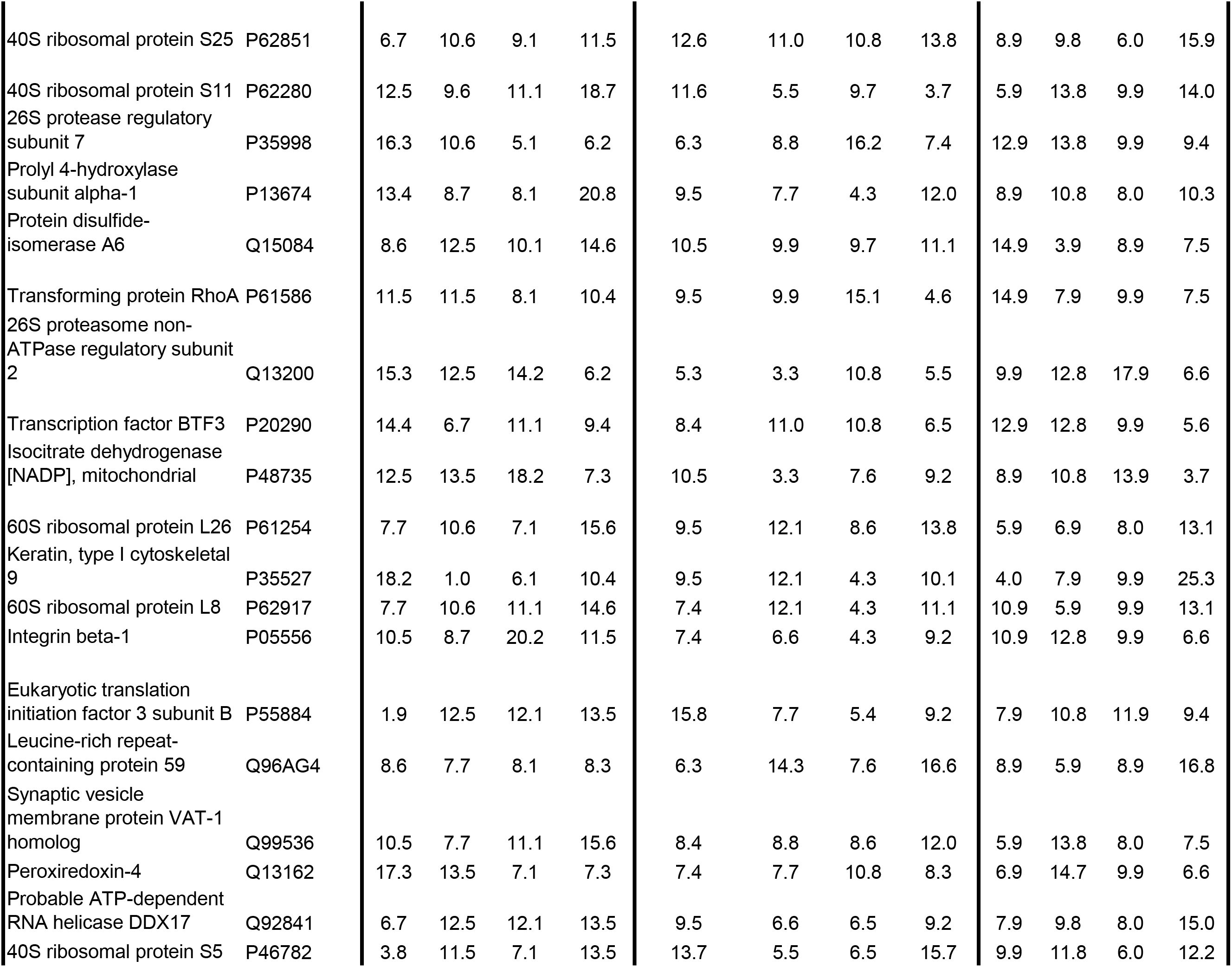
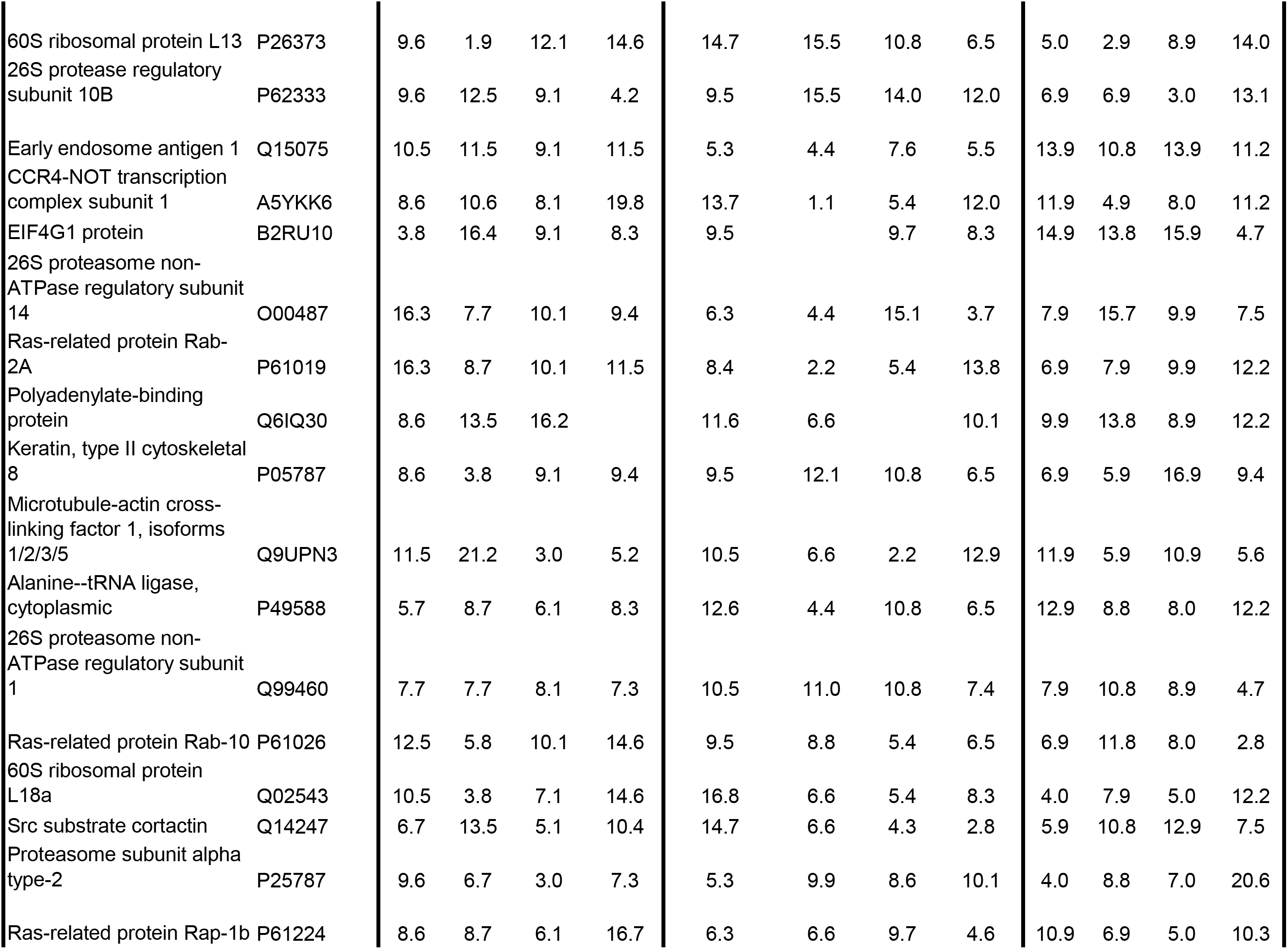
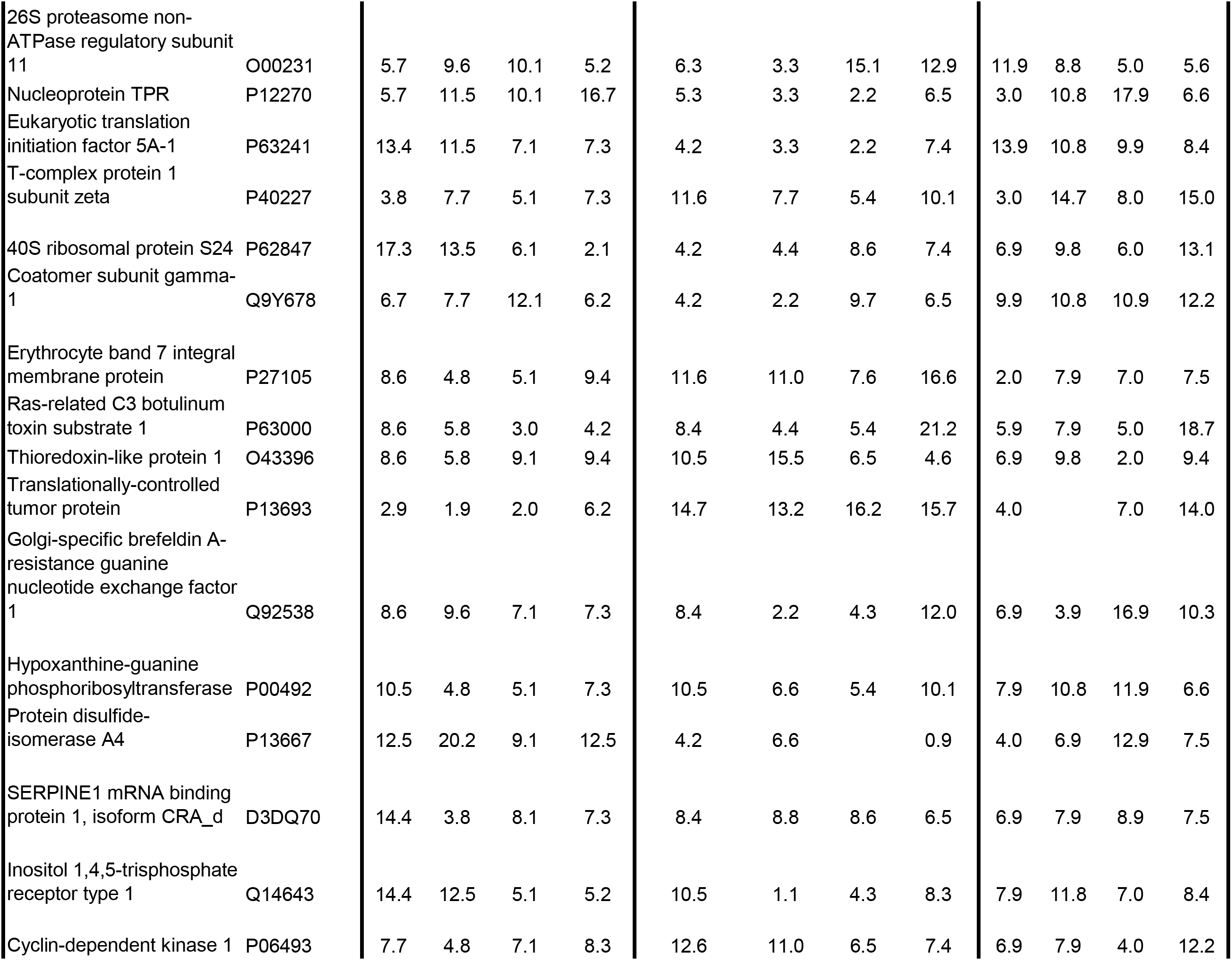
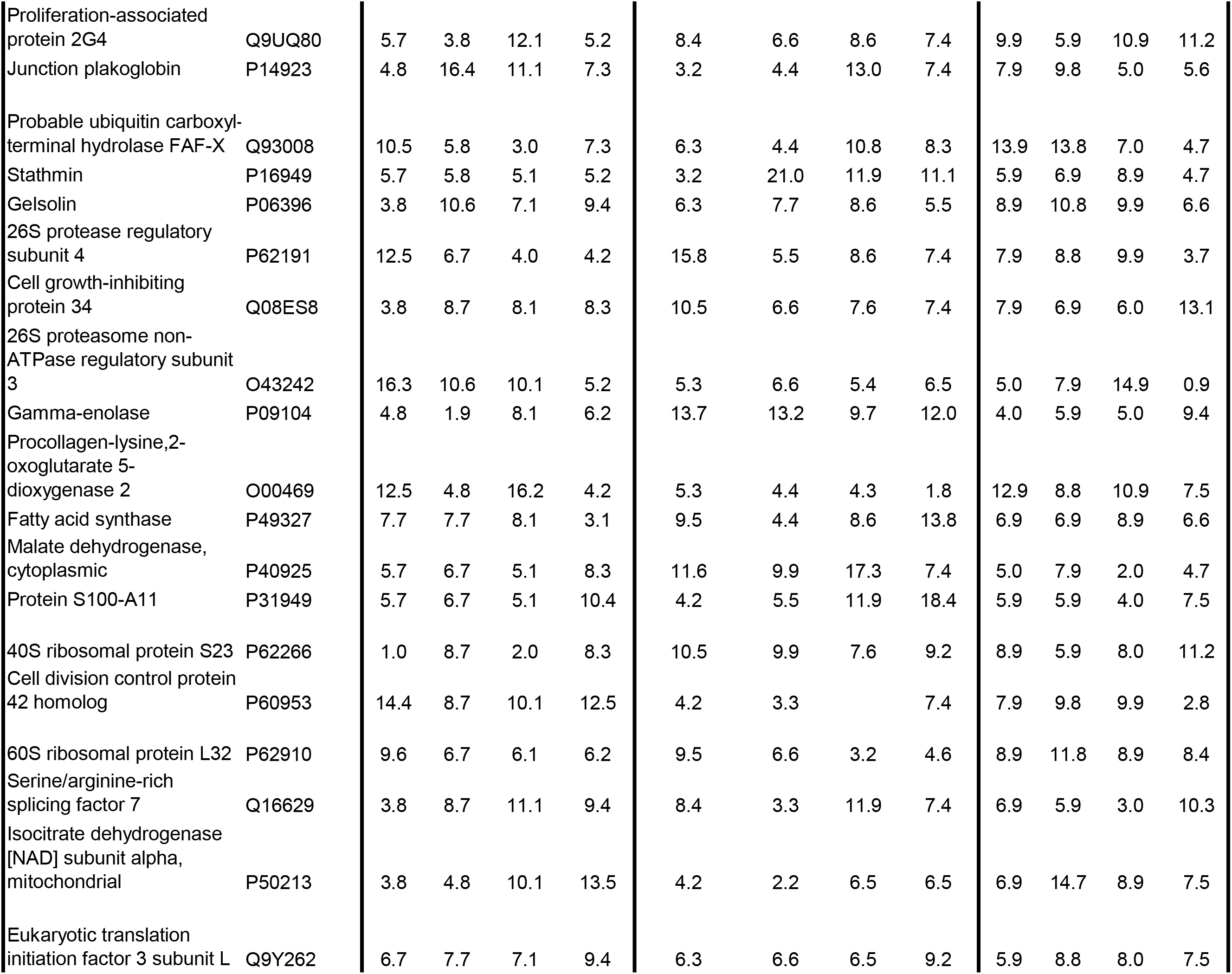
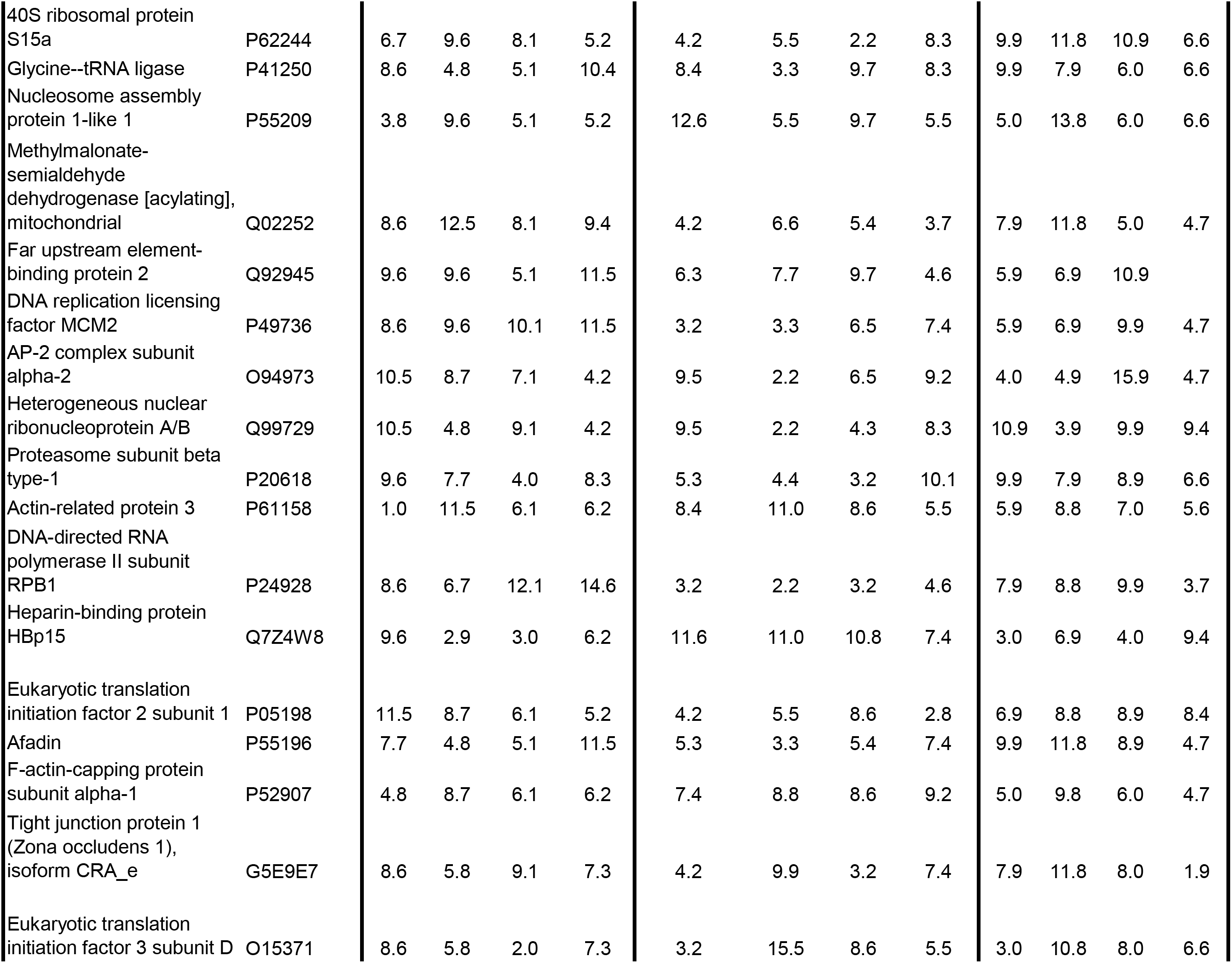
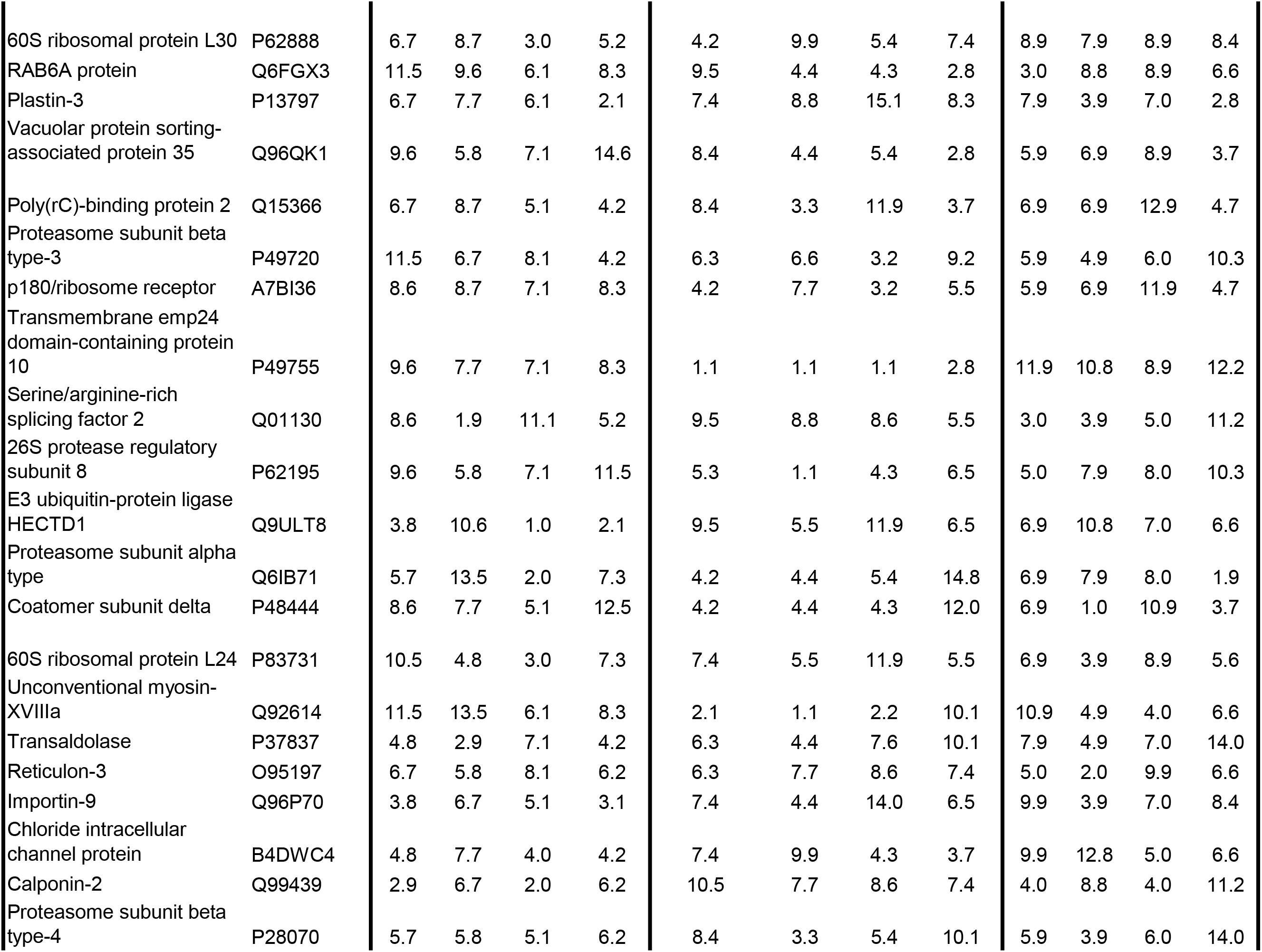
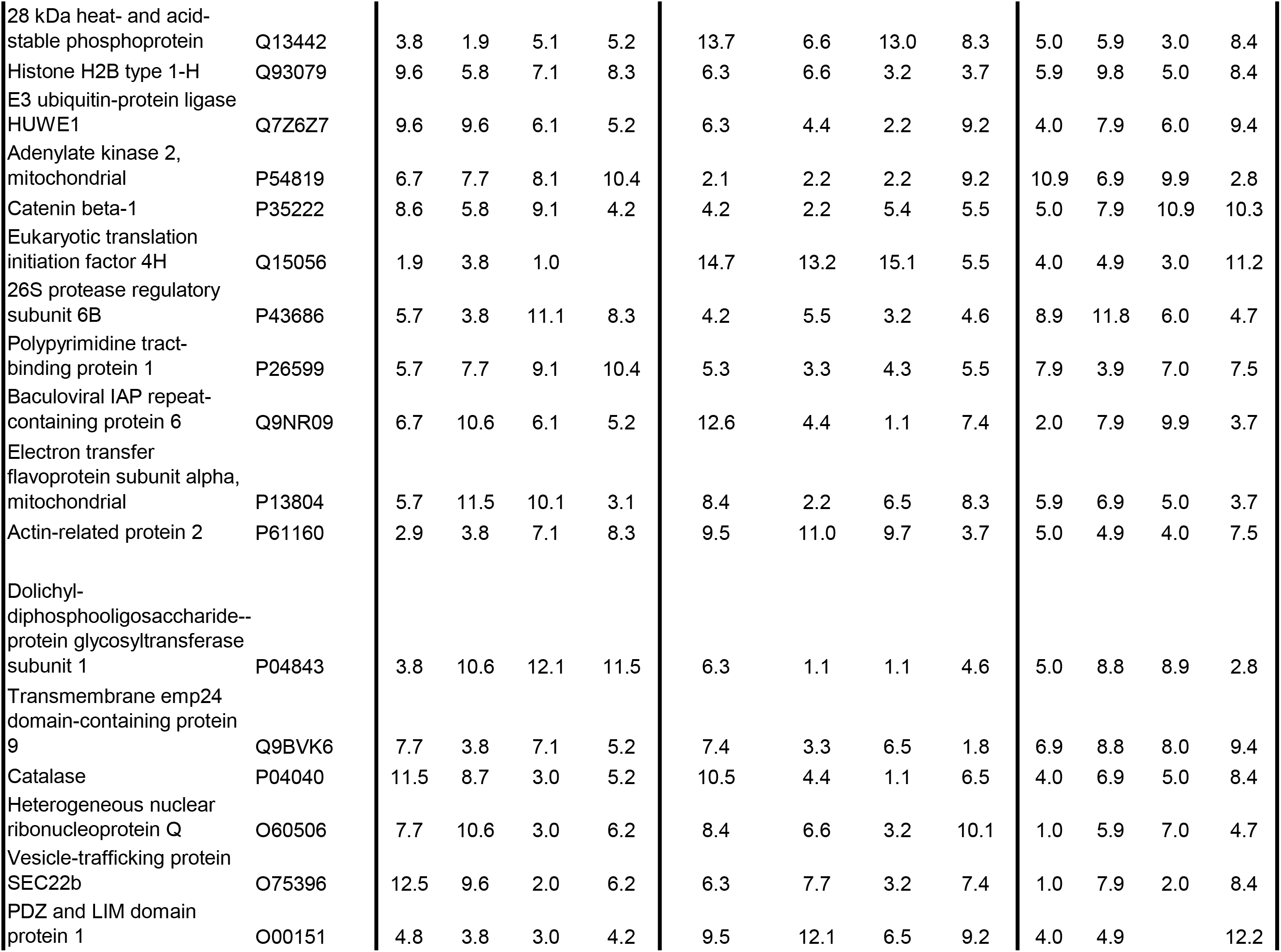
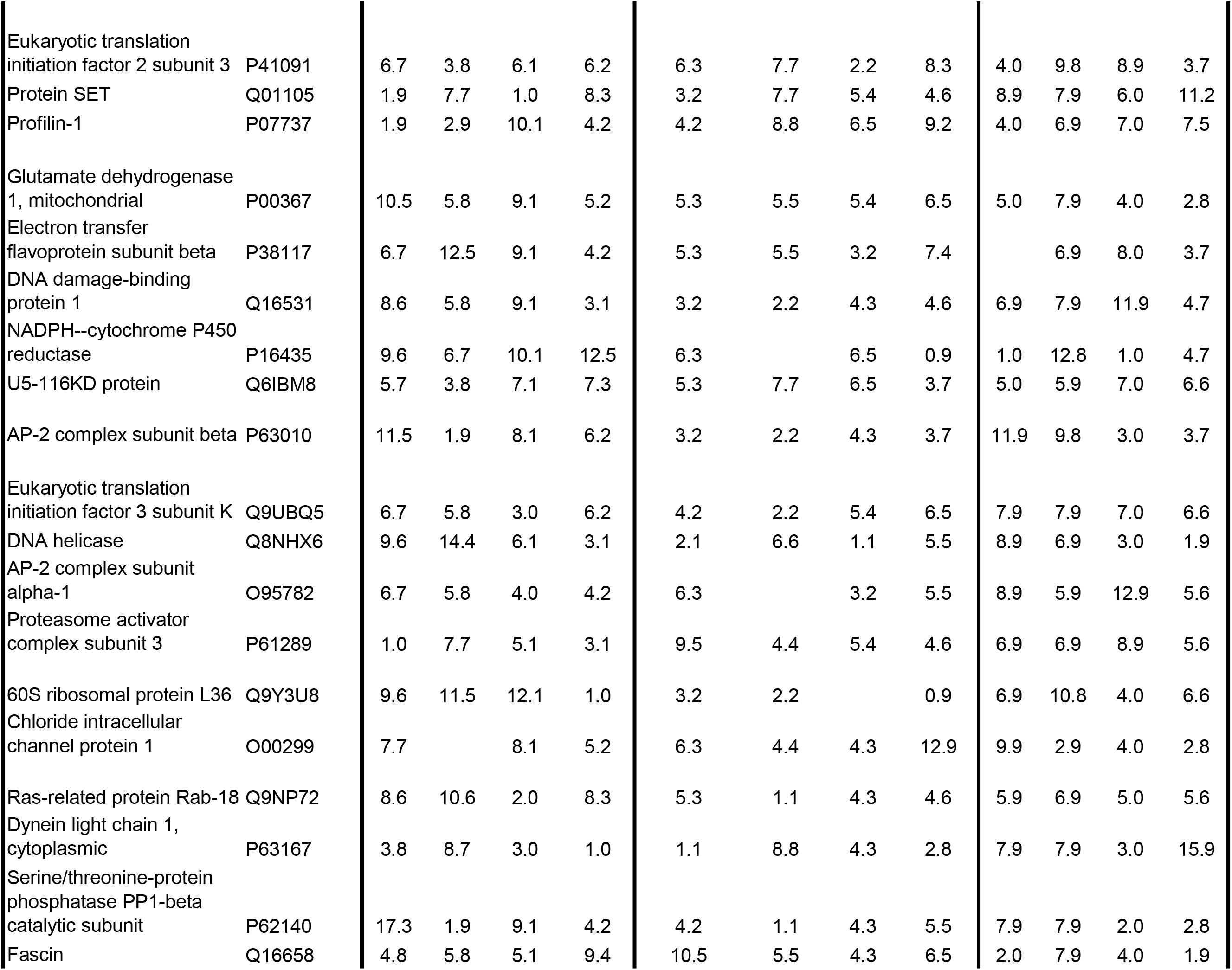
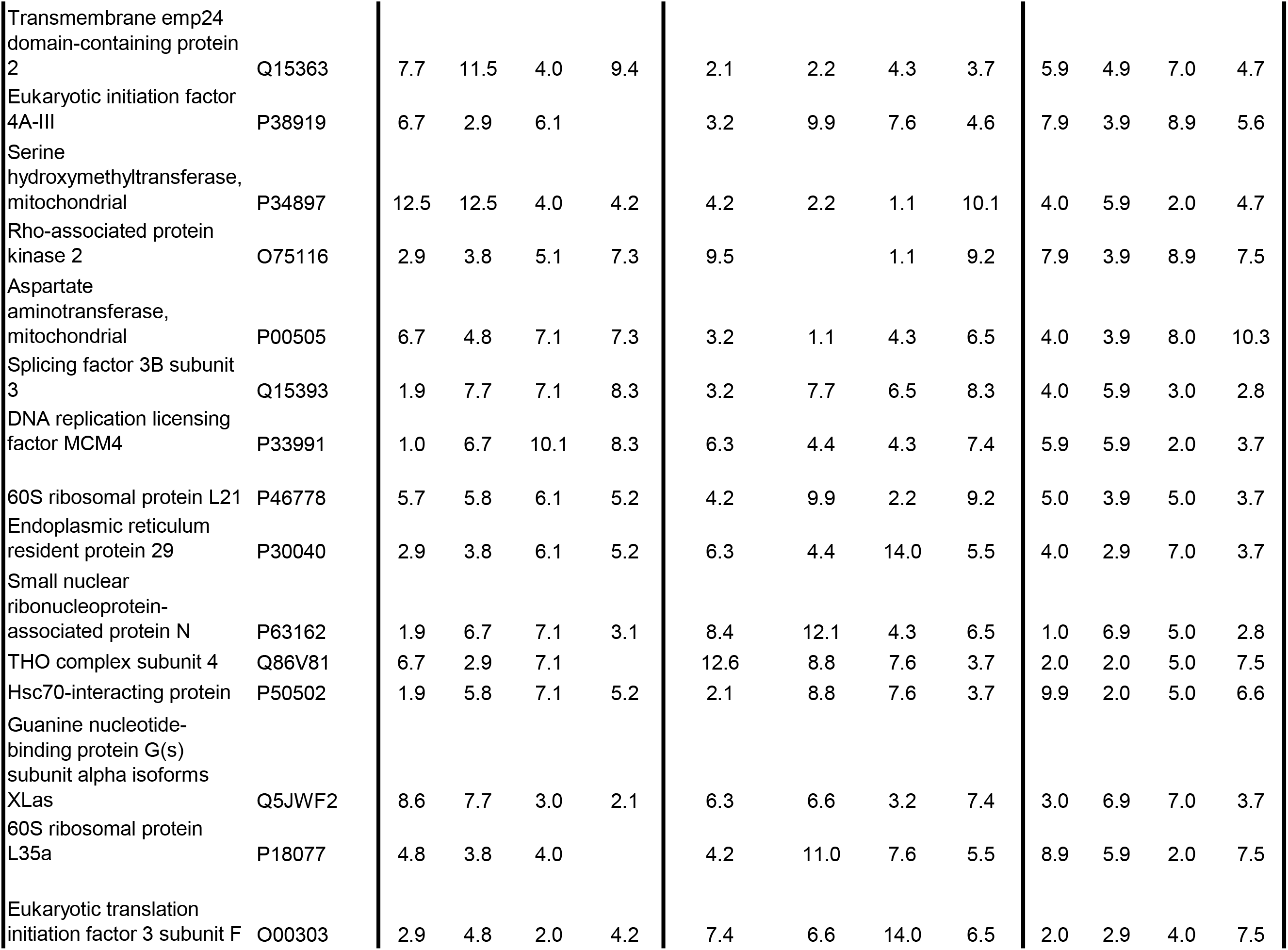
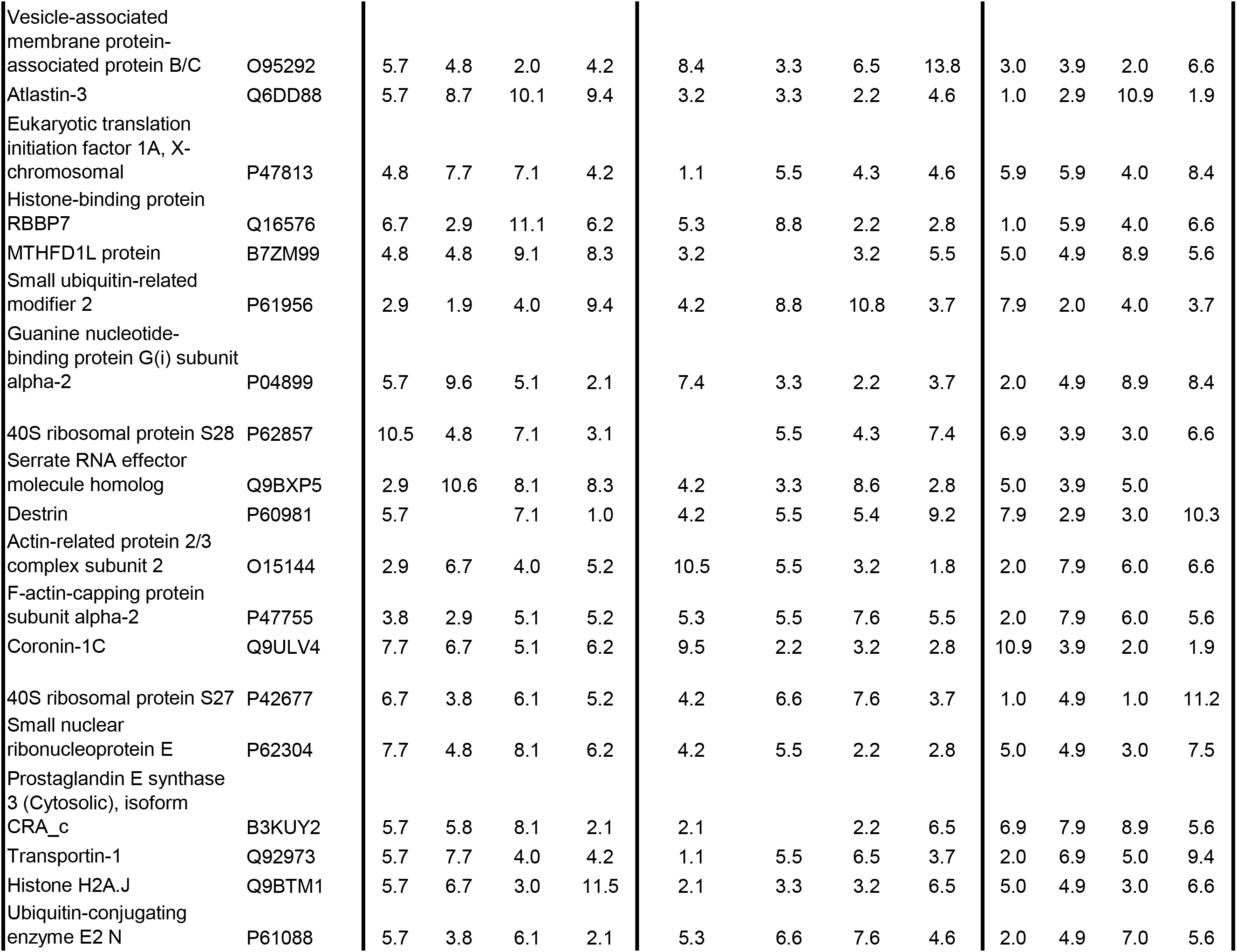
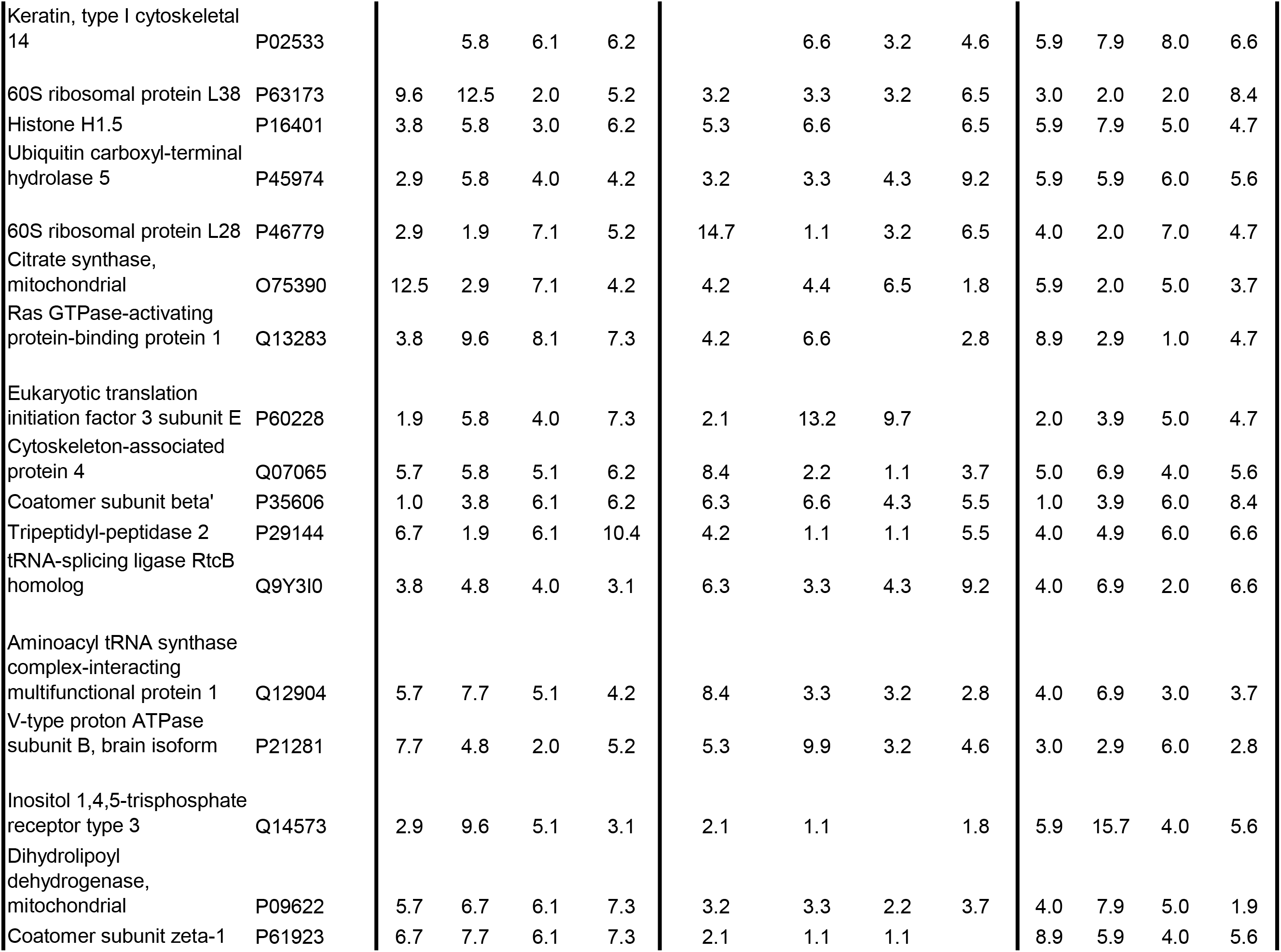
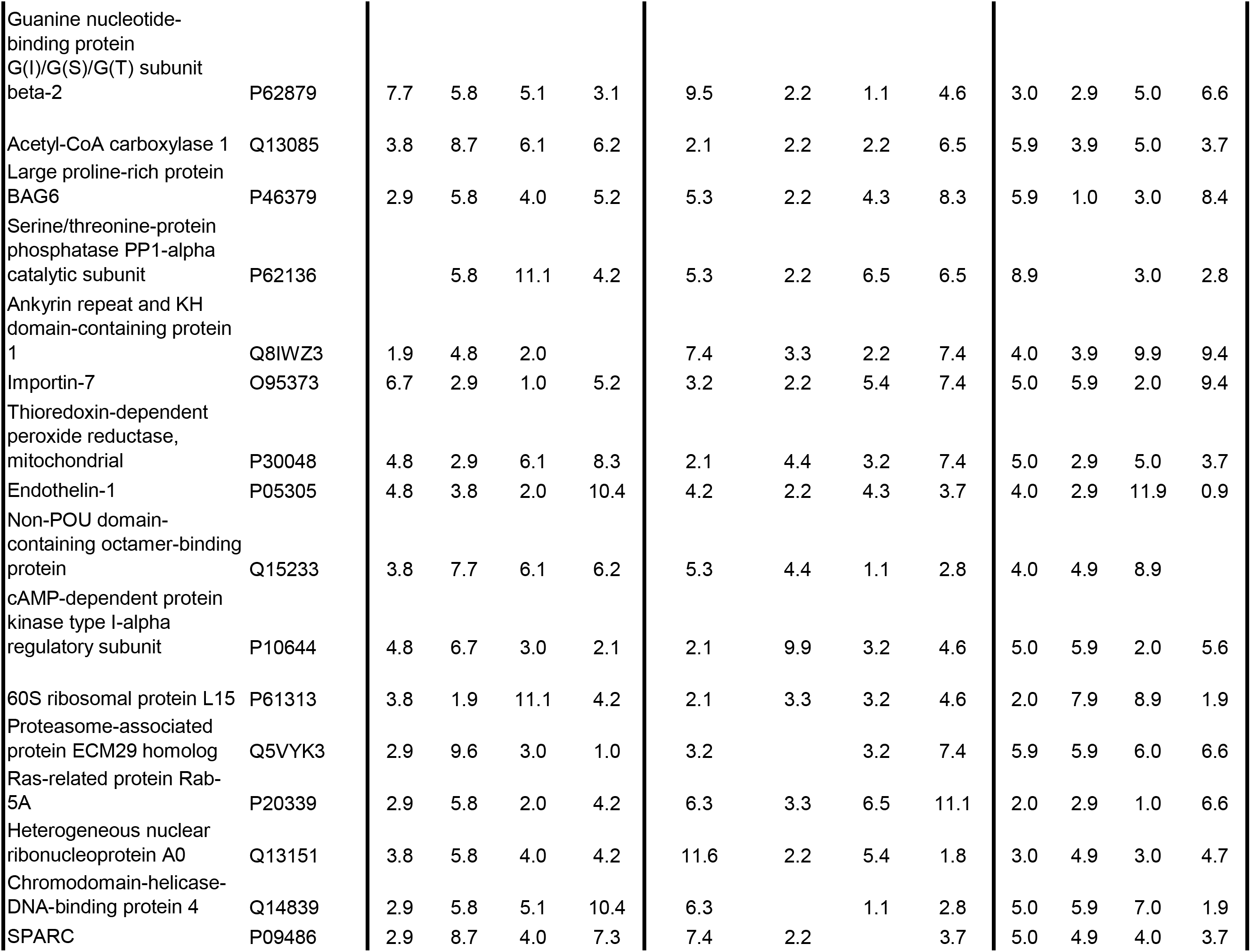
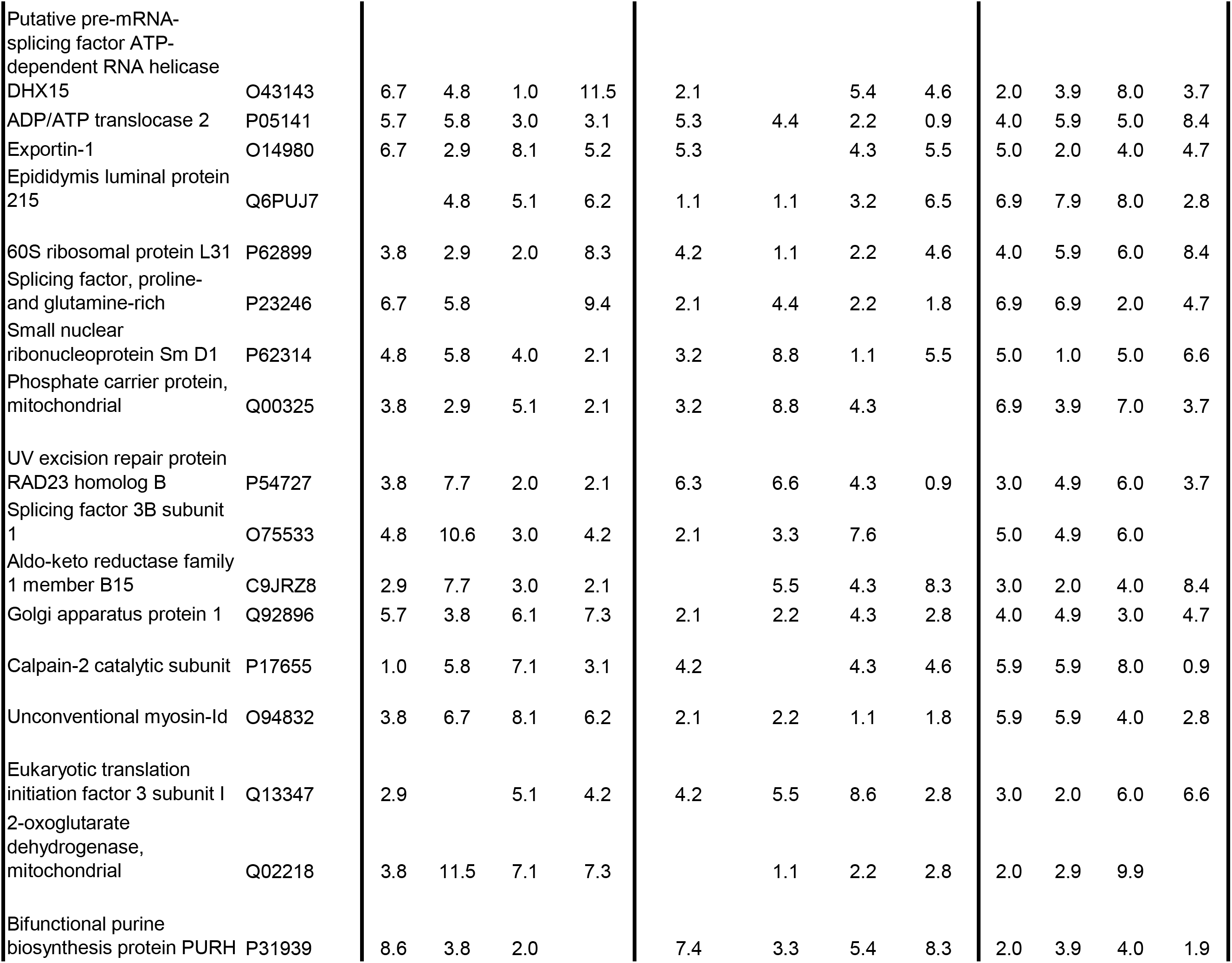
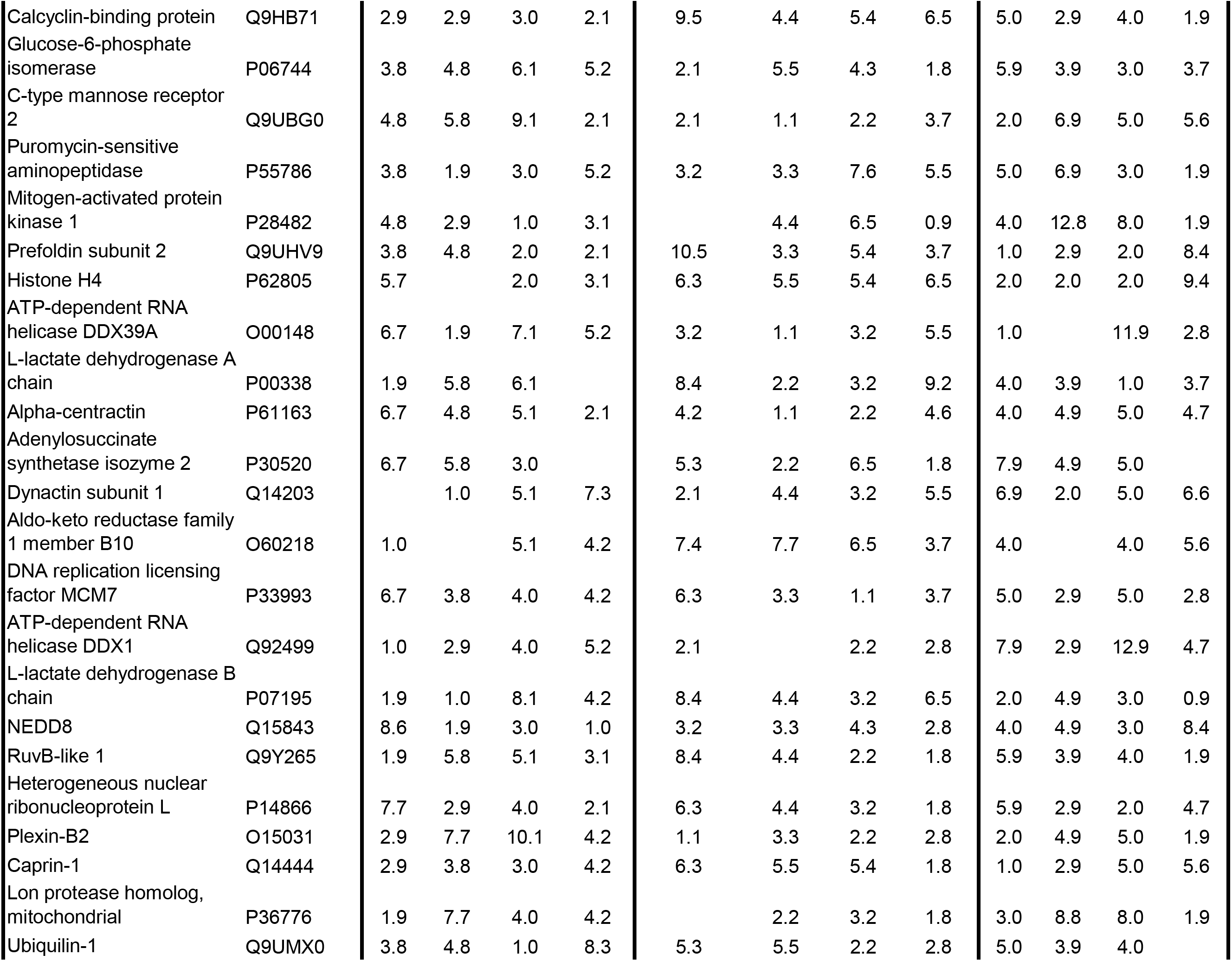
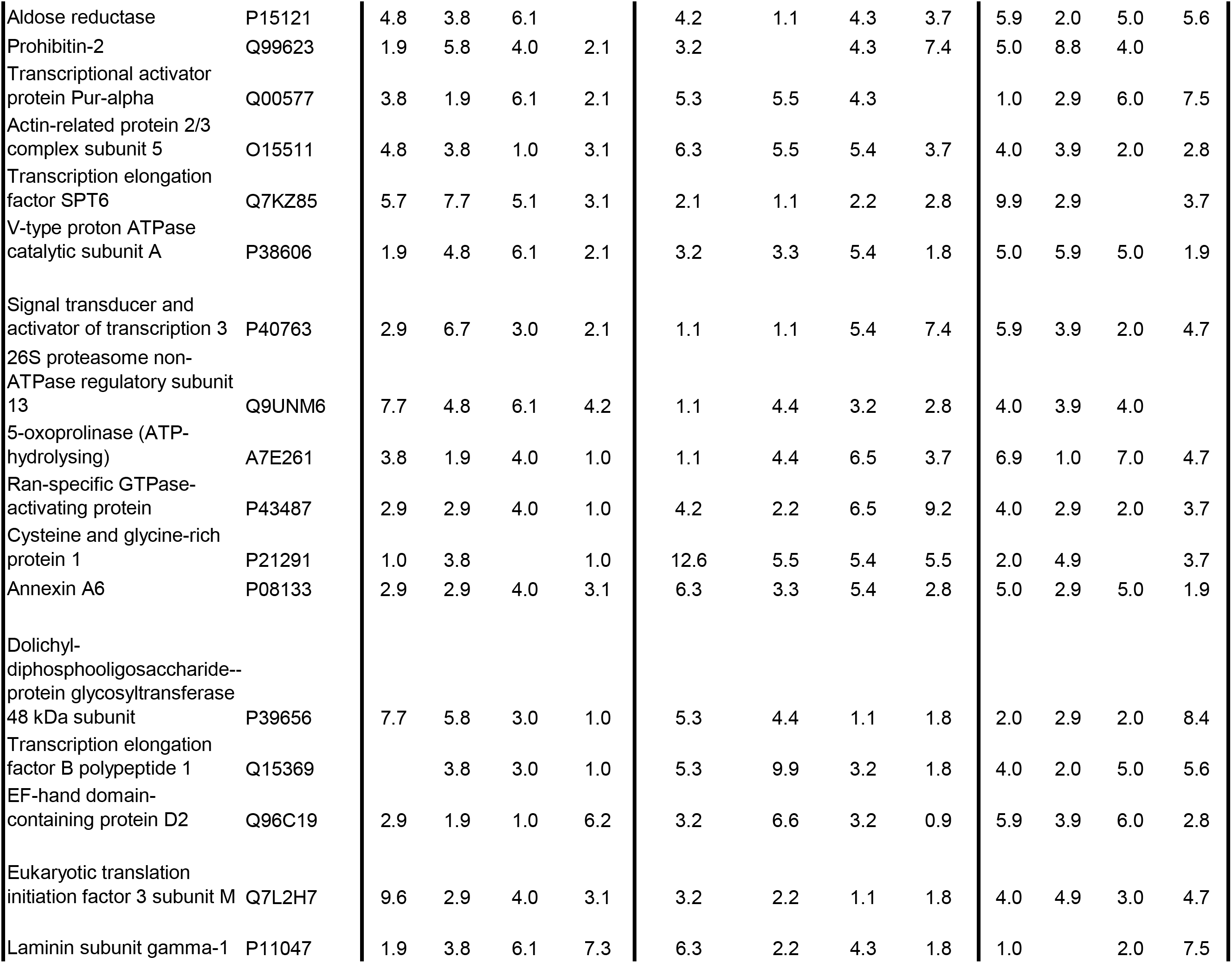
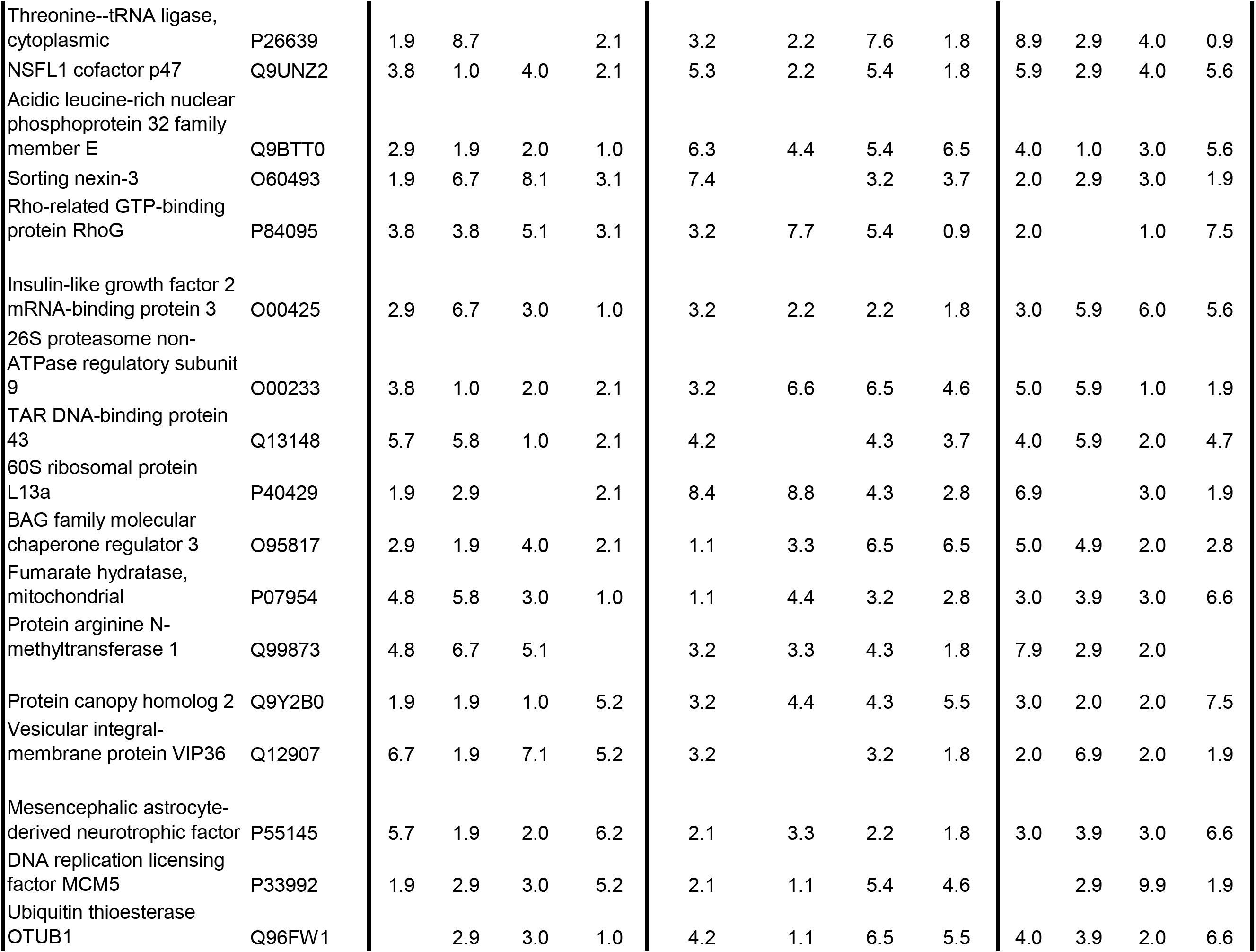
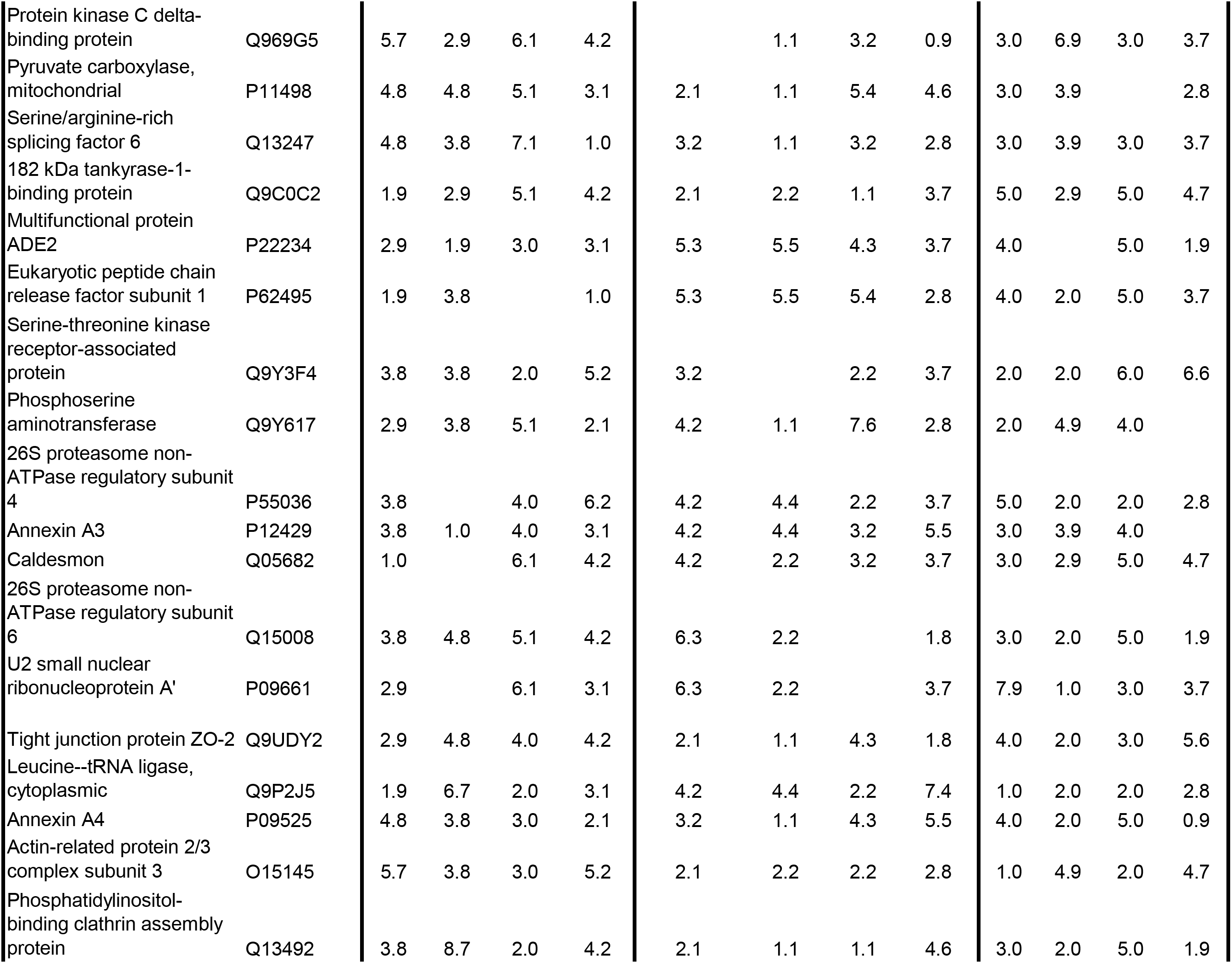
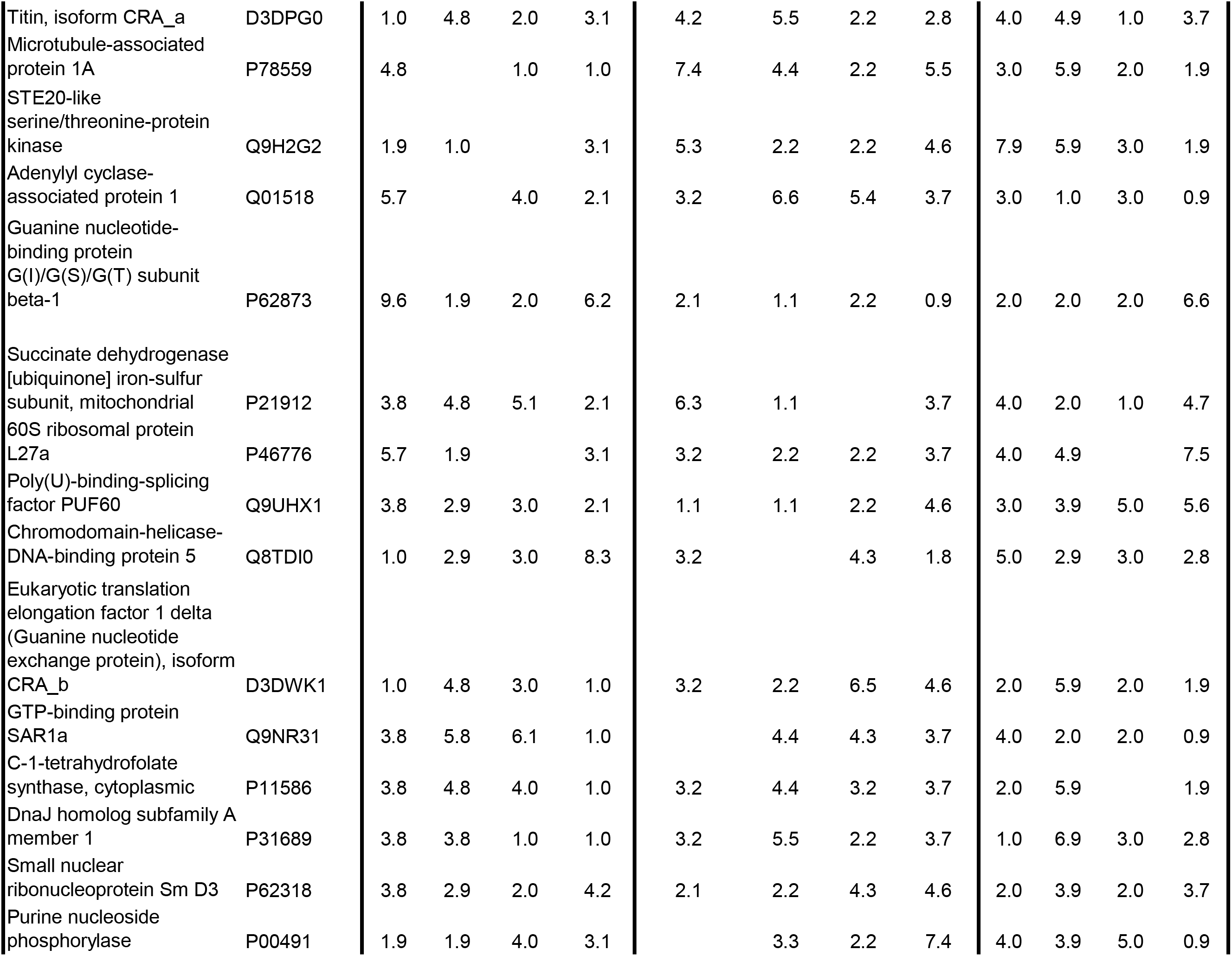
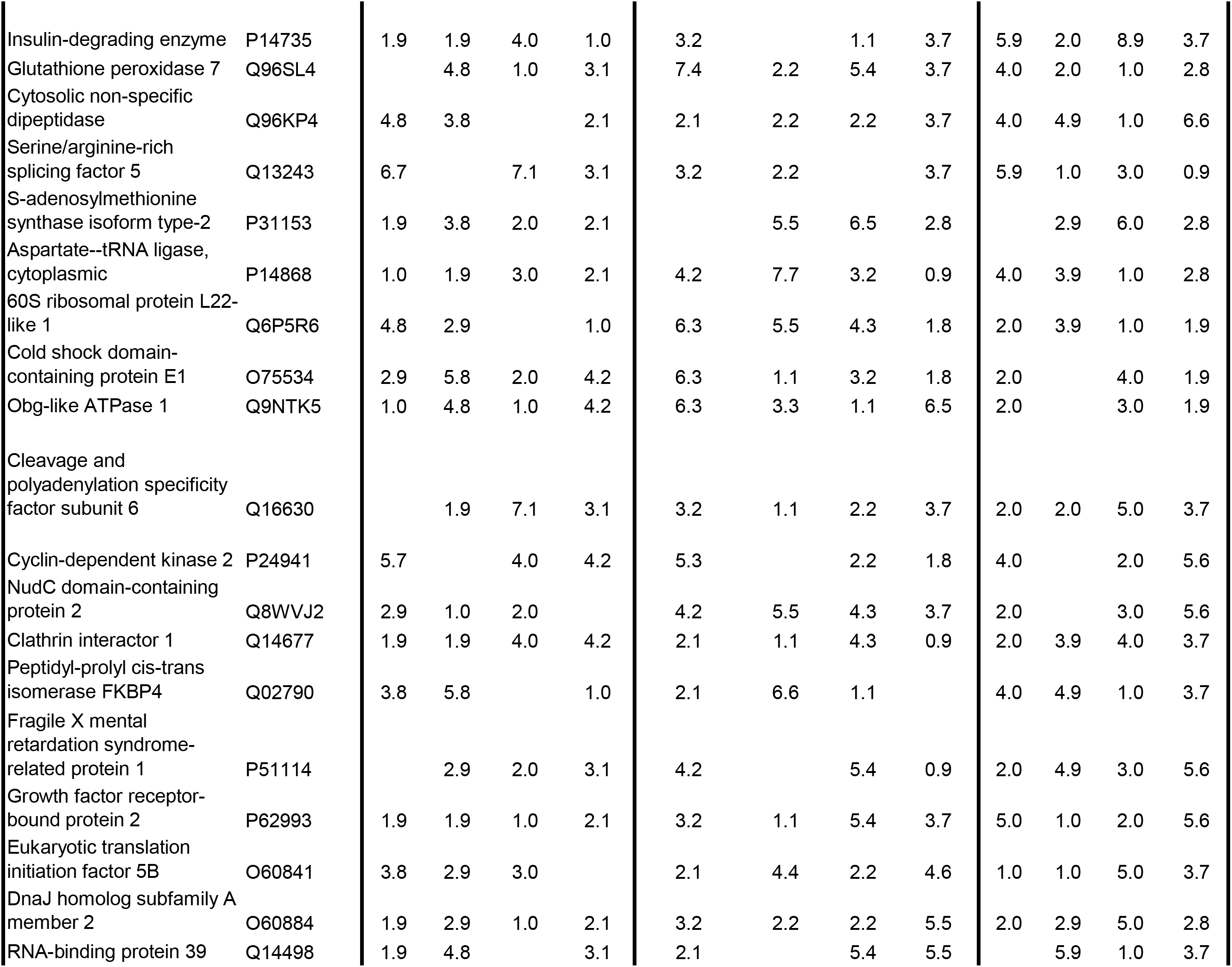
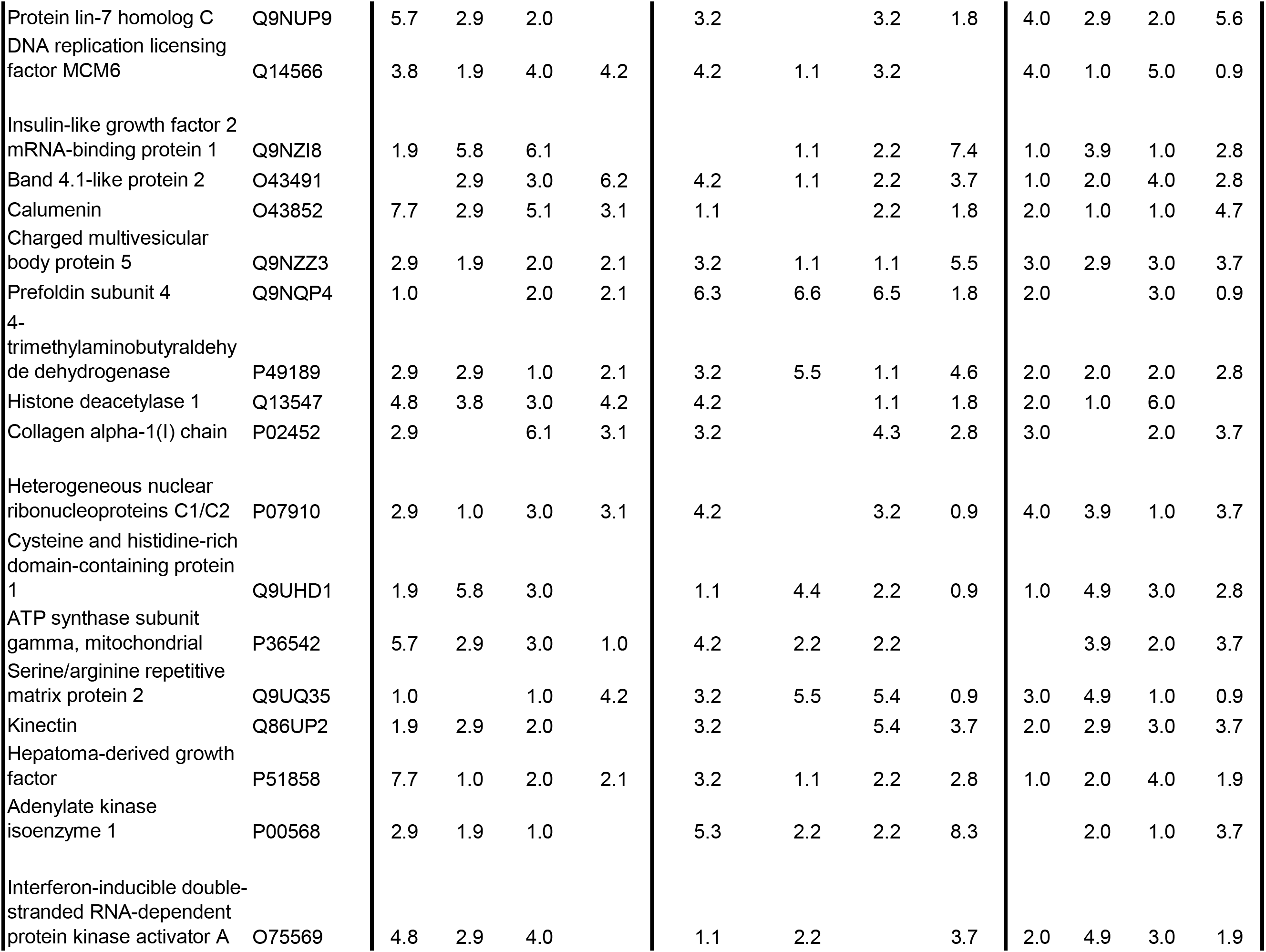
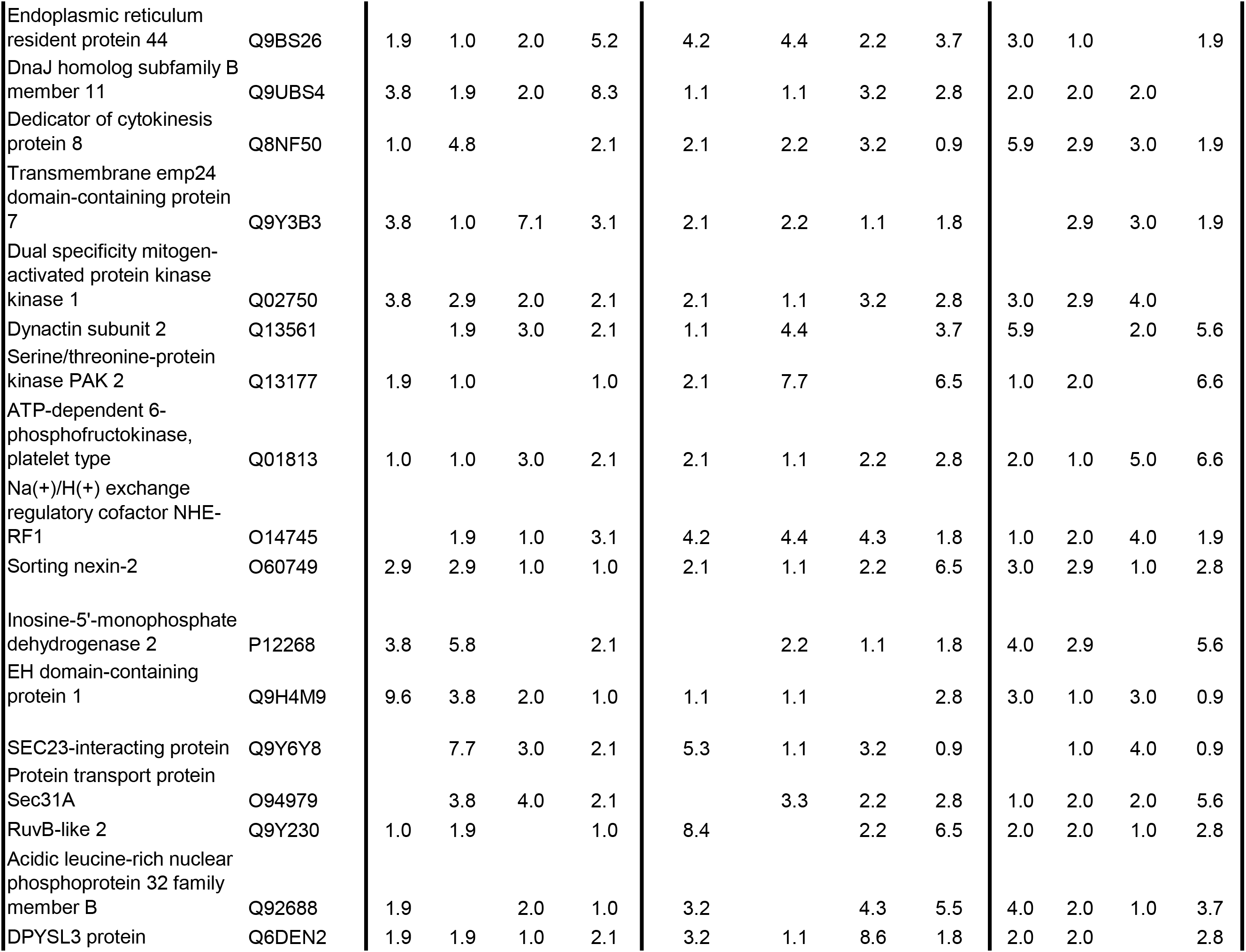
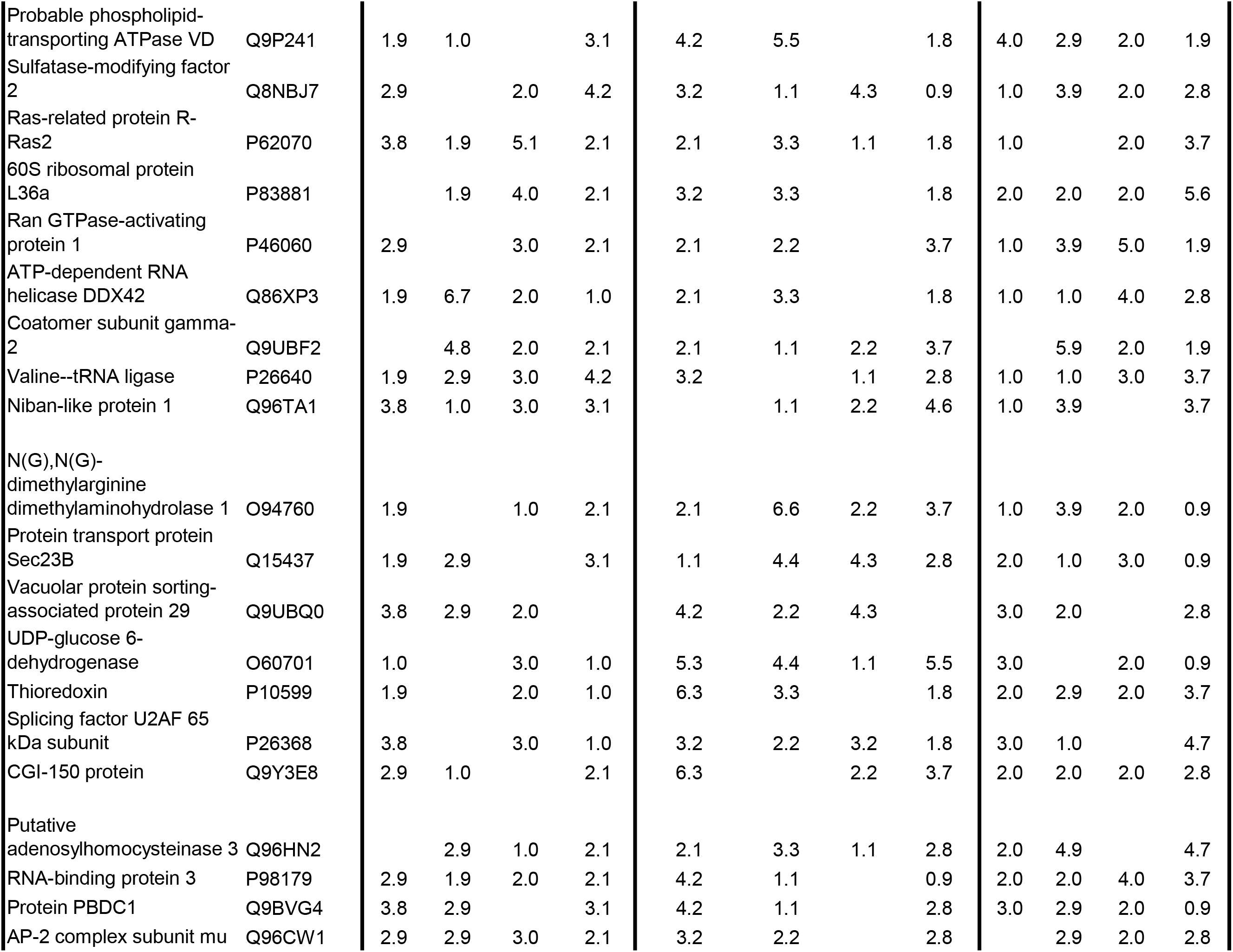
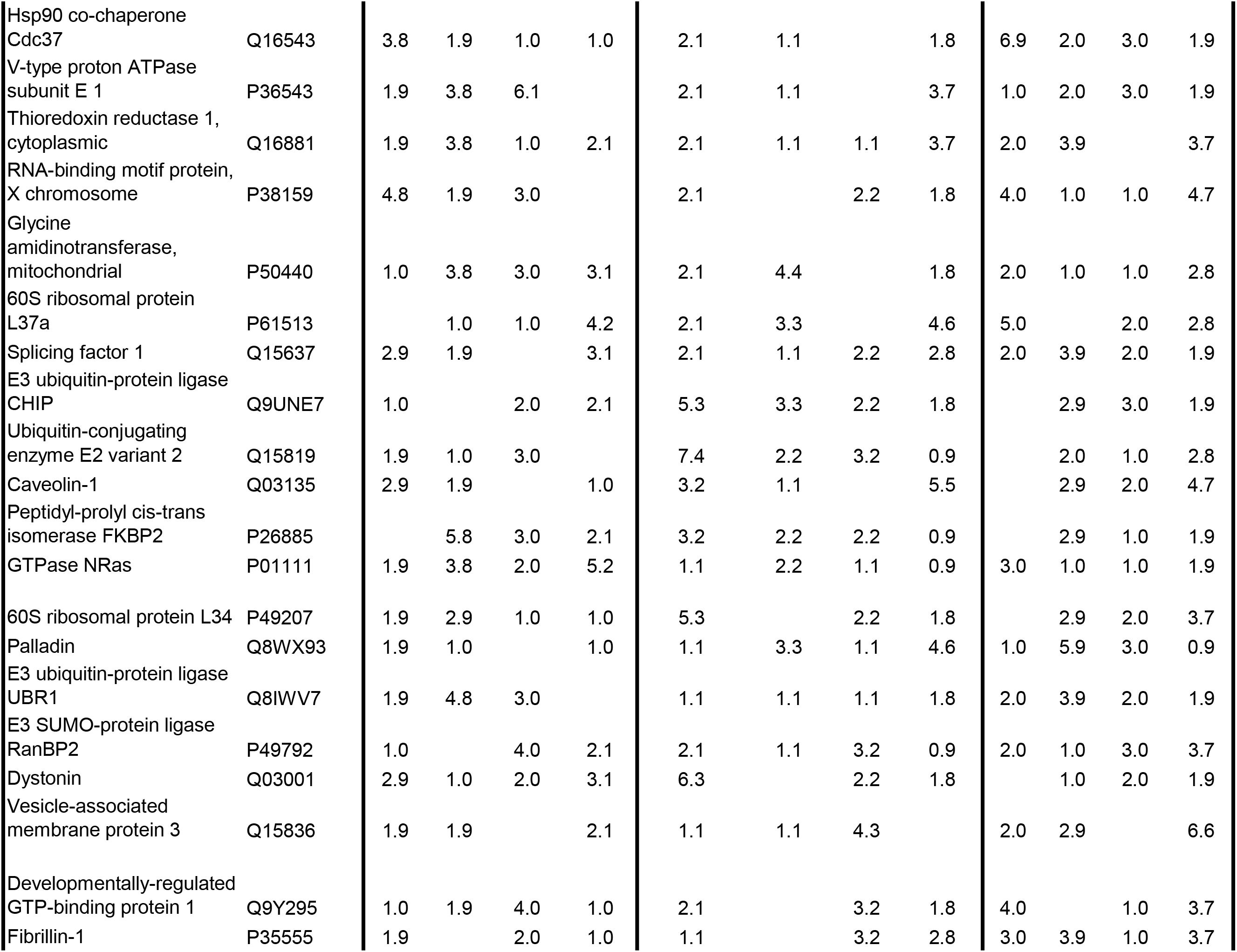
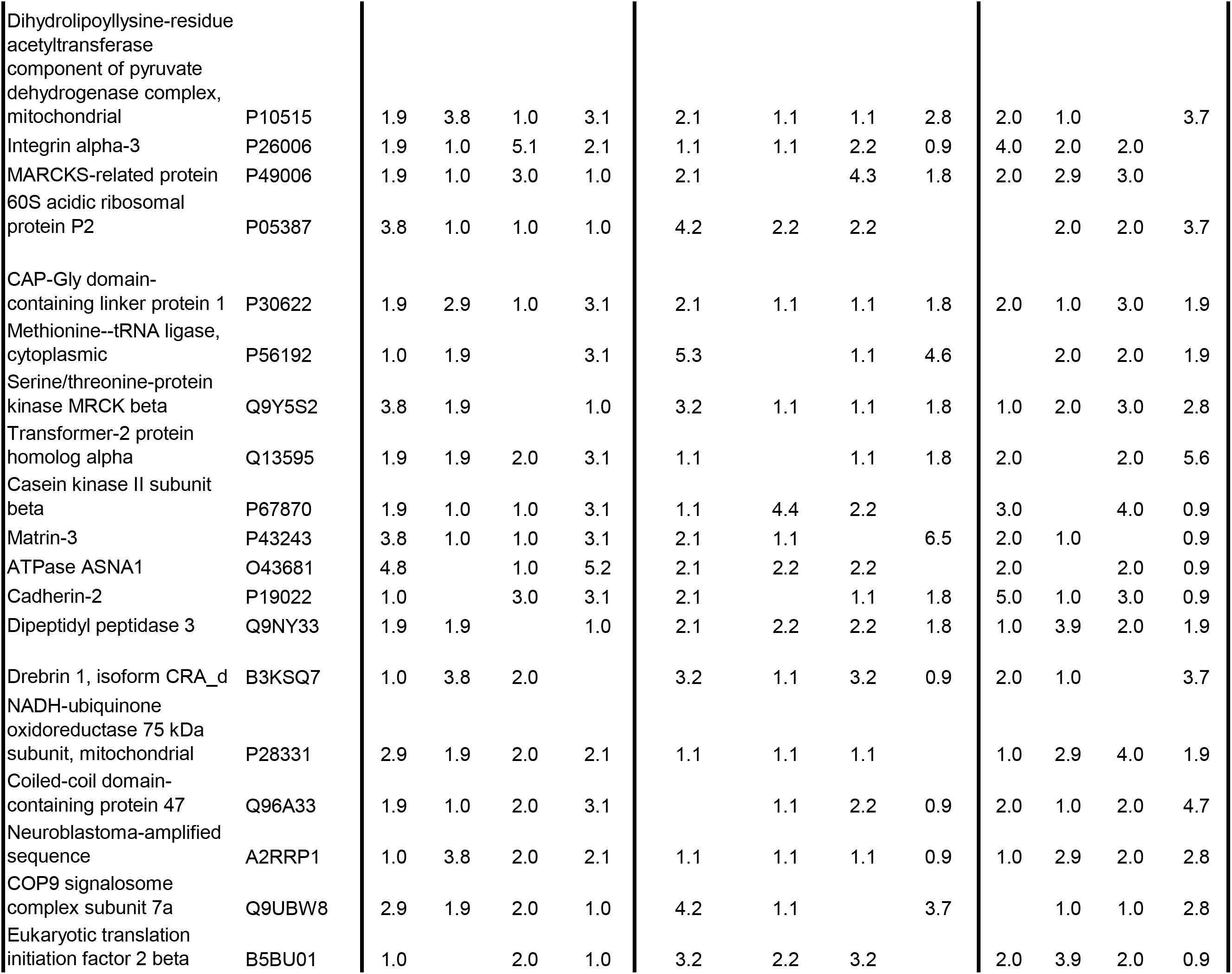
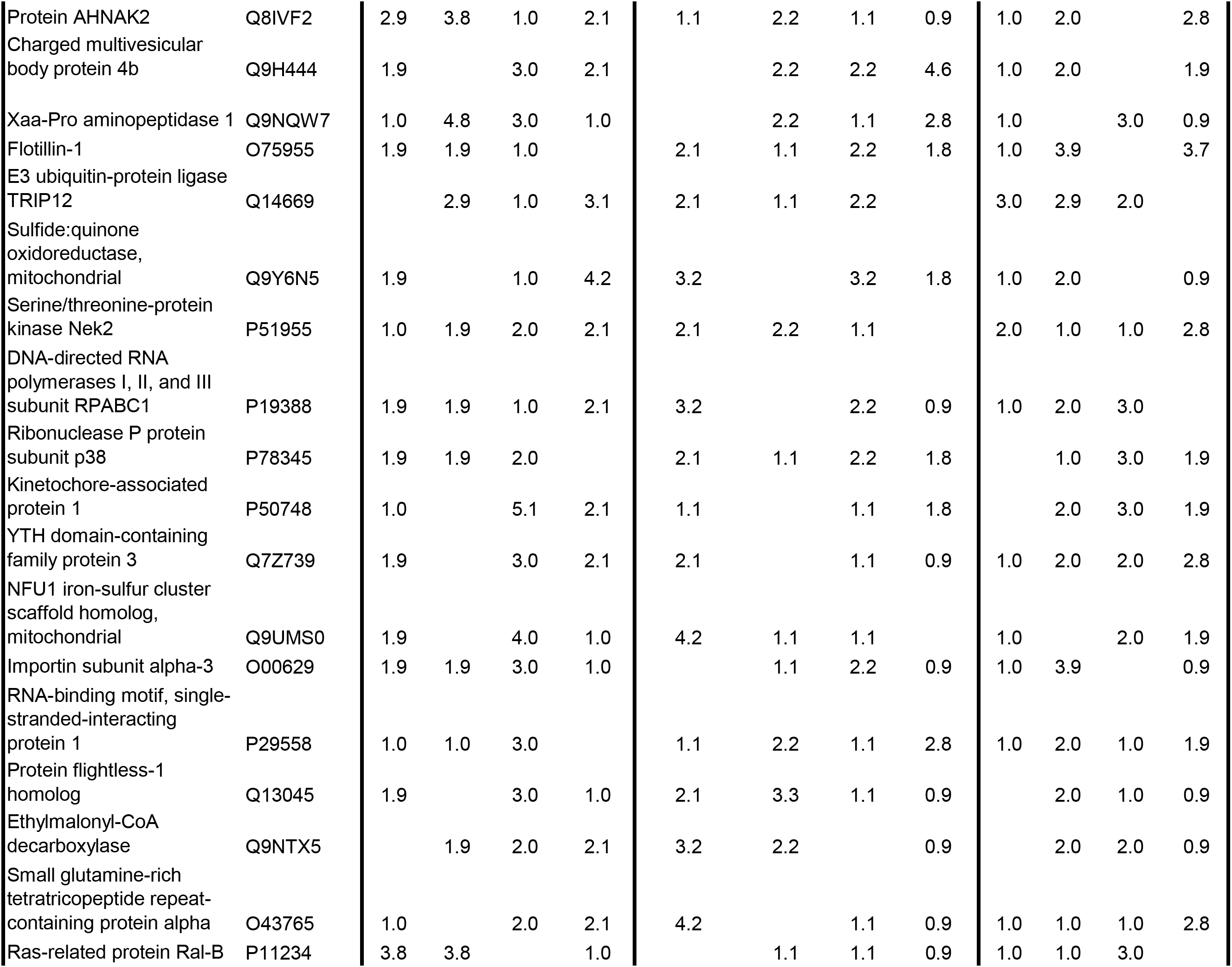
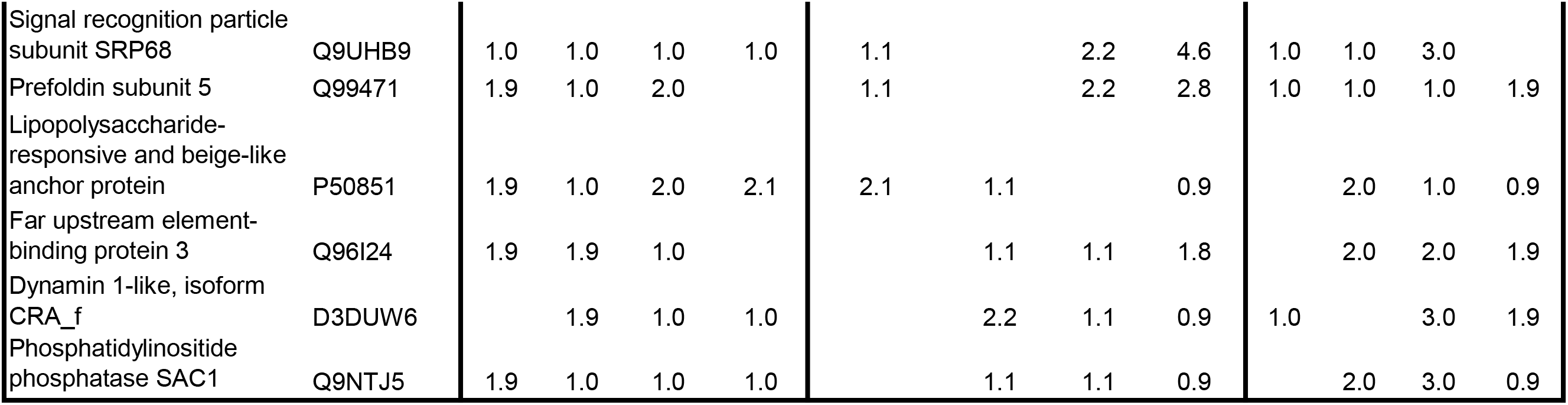
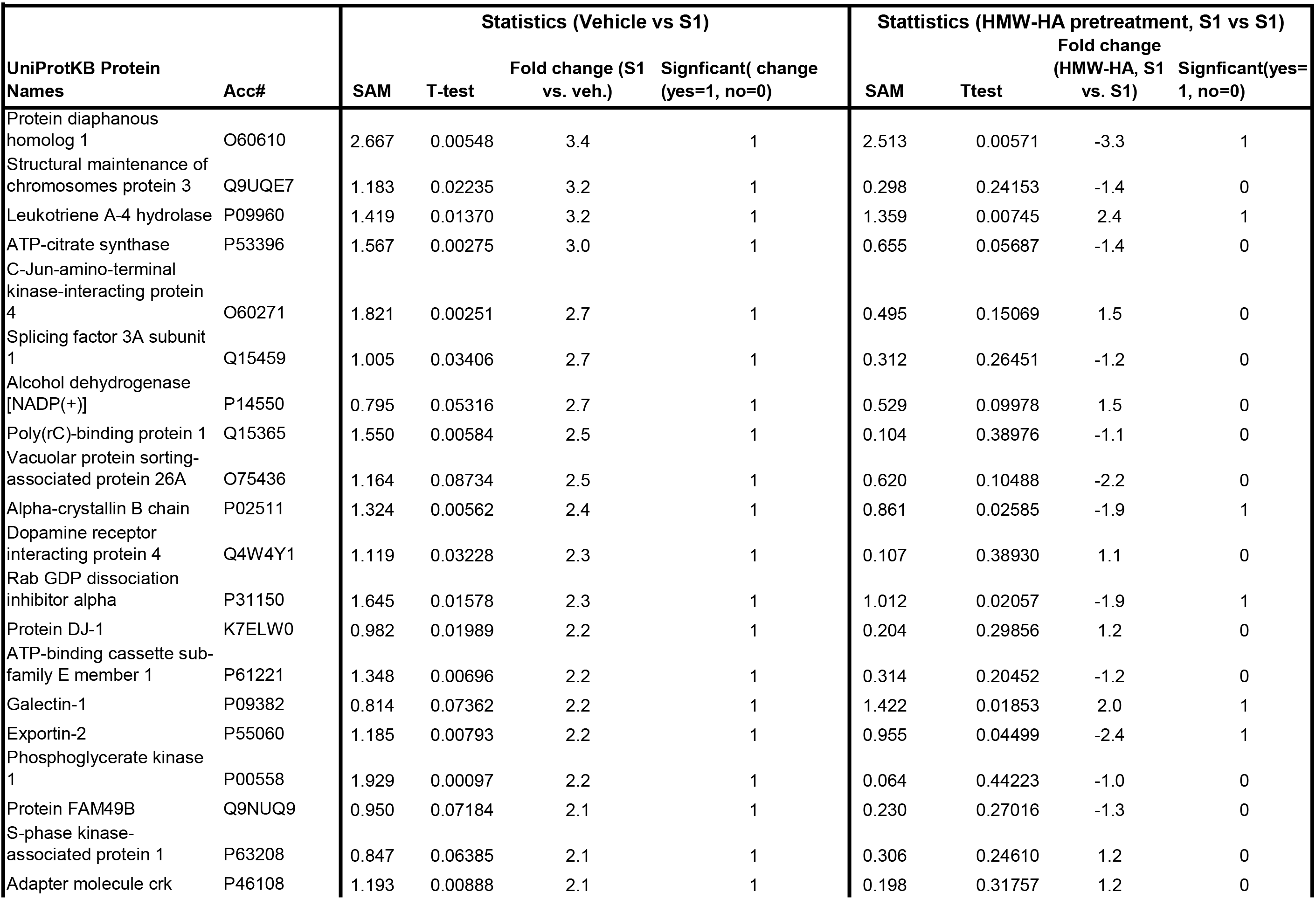
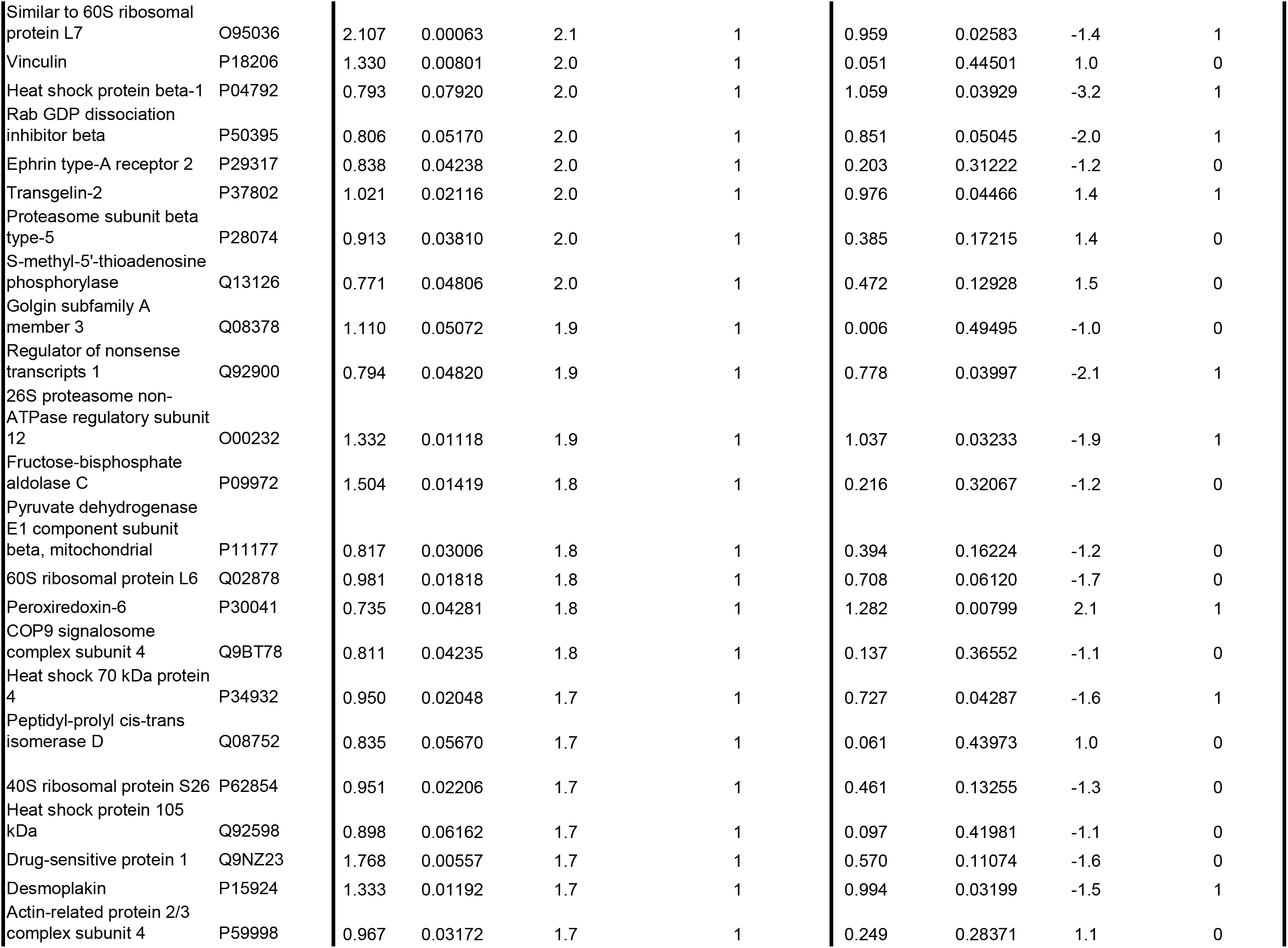
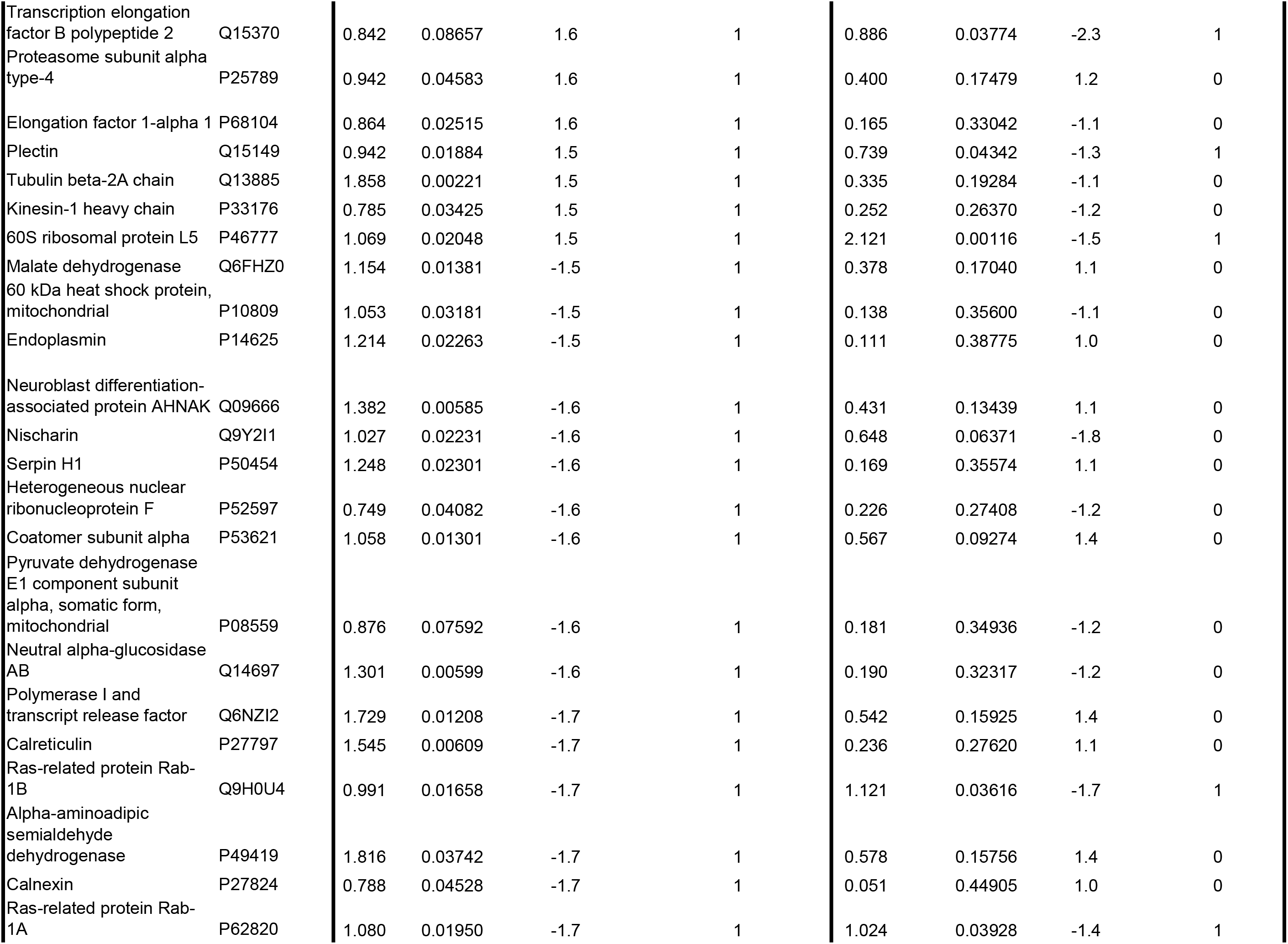
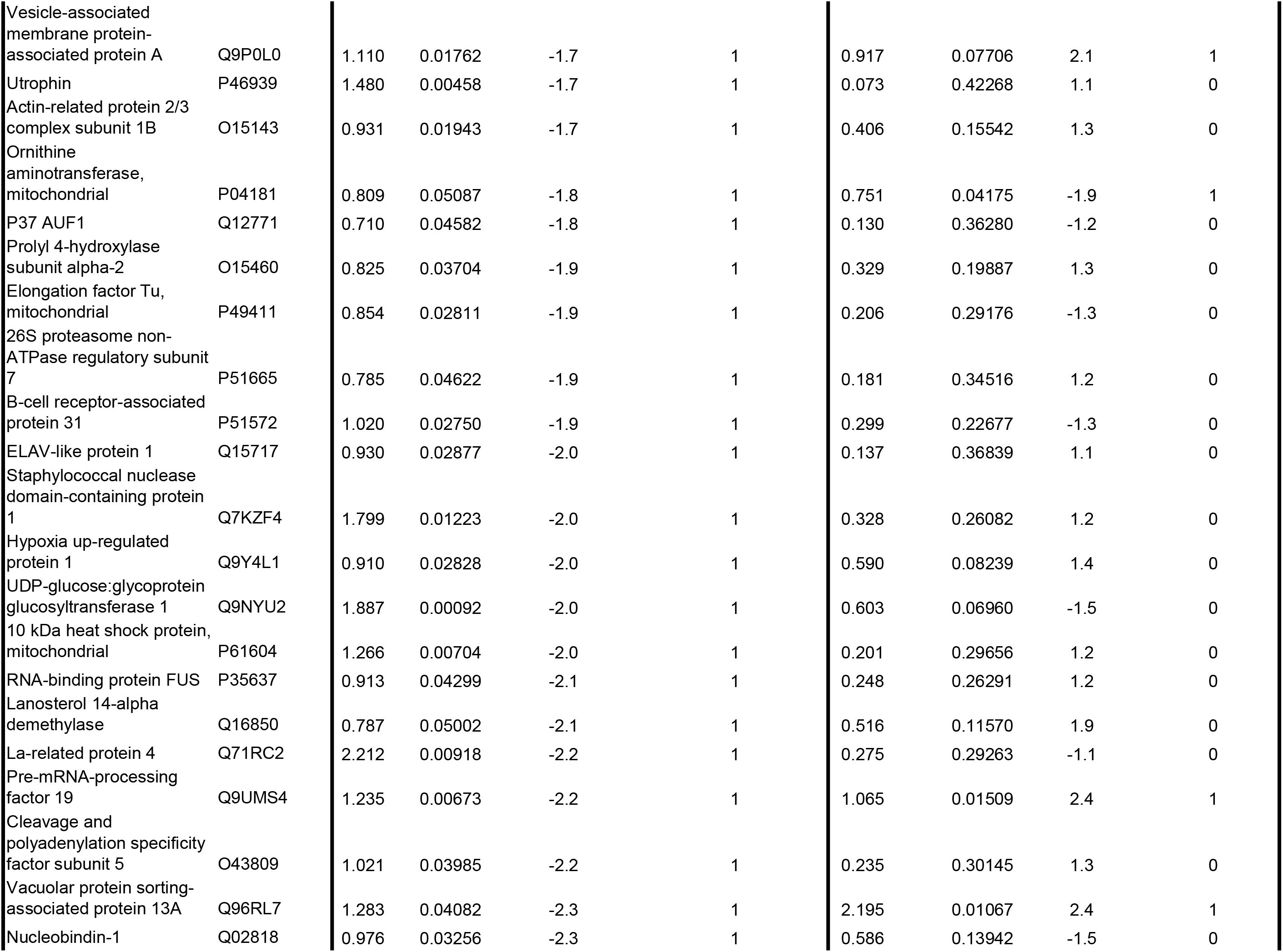
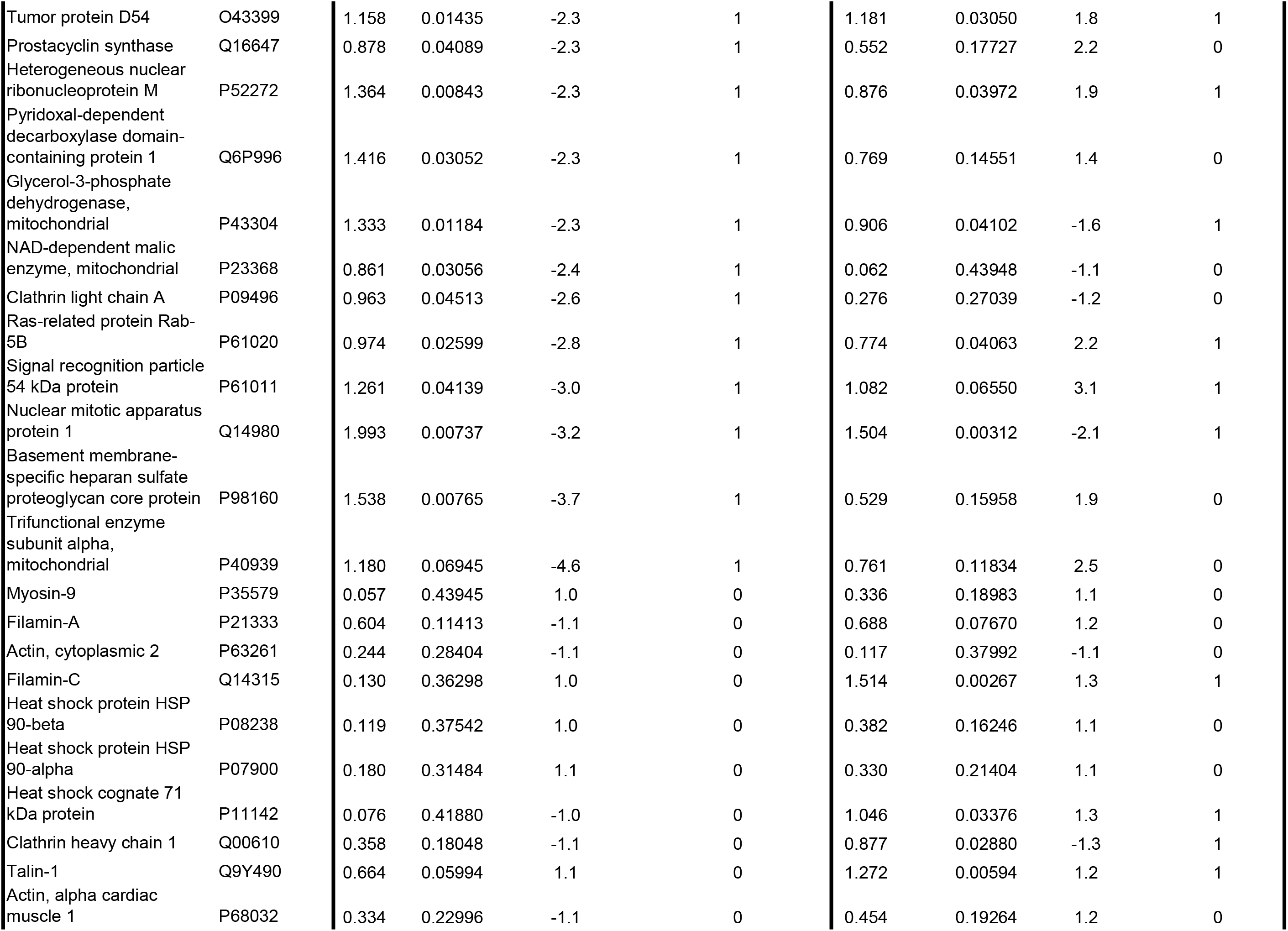
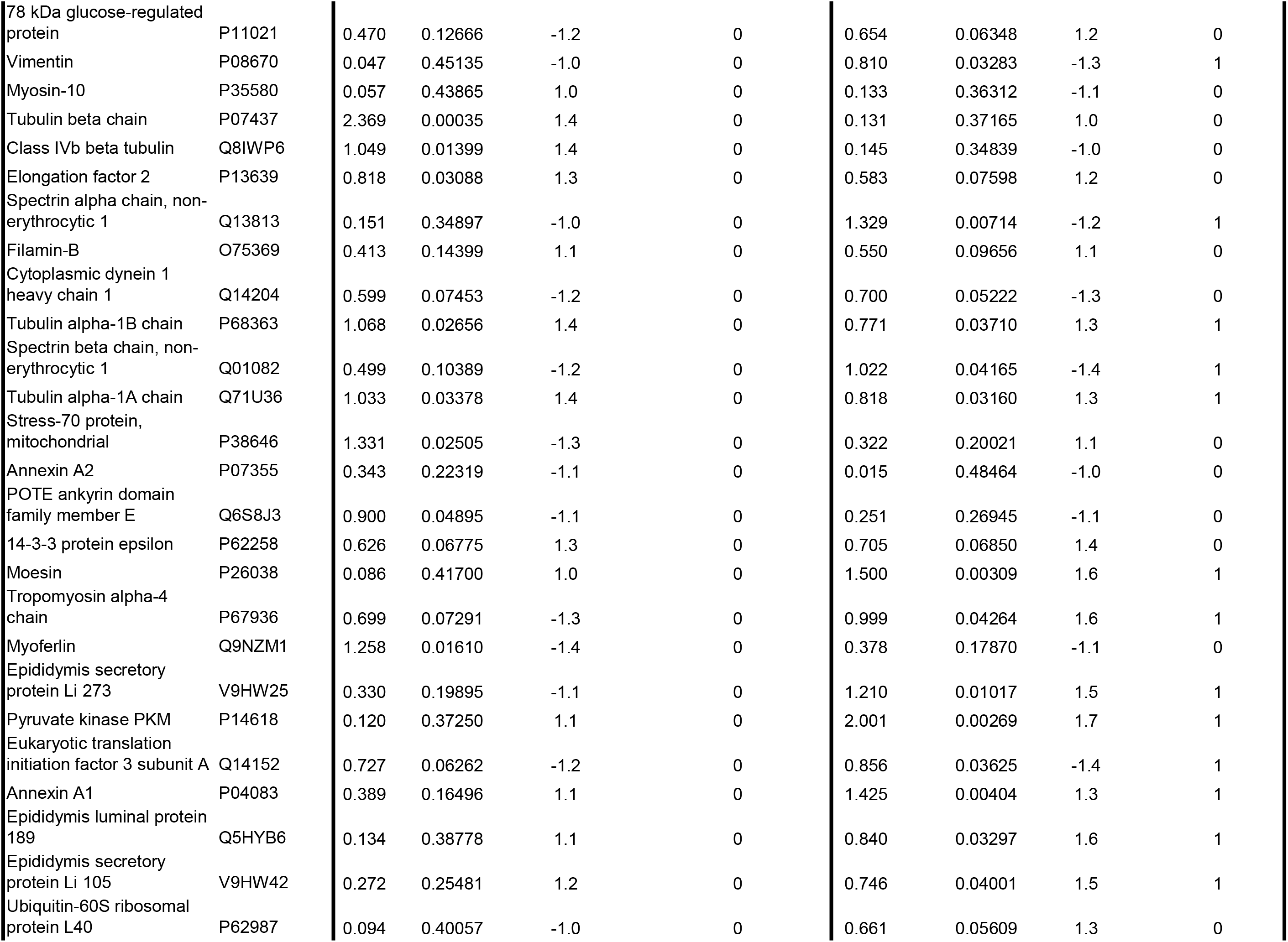
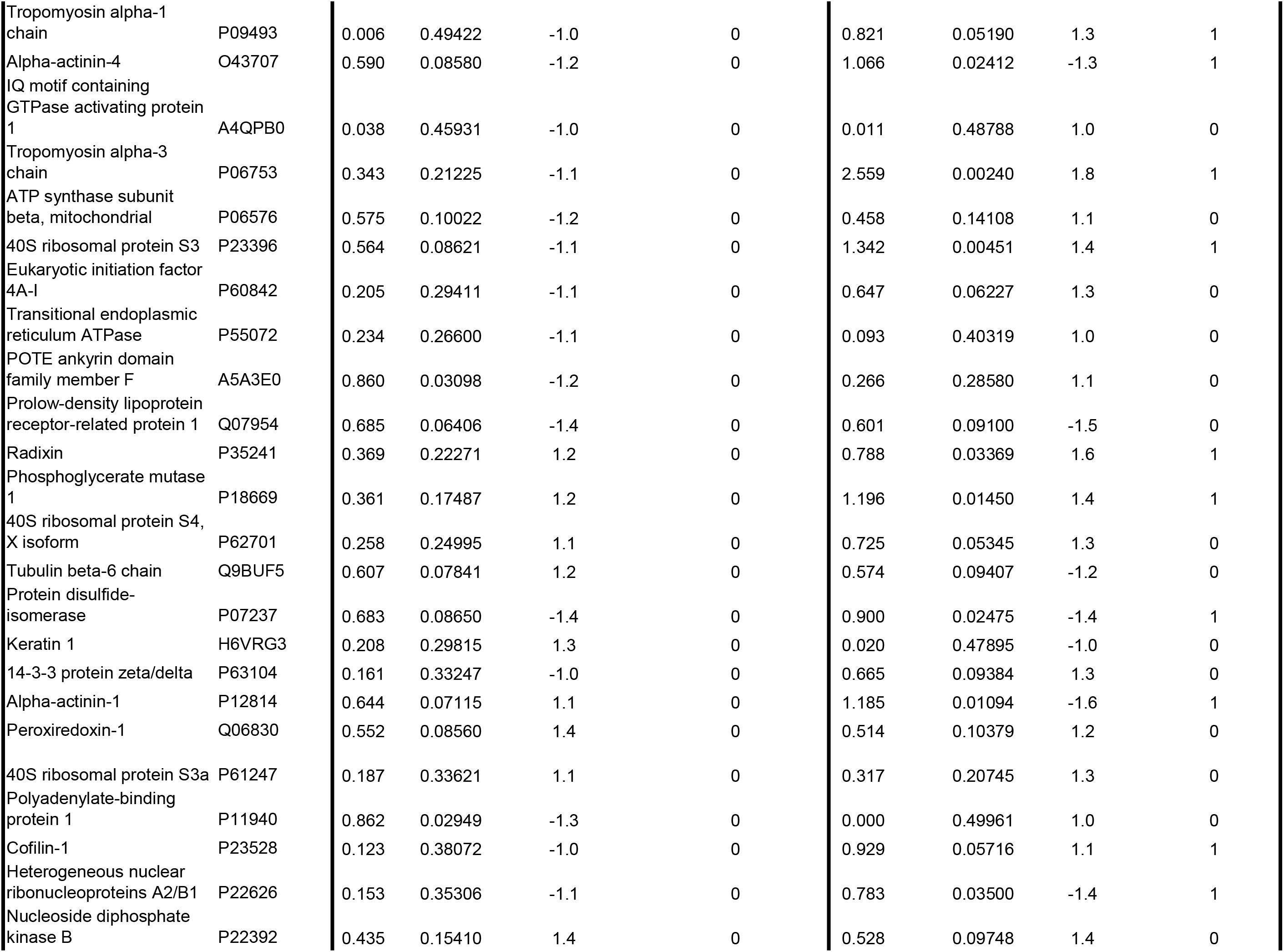
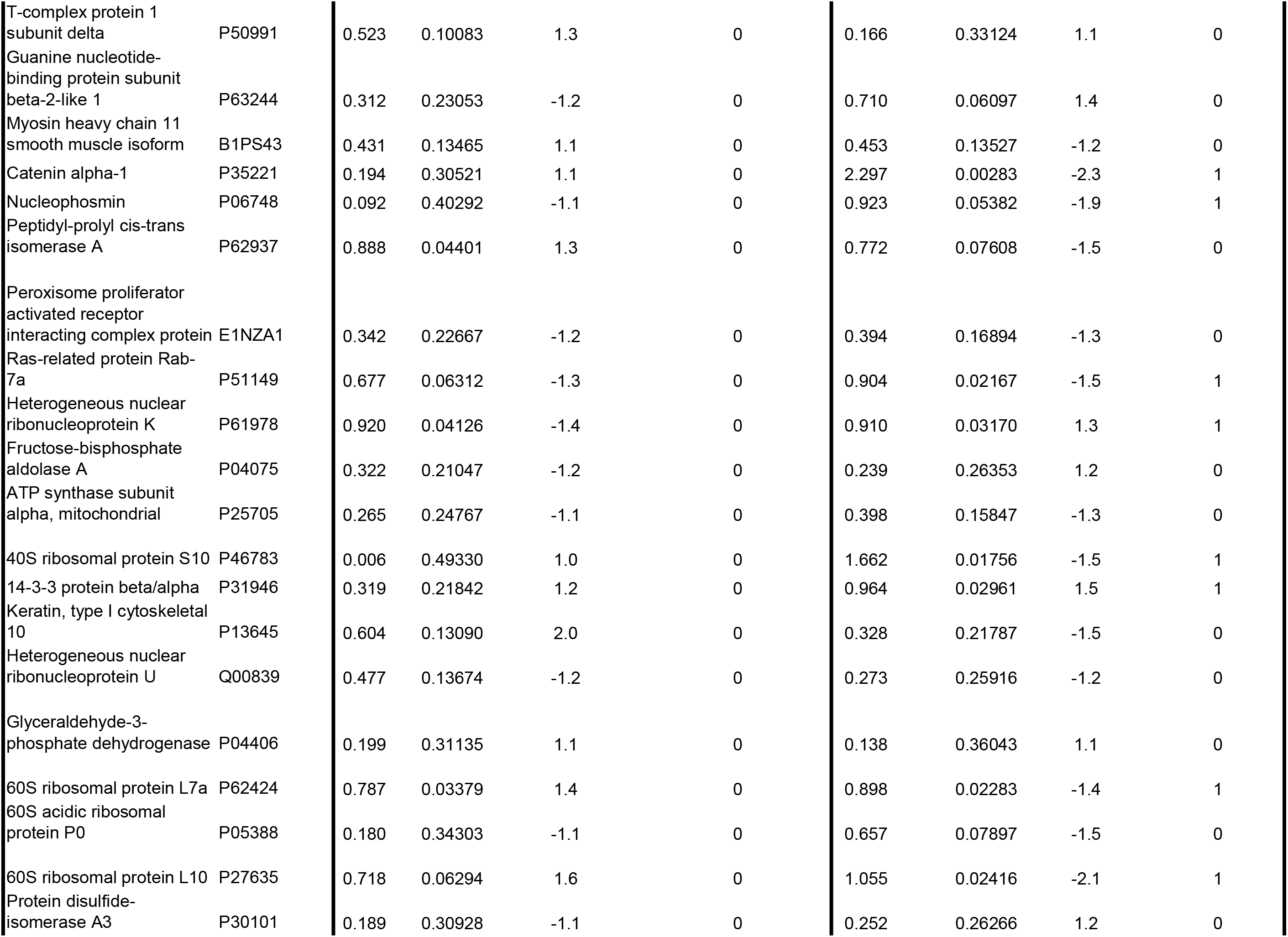
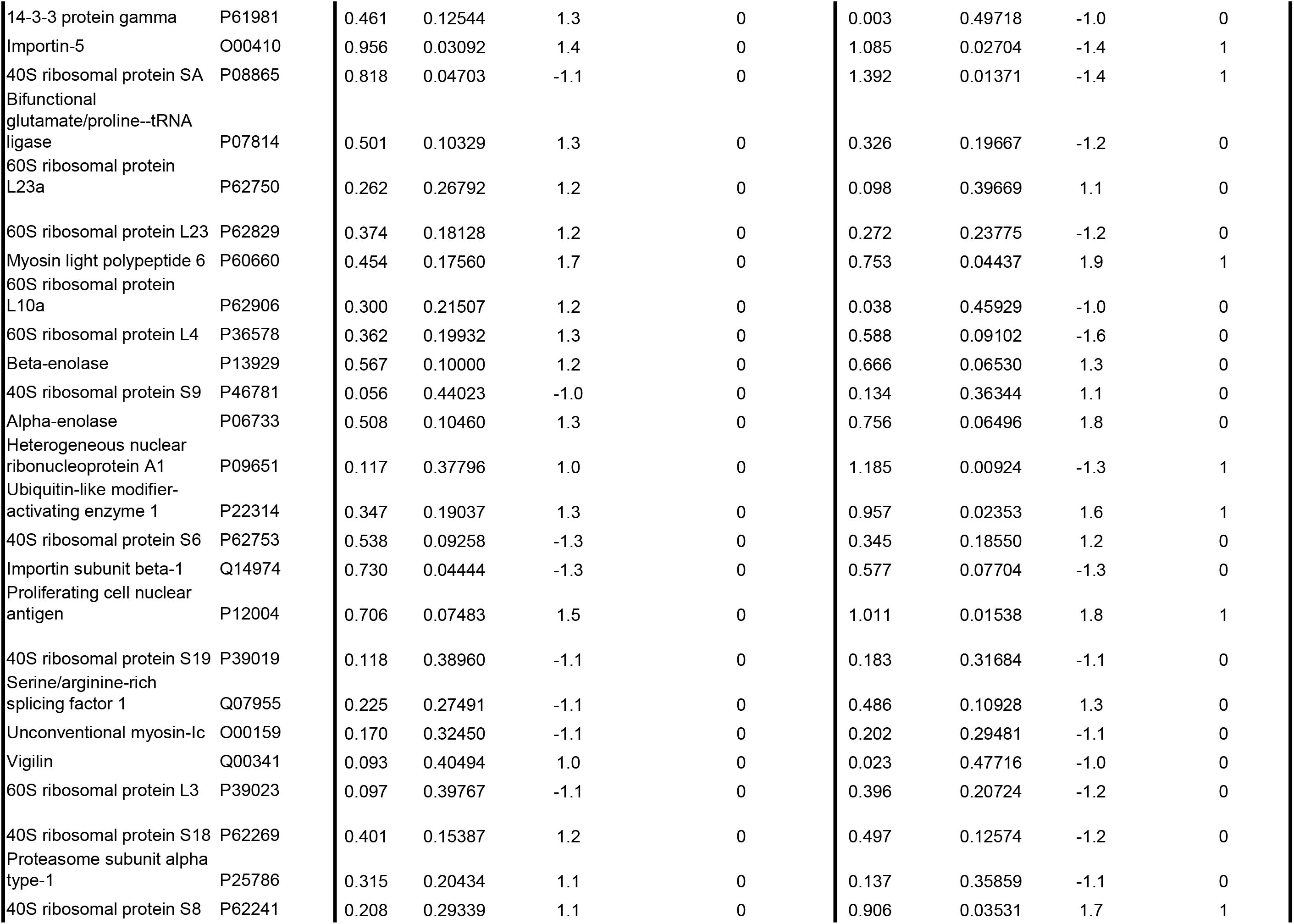
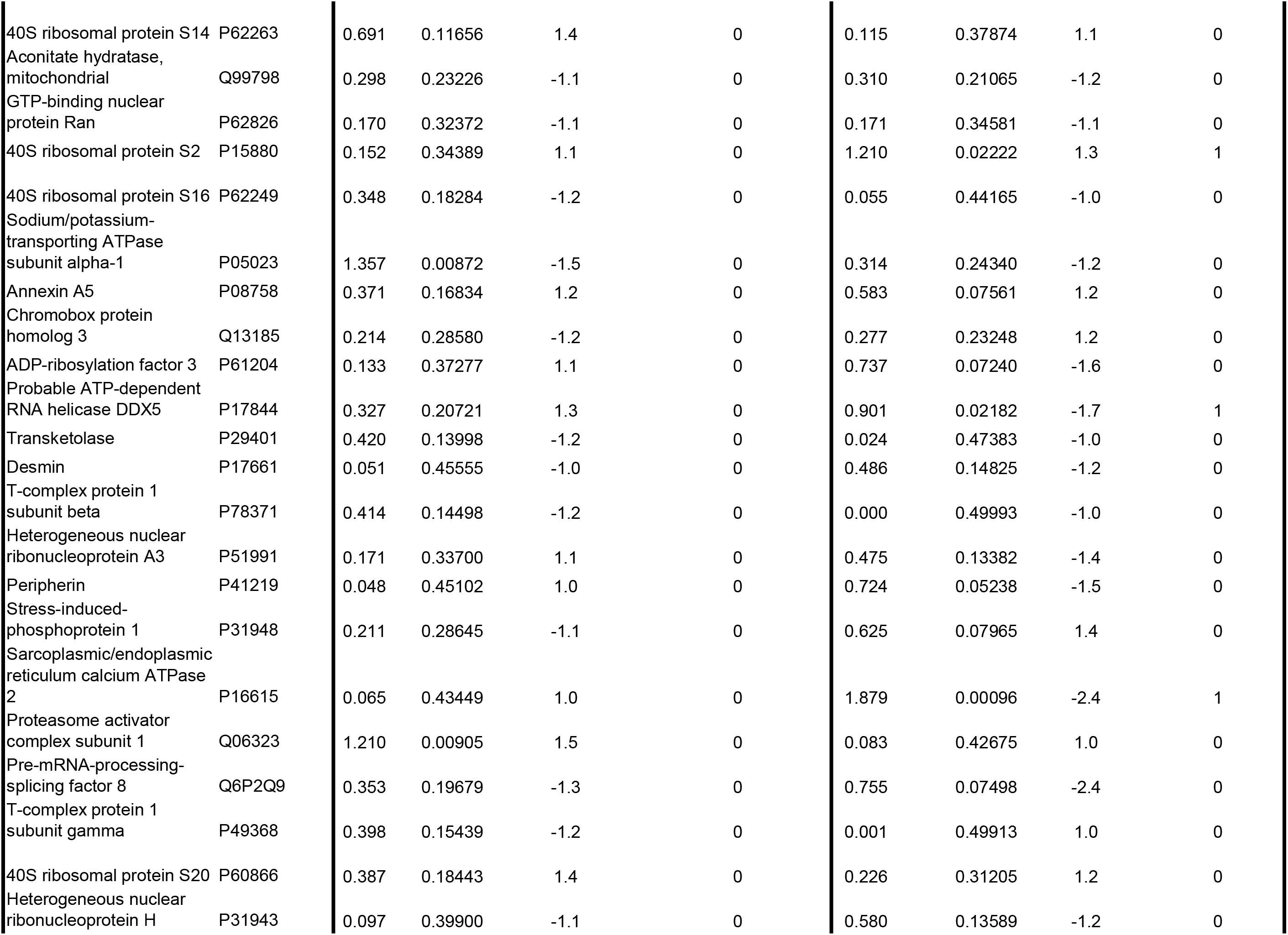
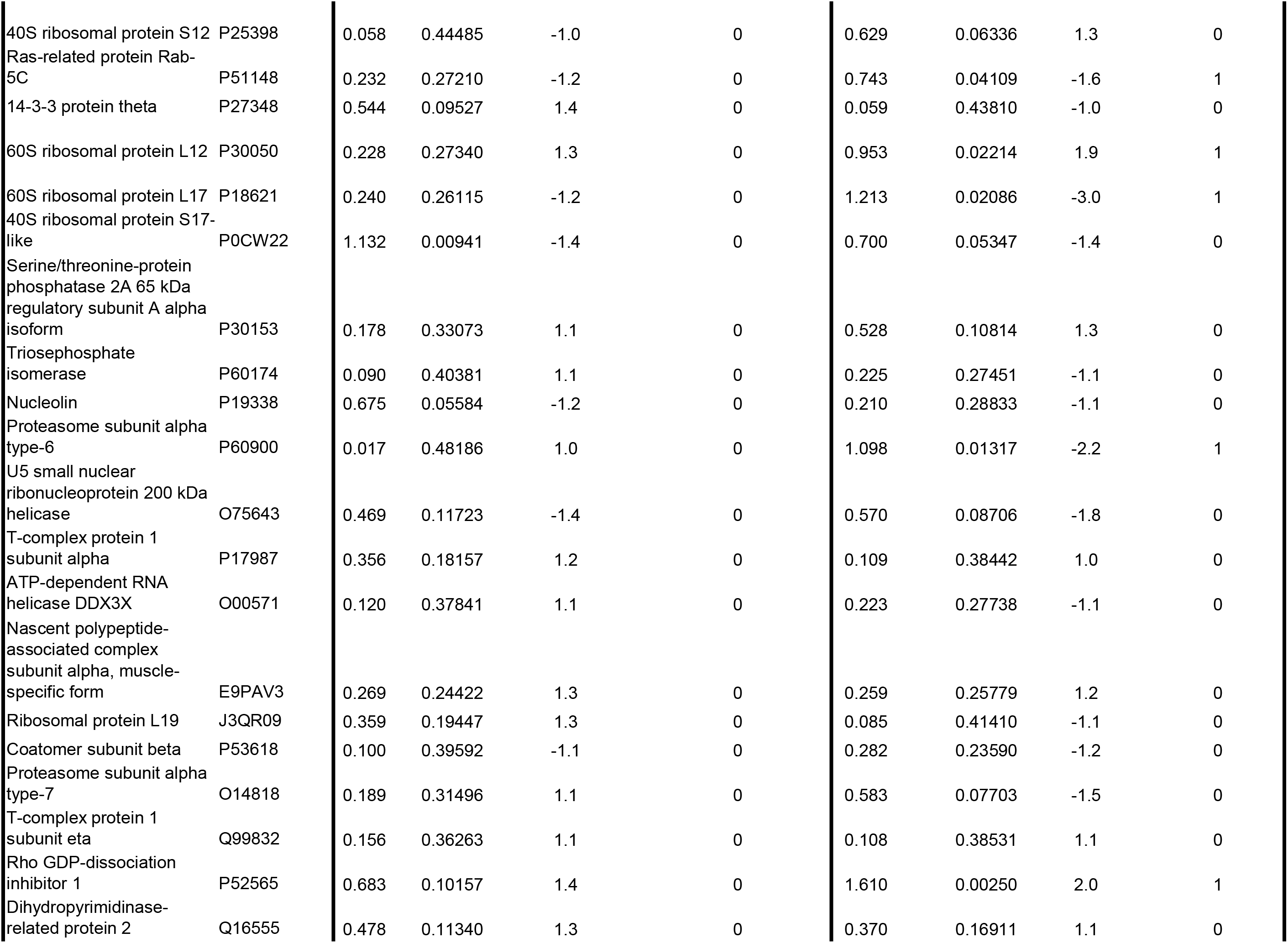
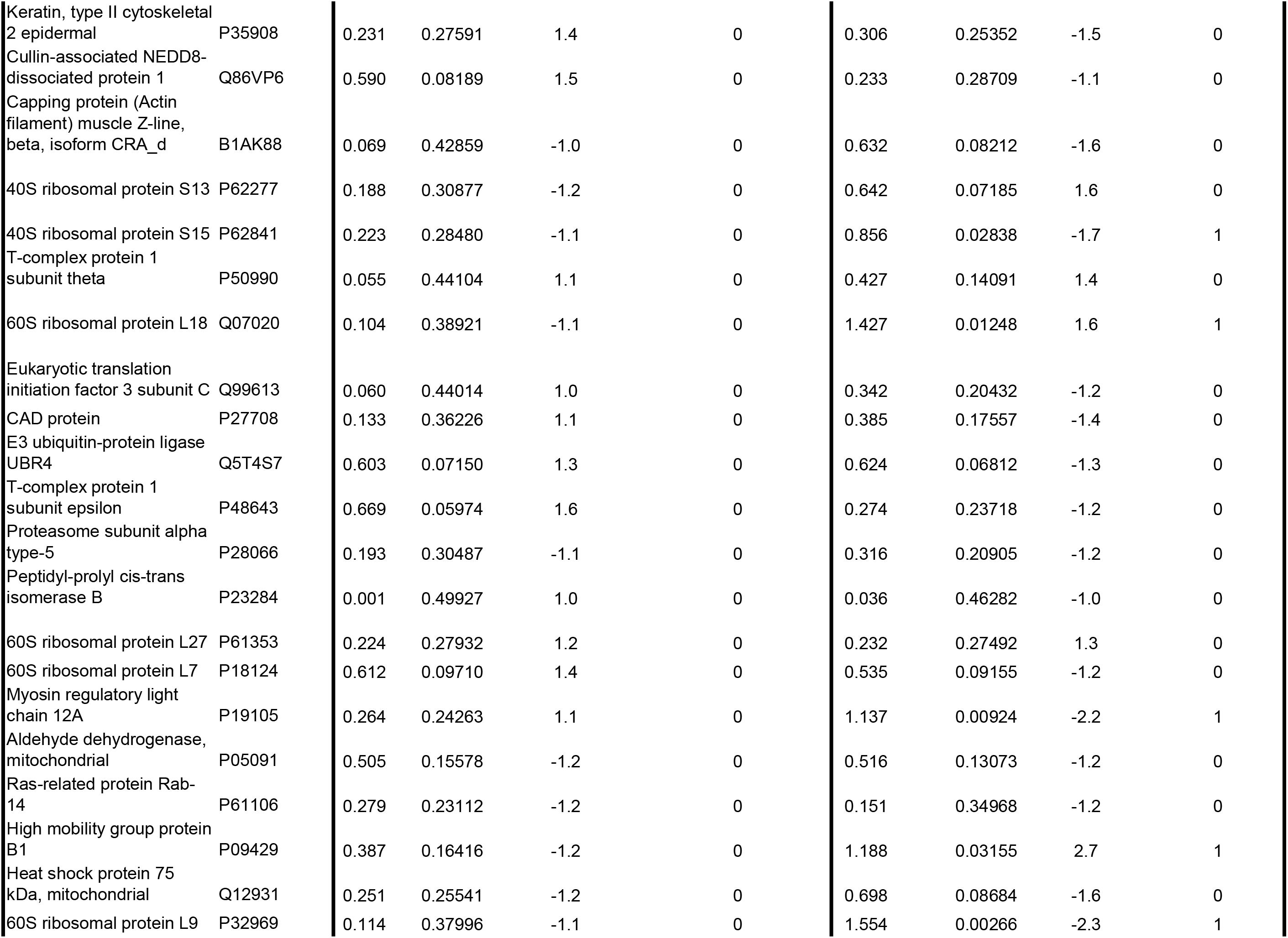
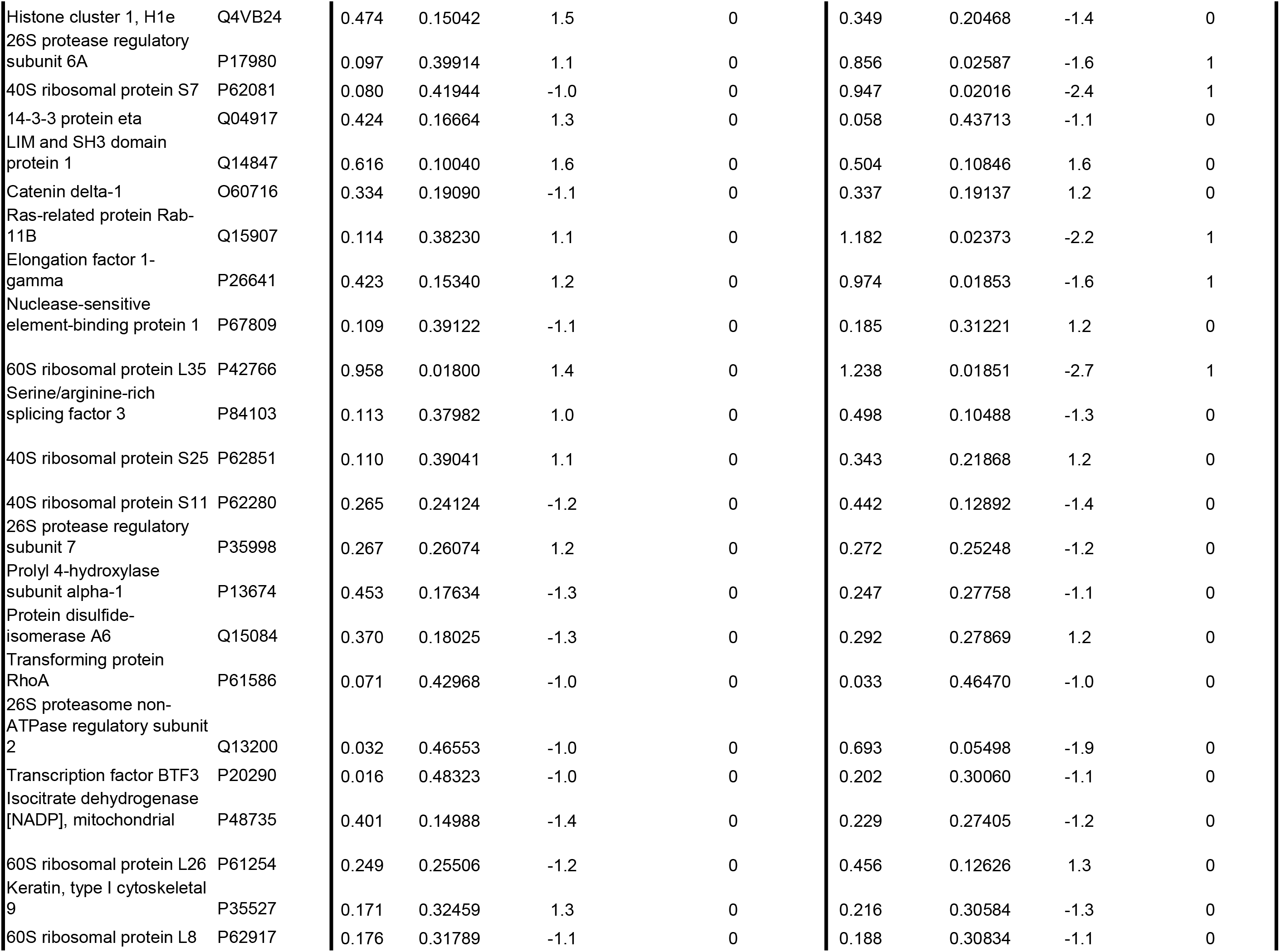
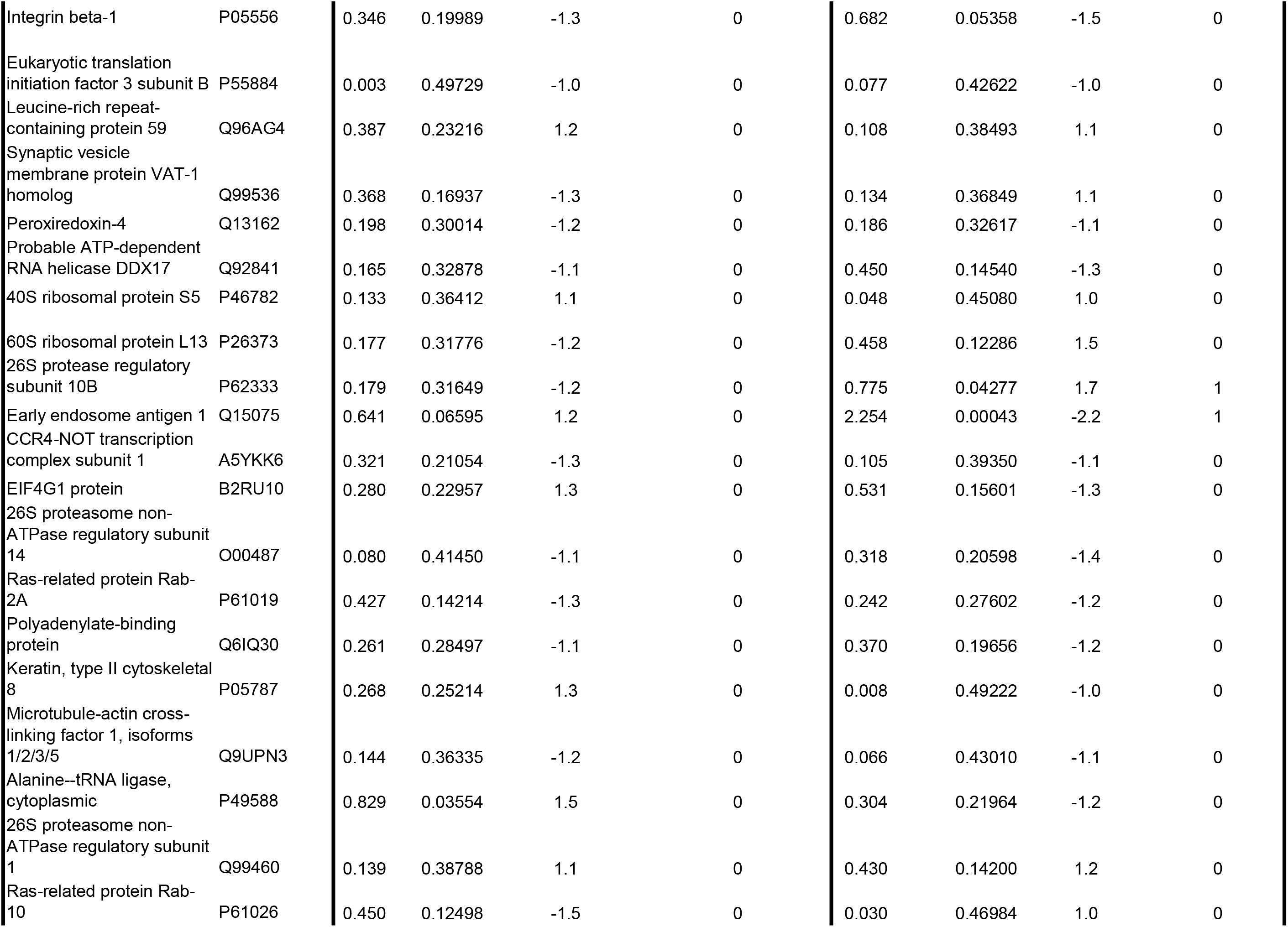
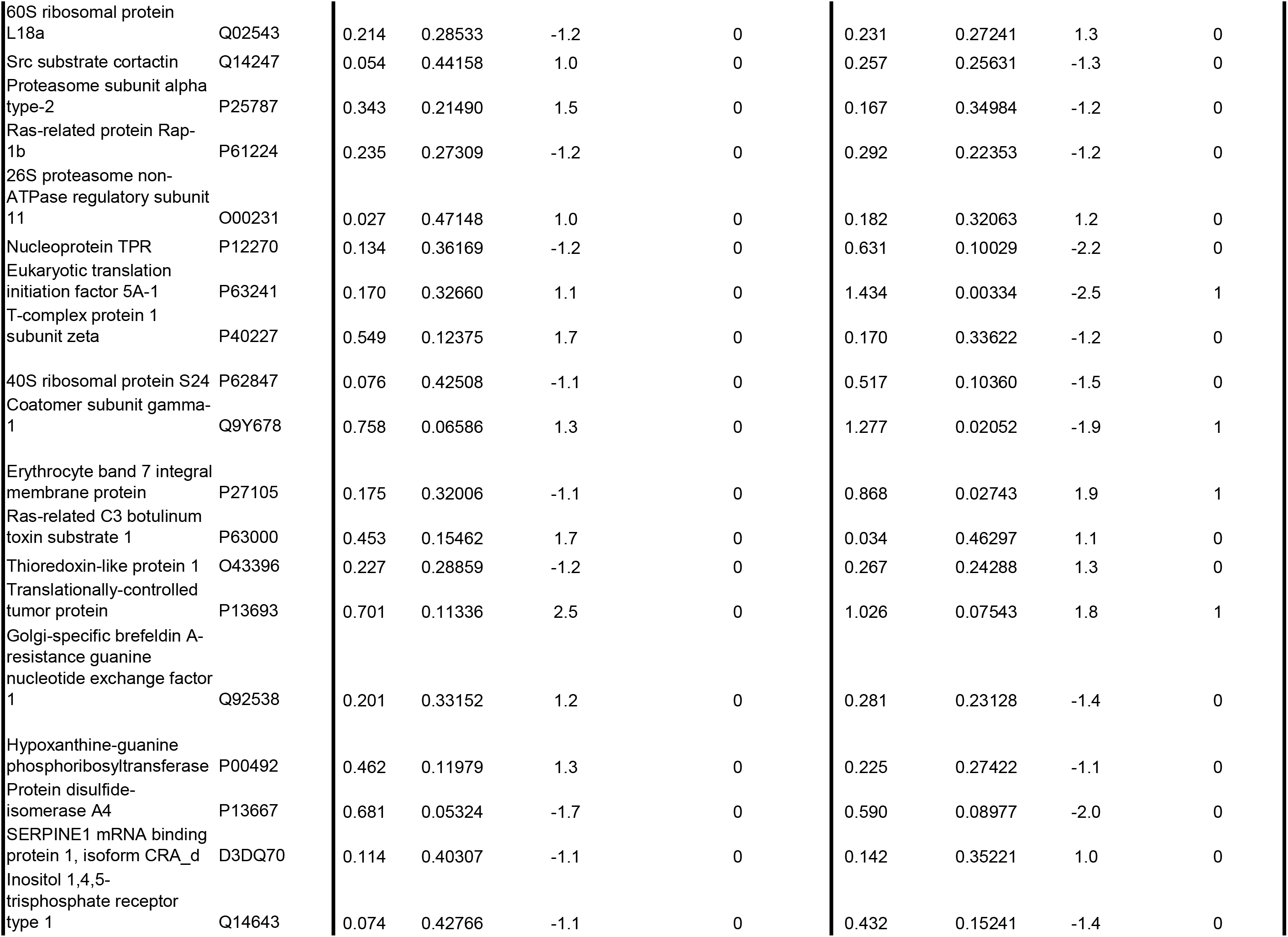
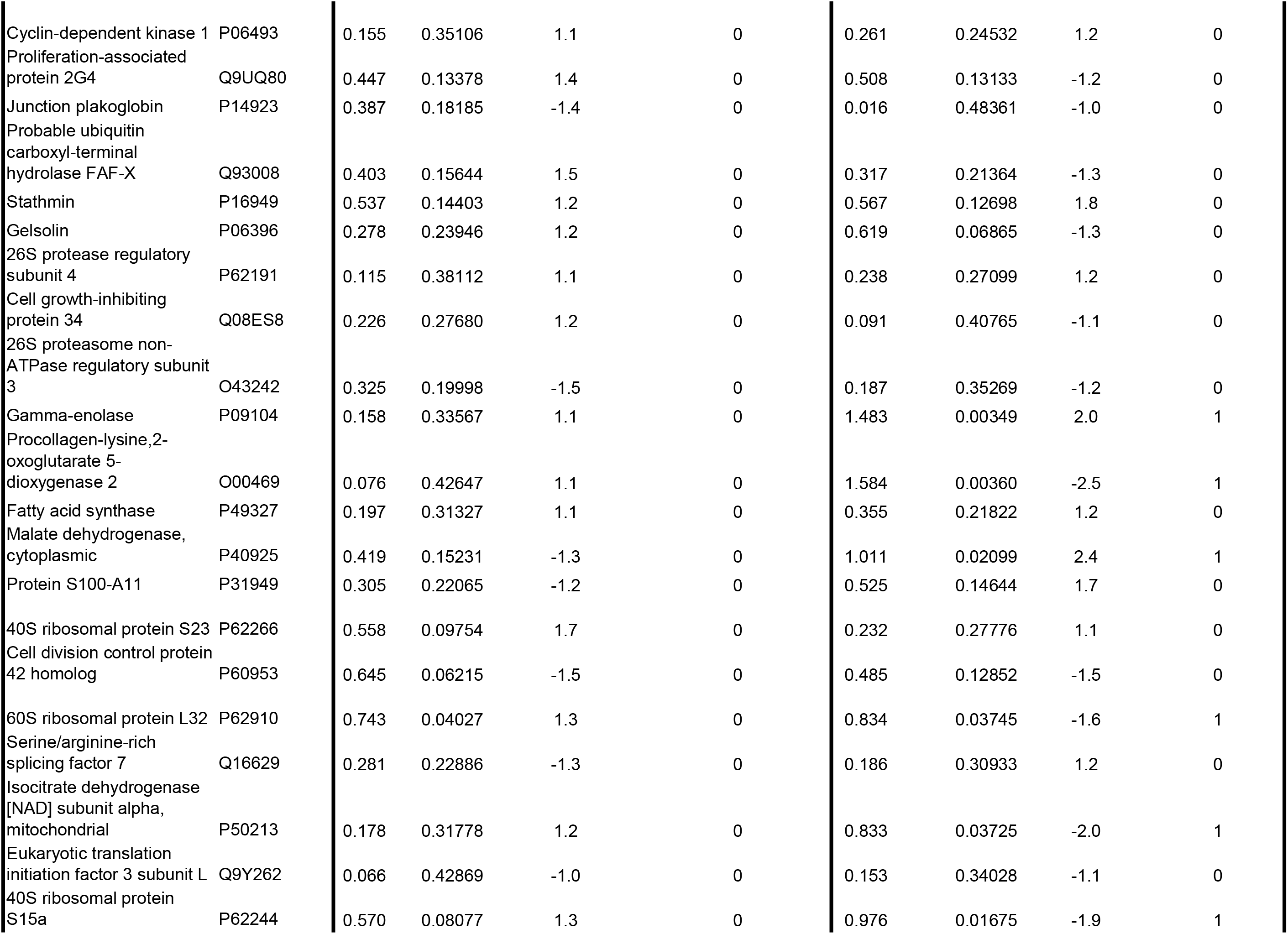
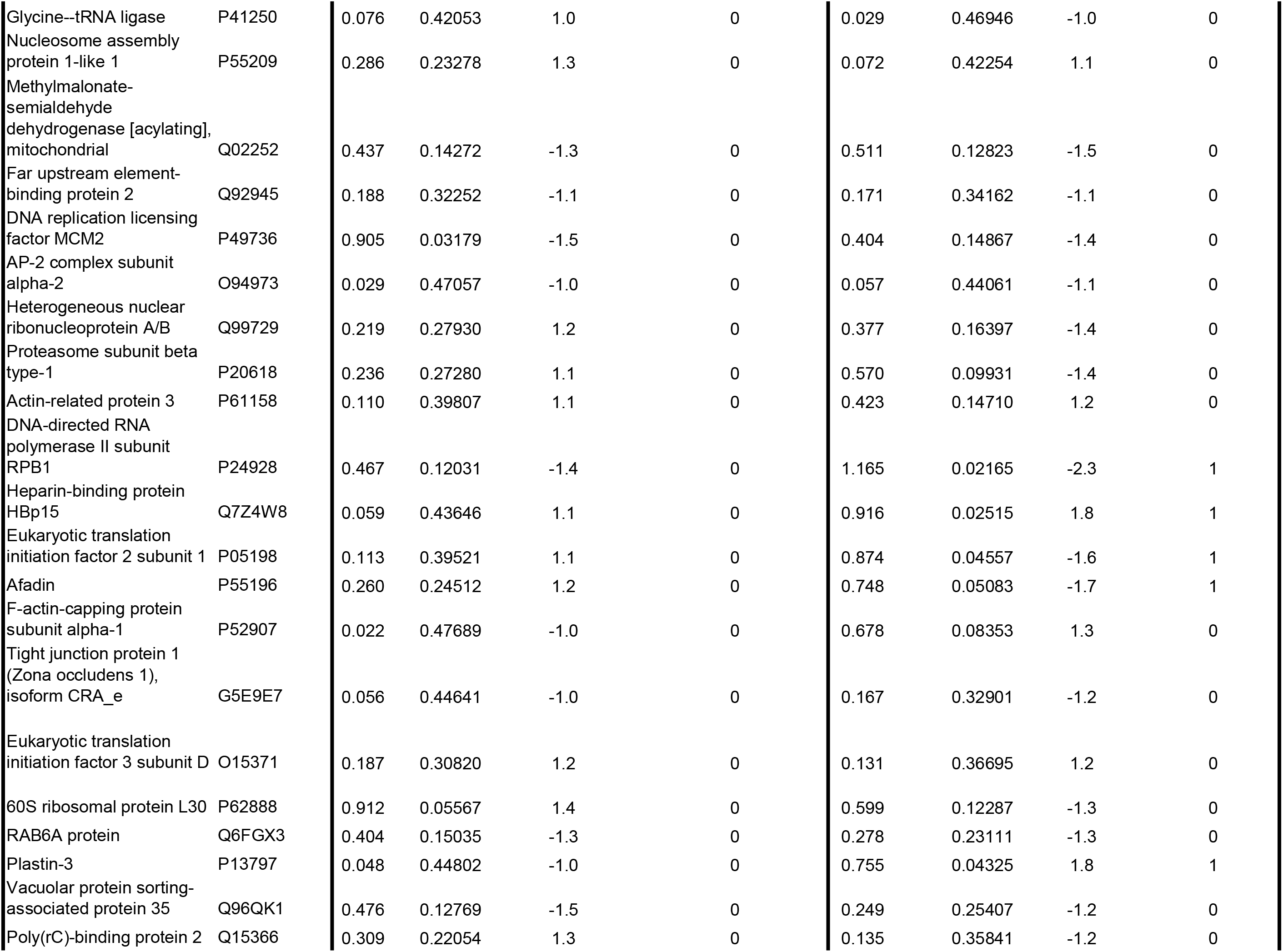
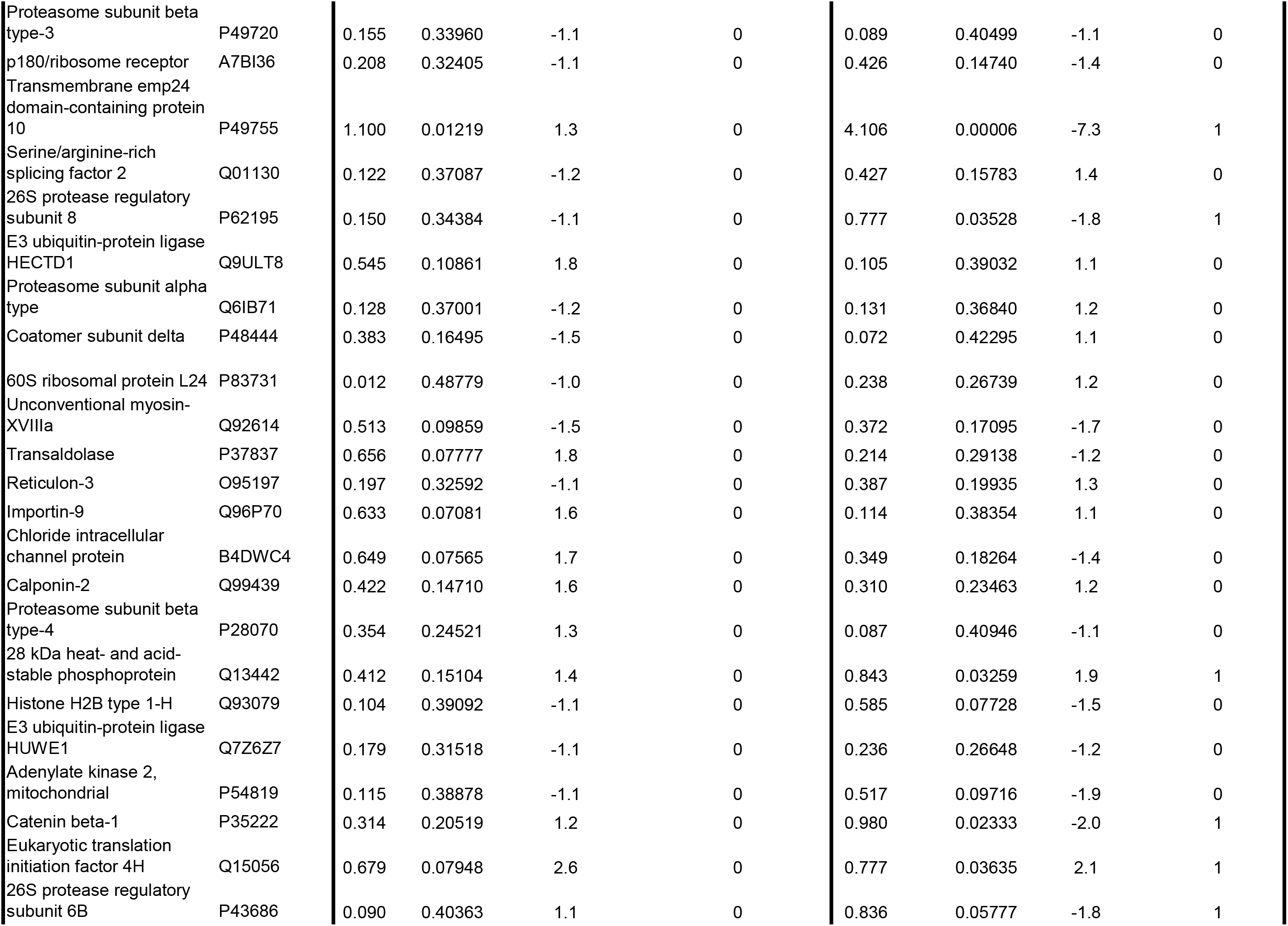
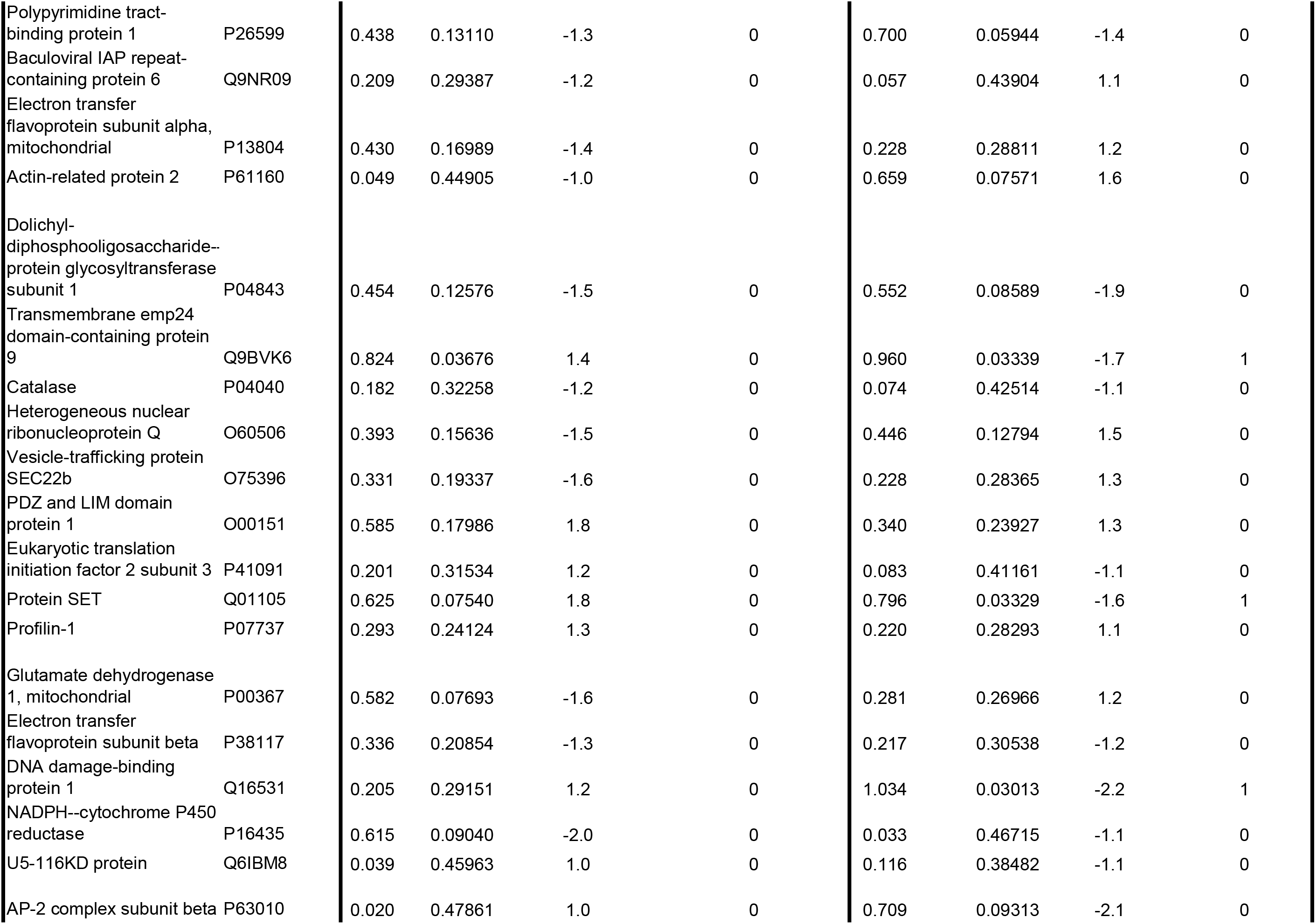
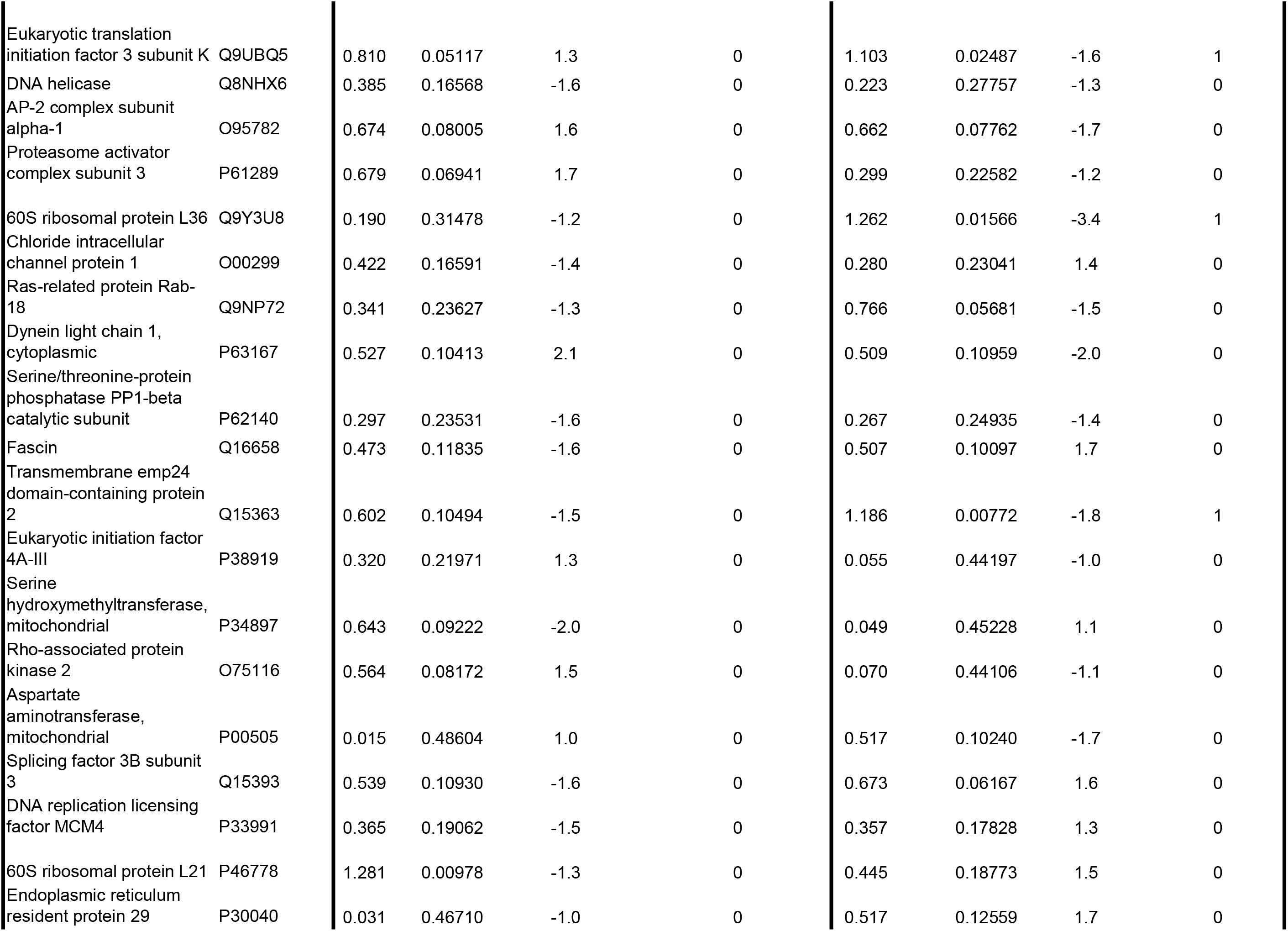
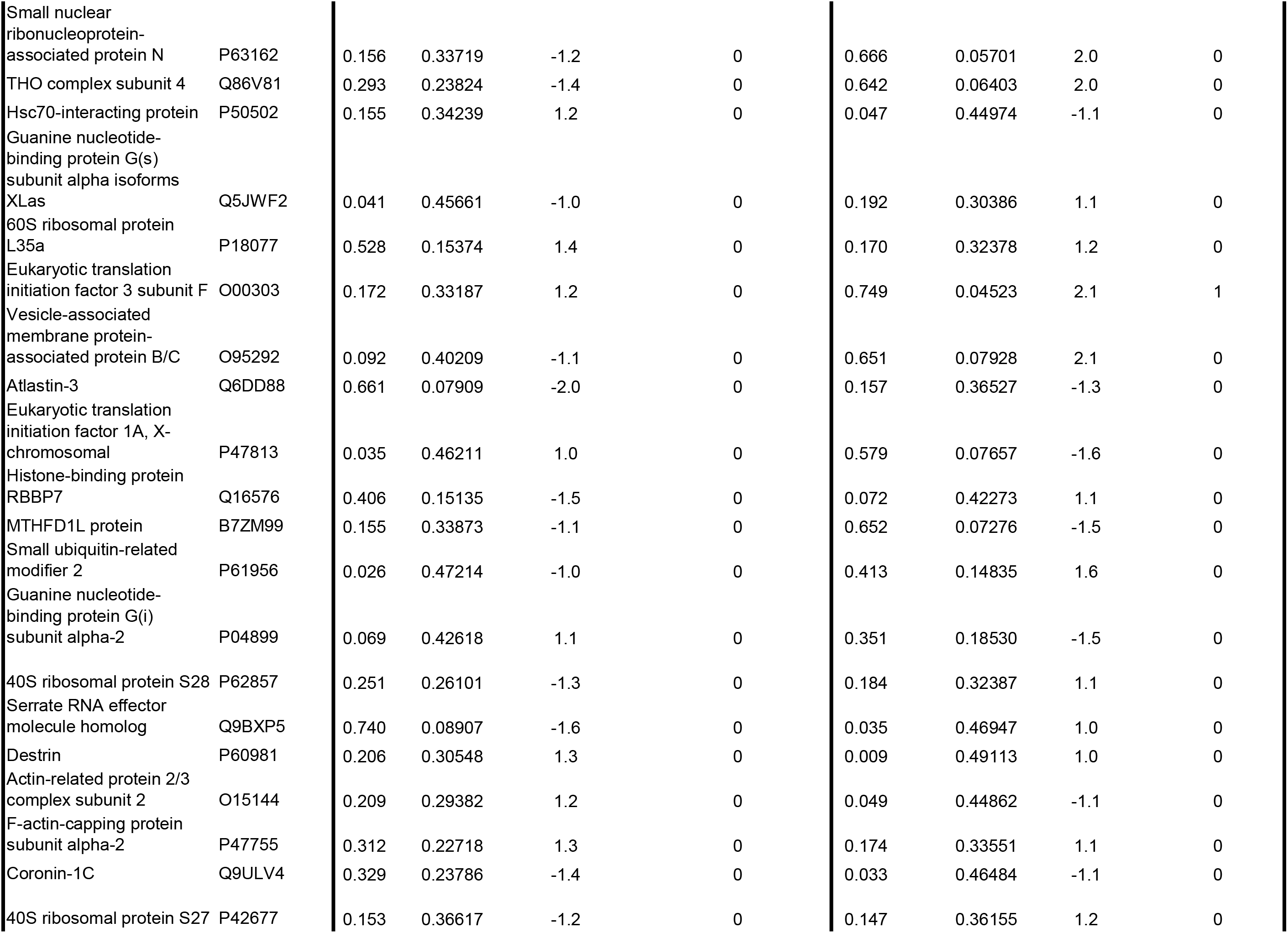
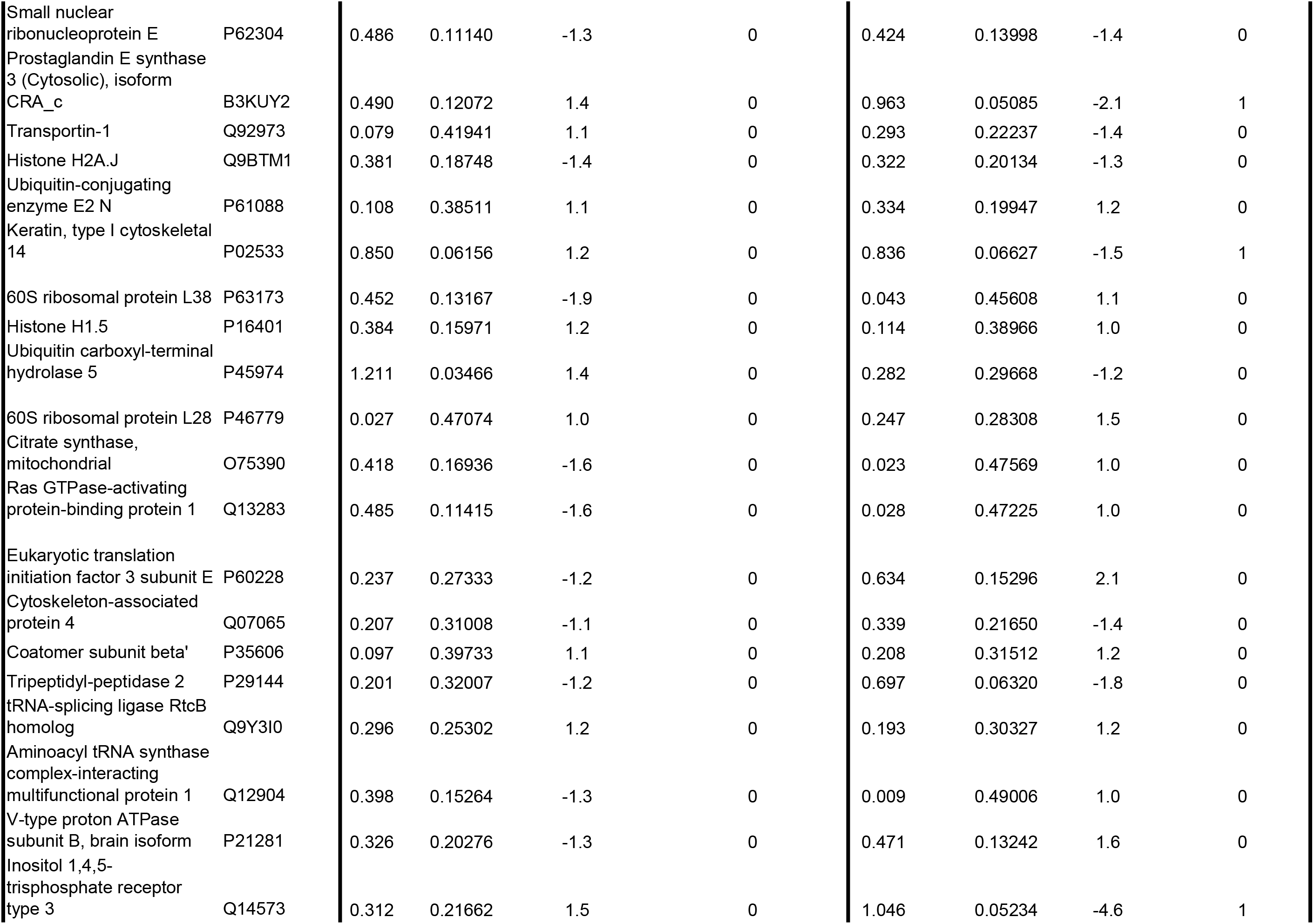
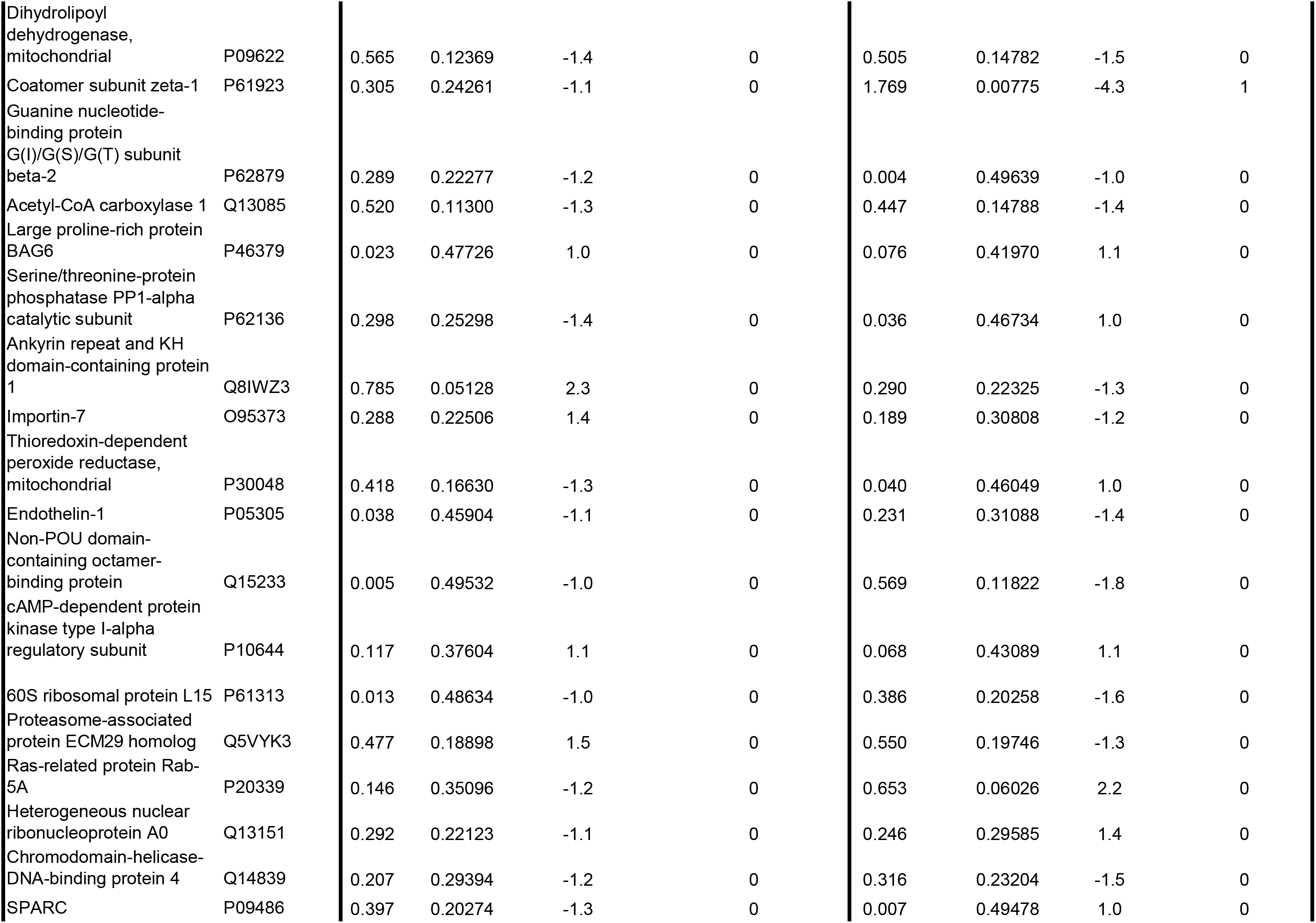
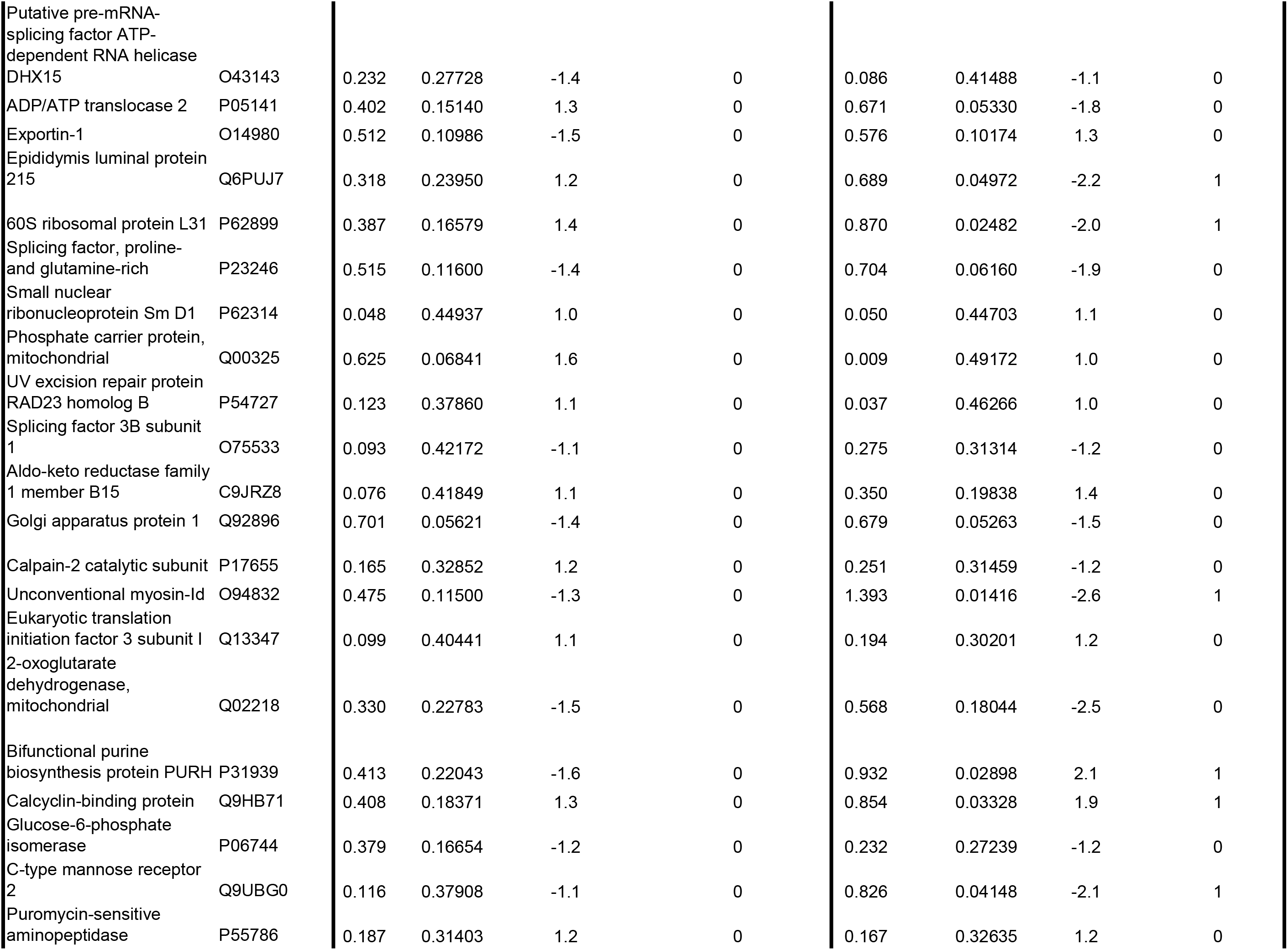
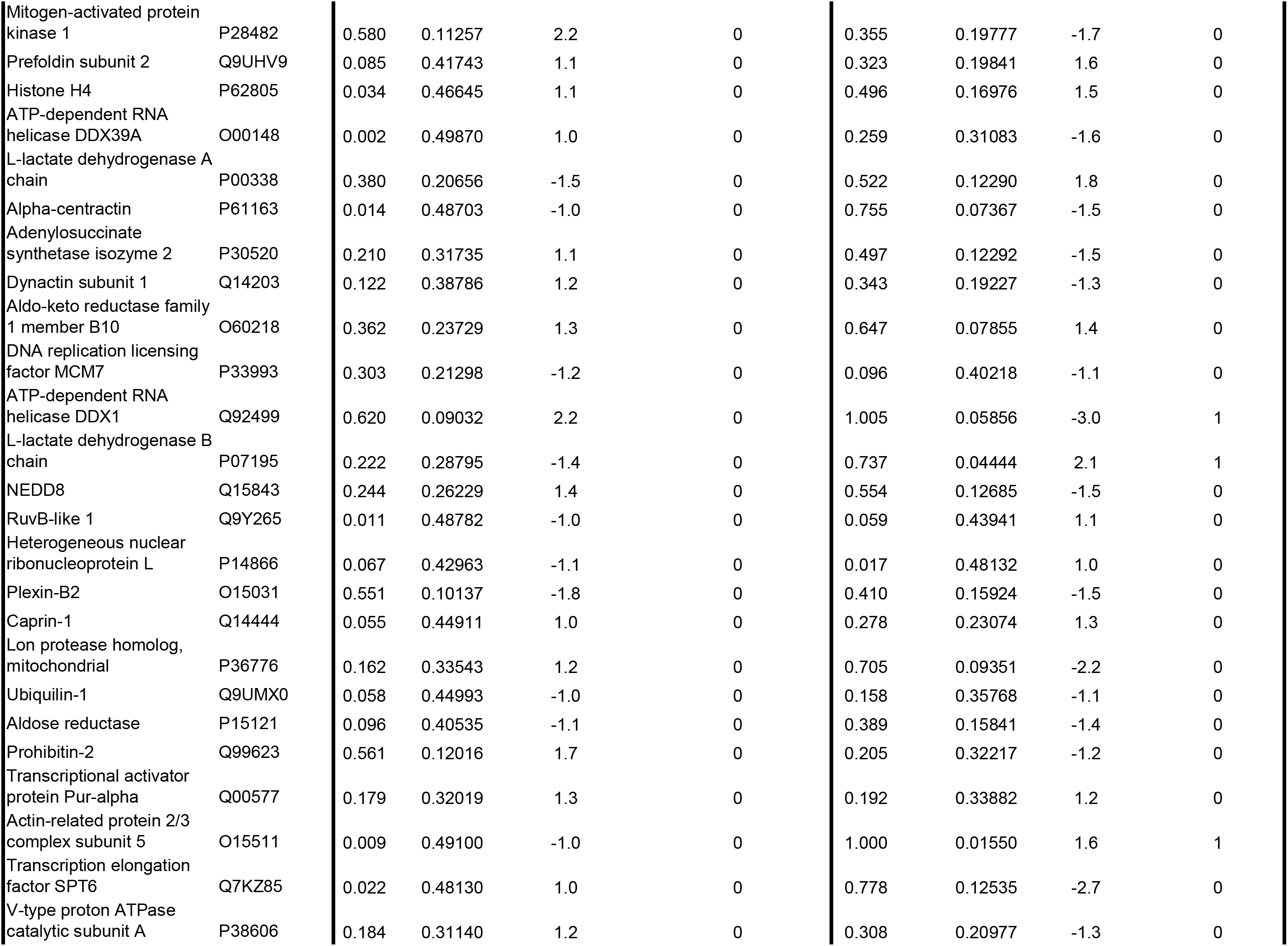
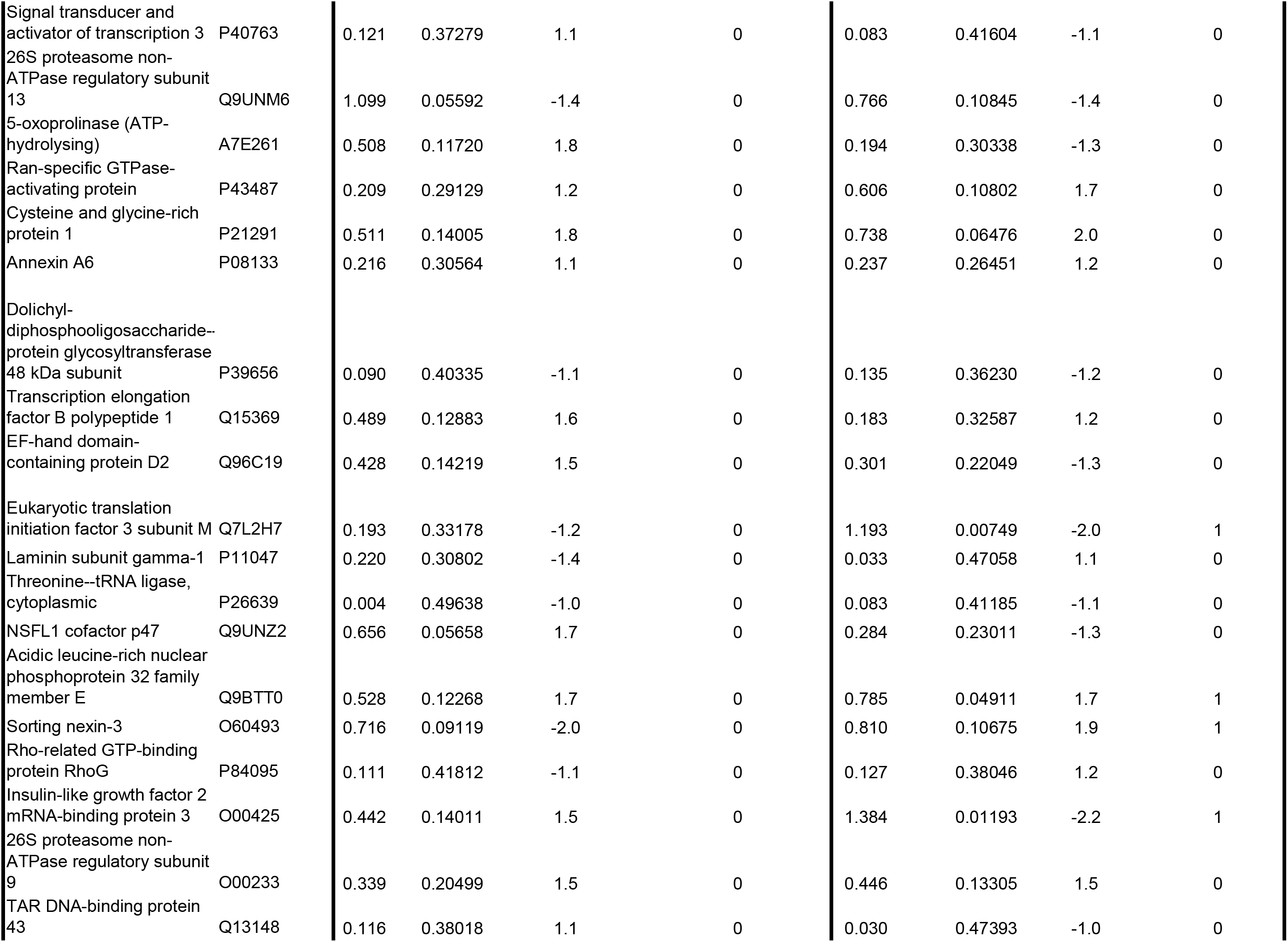
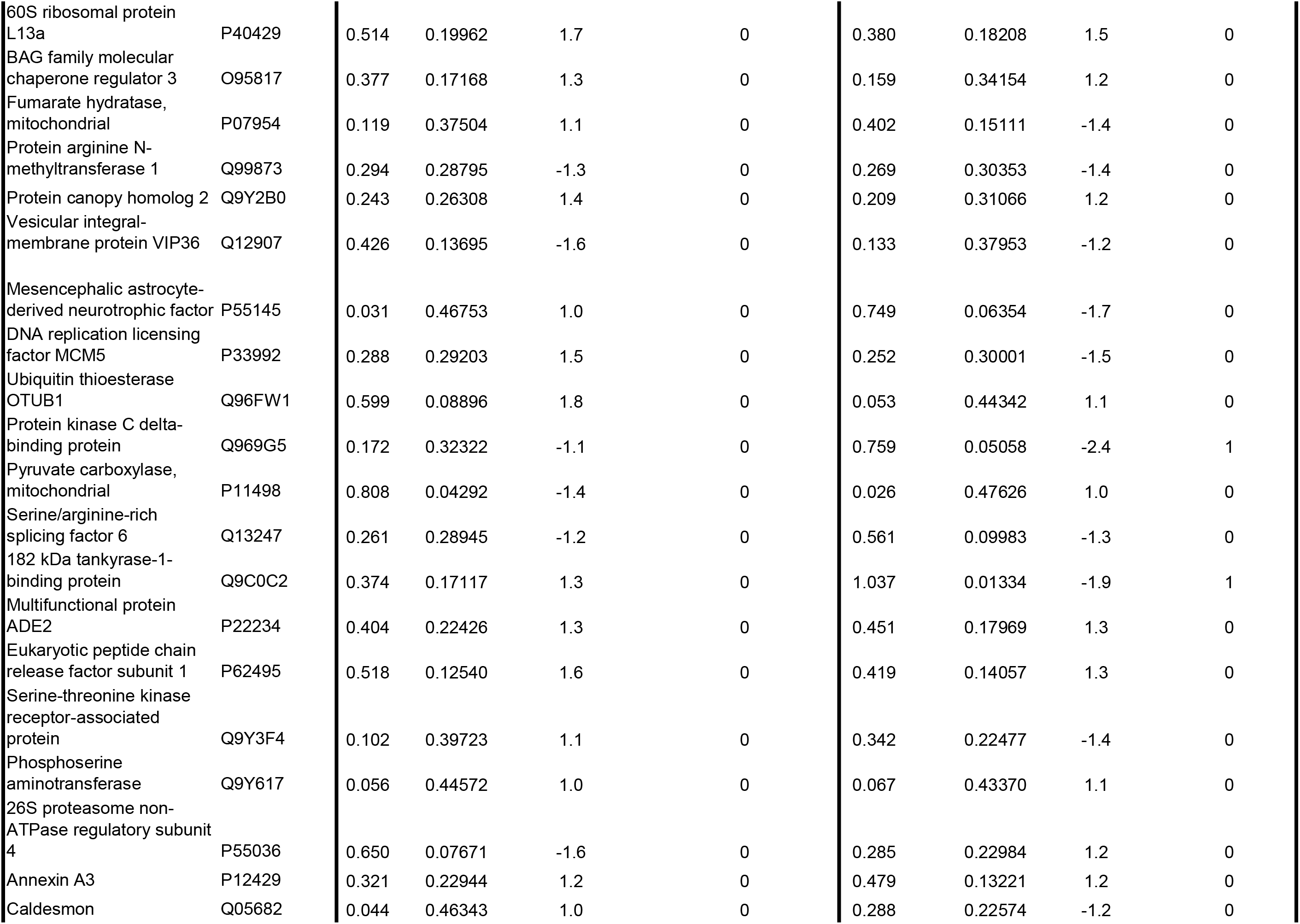
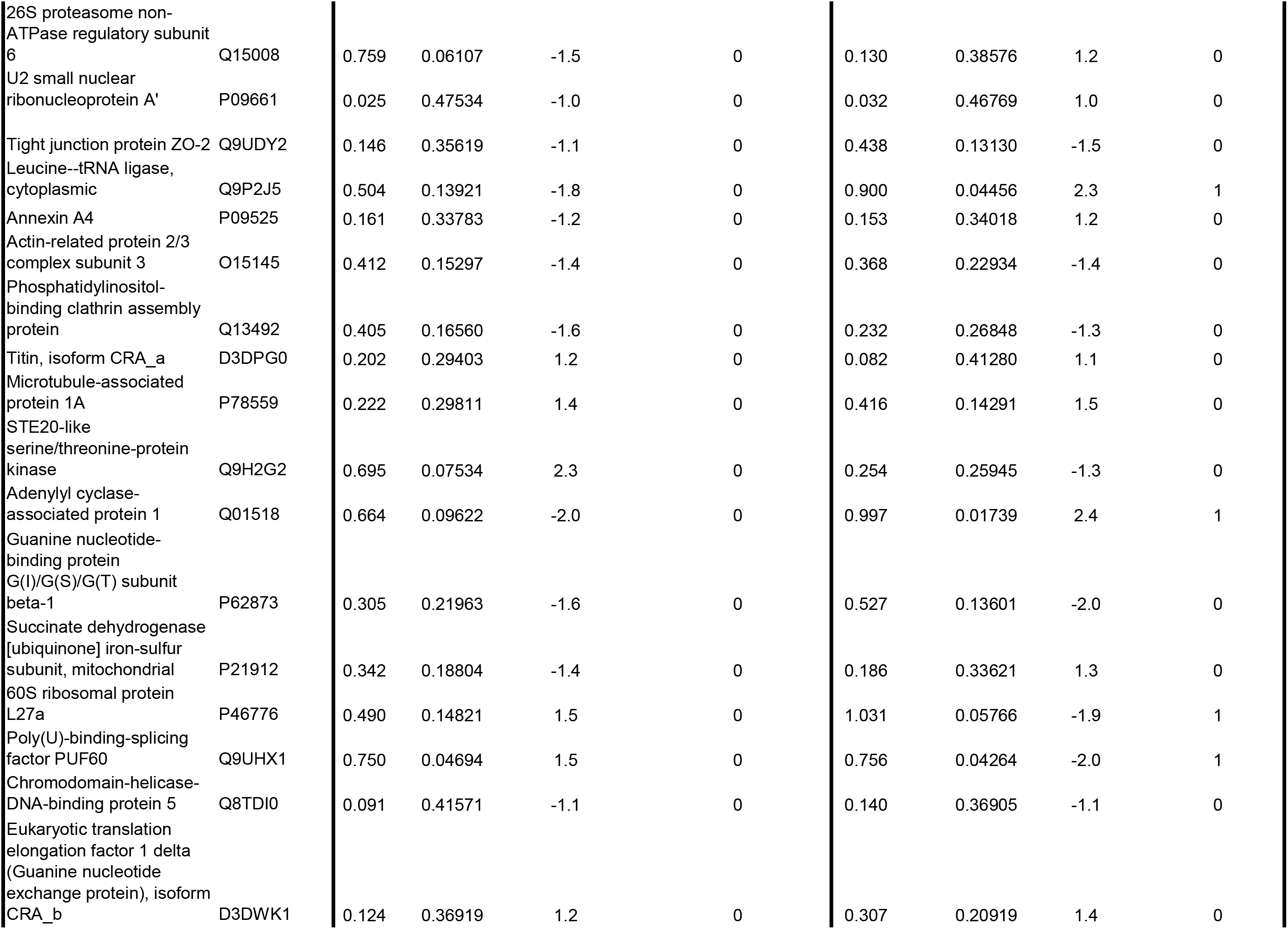
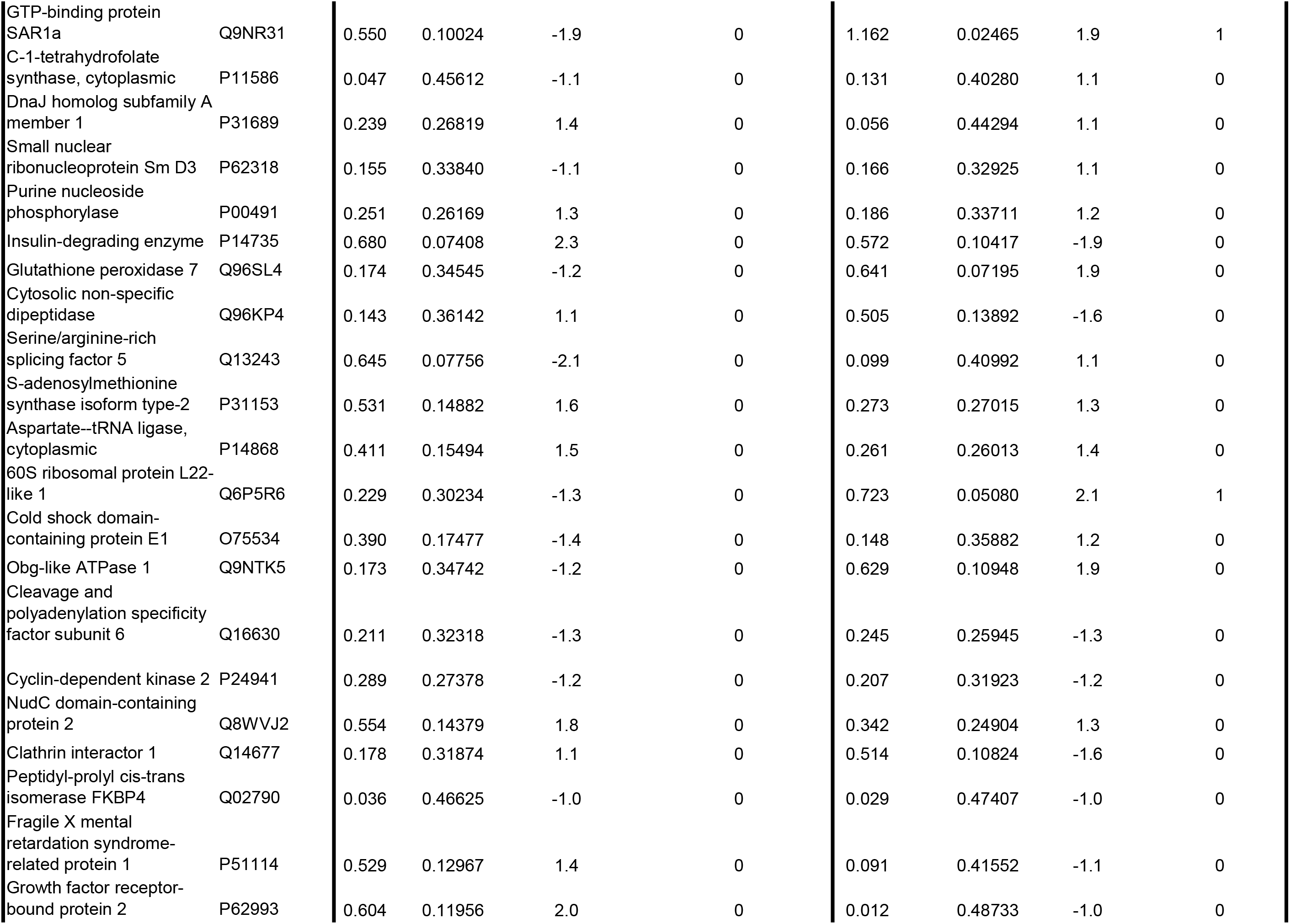
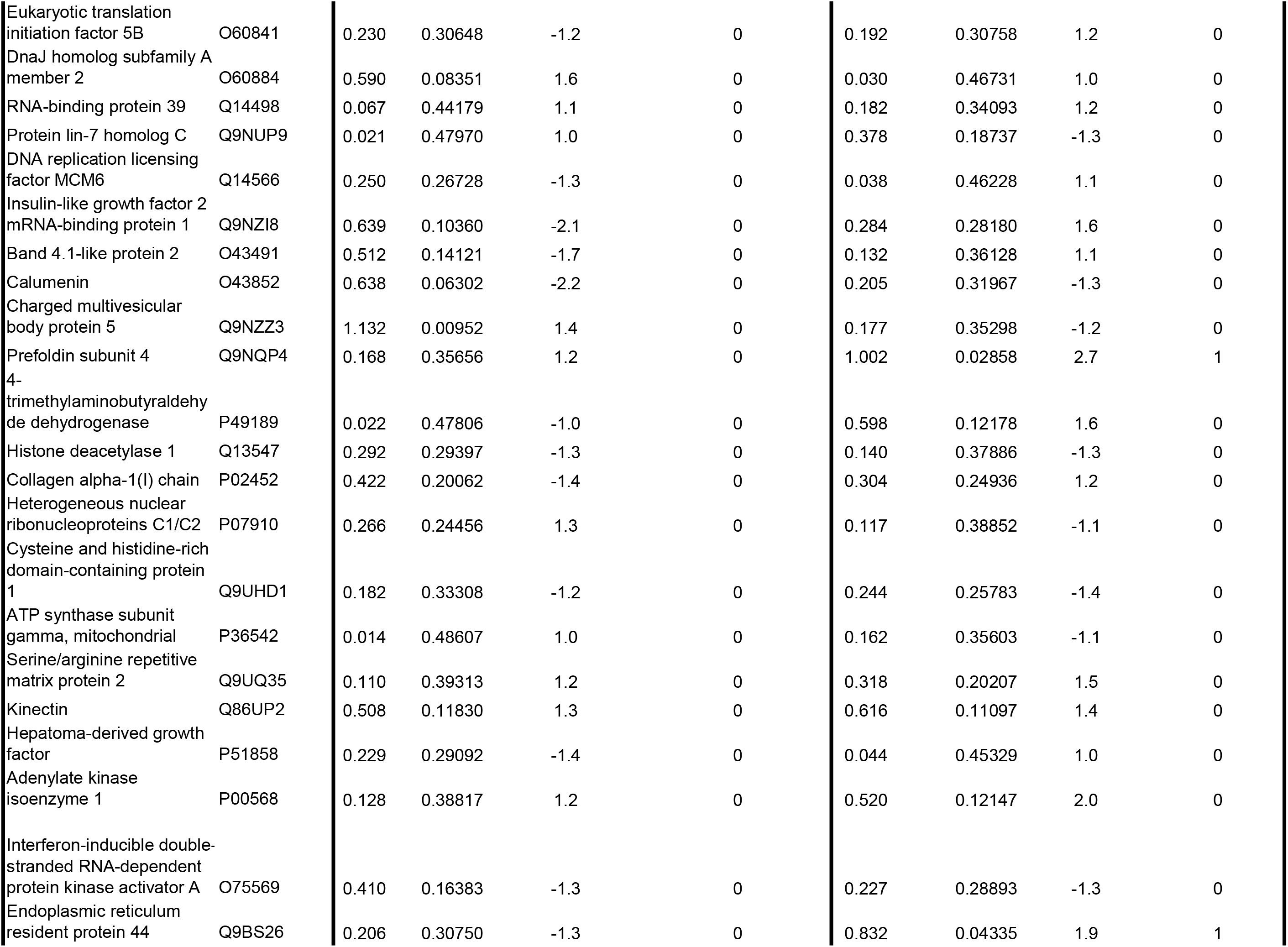
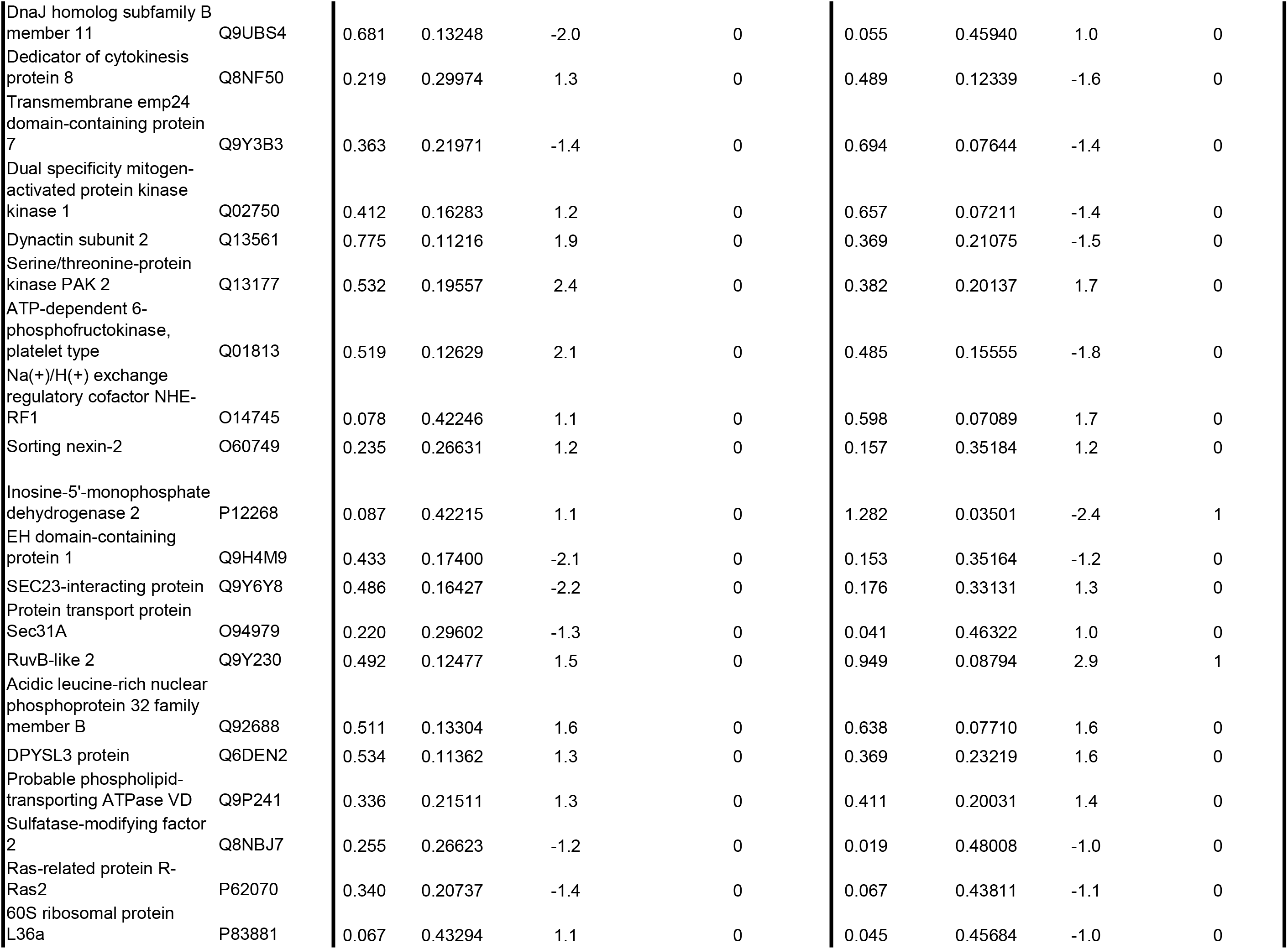
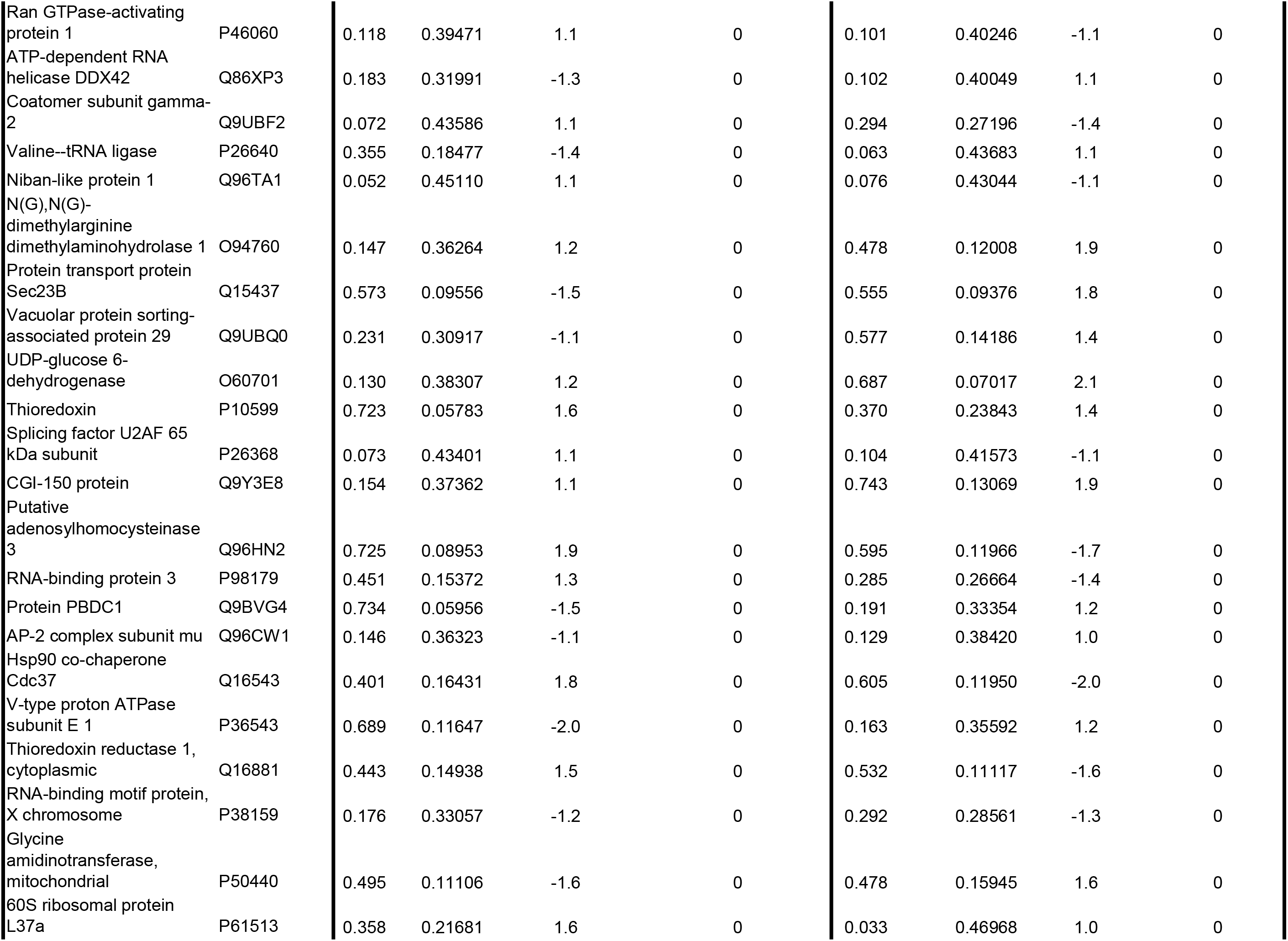
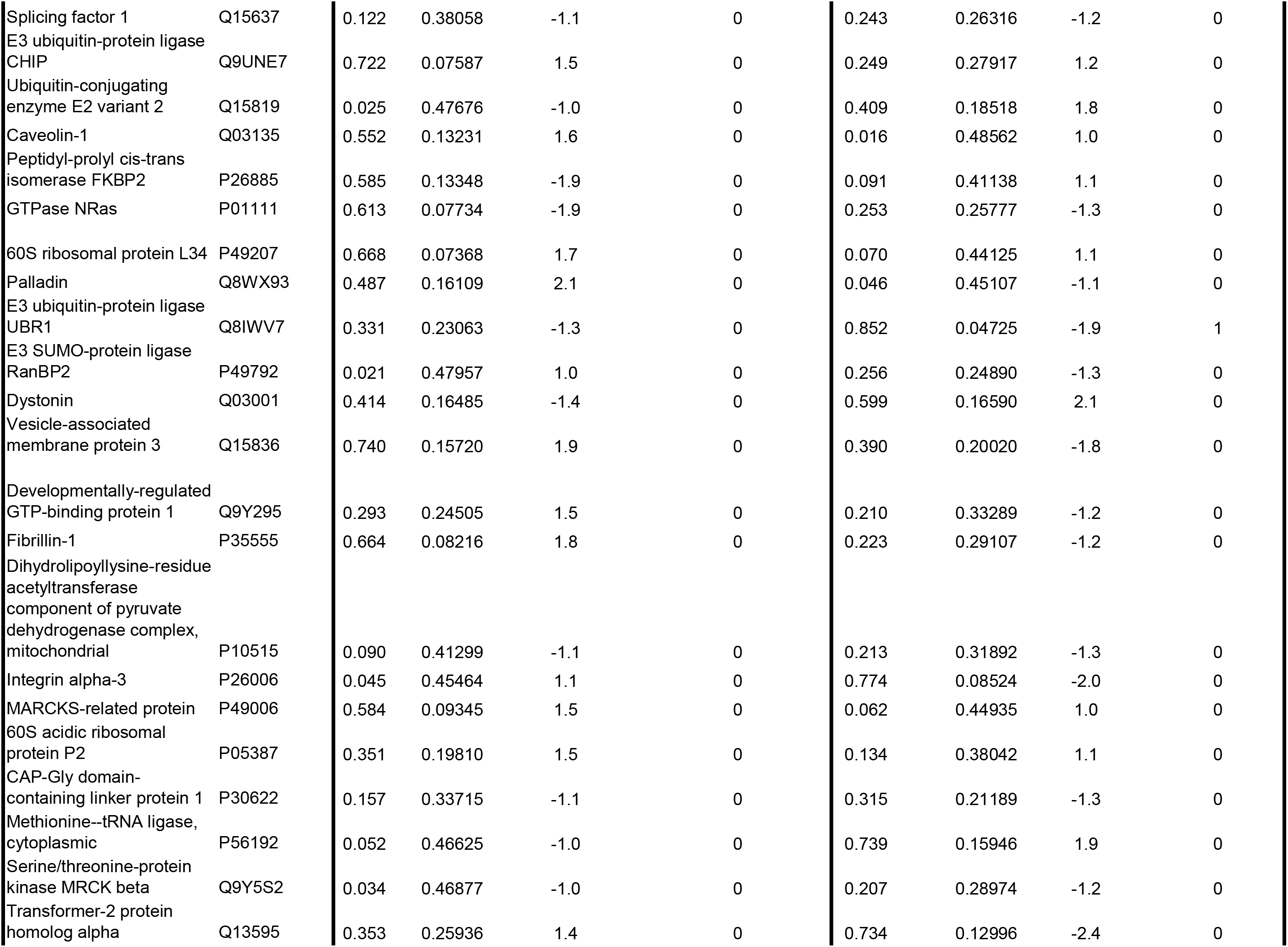
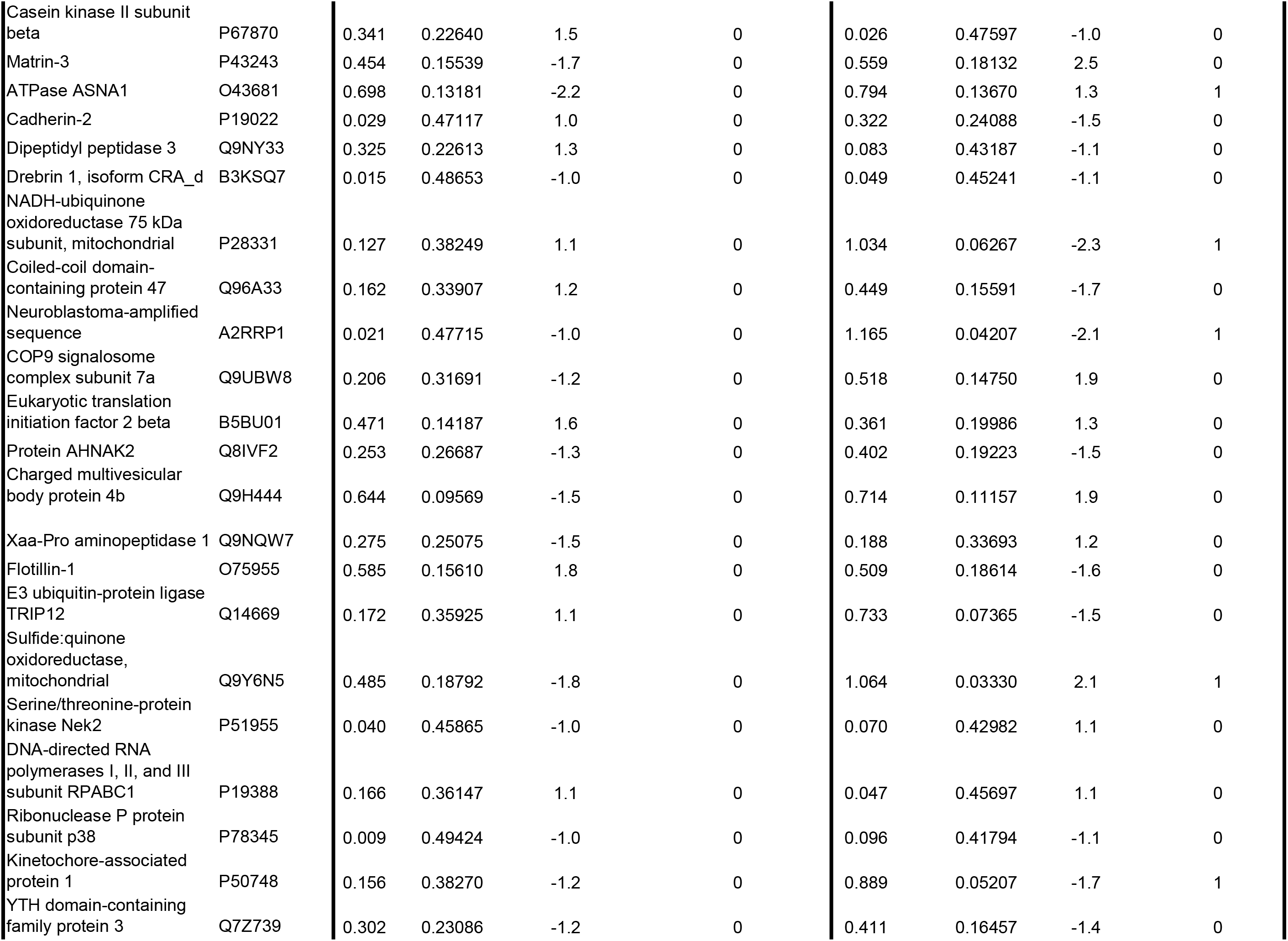
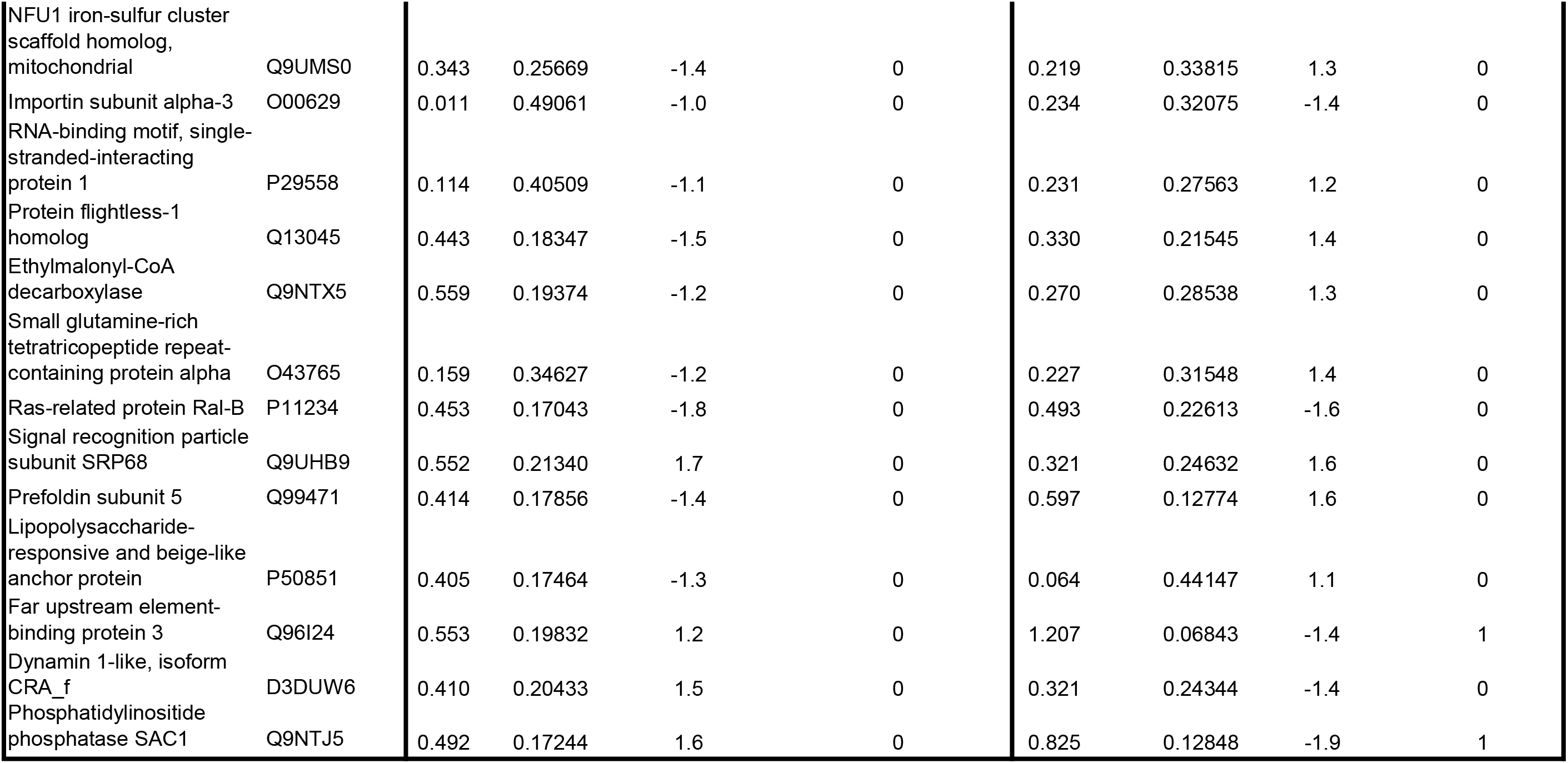
Full List; Seven hundred and four (734) proteins with at least three of four values >0)

**Supplemental Table 2.**
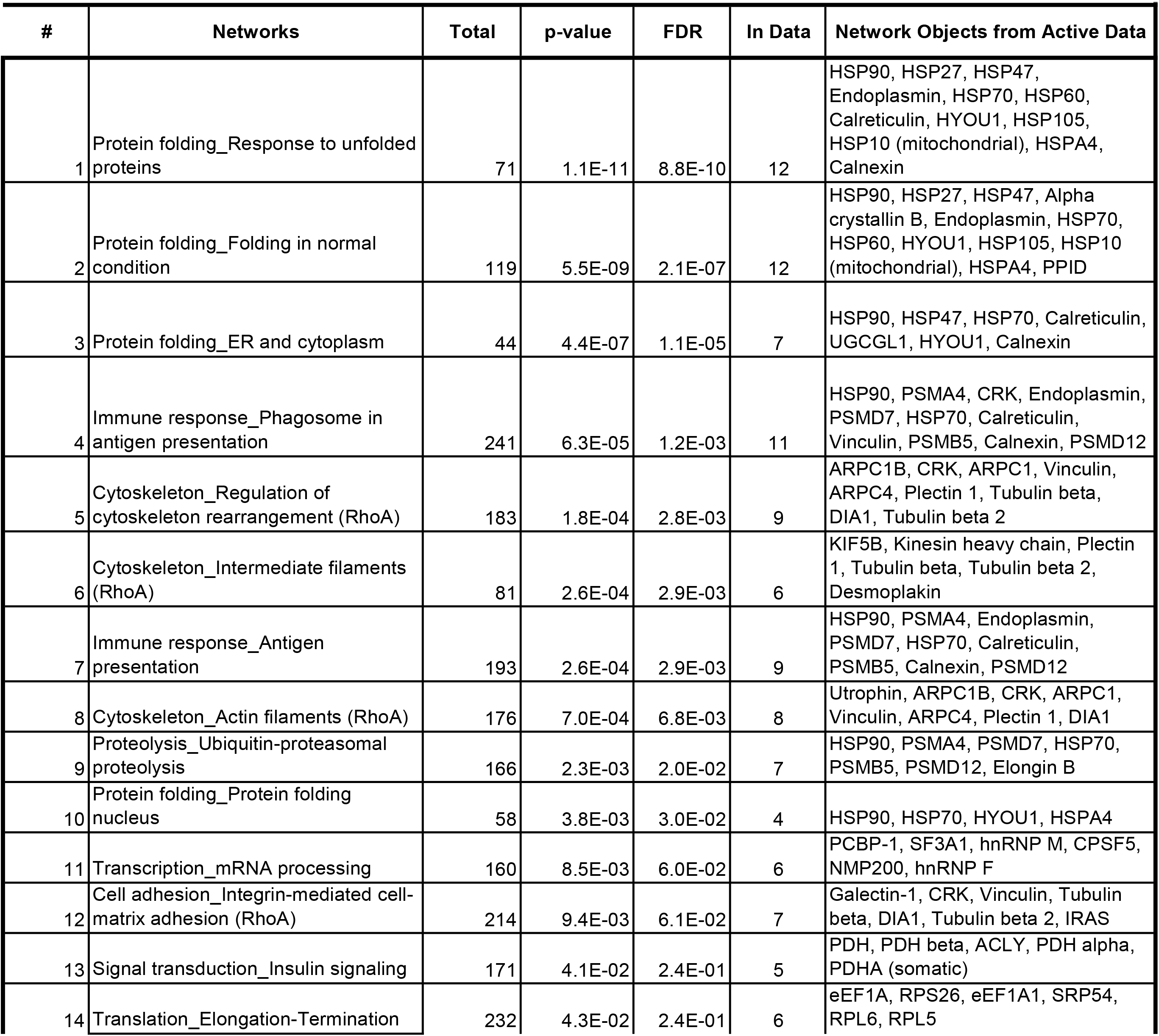
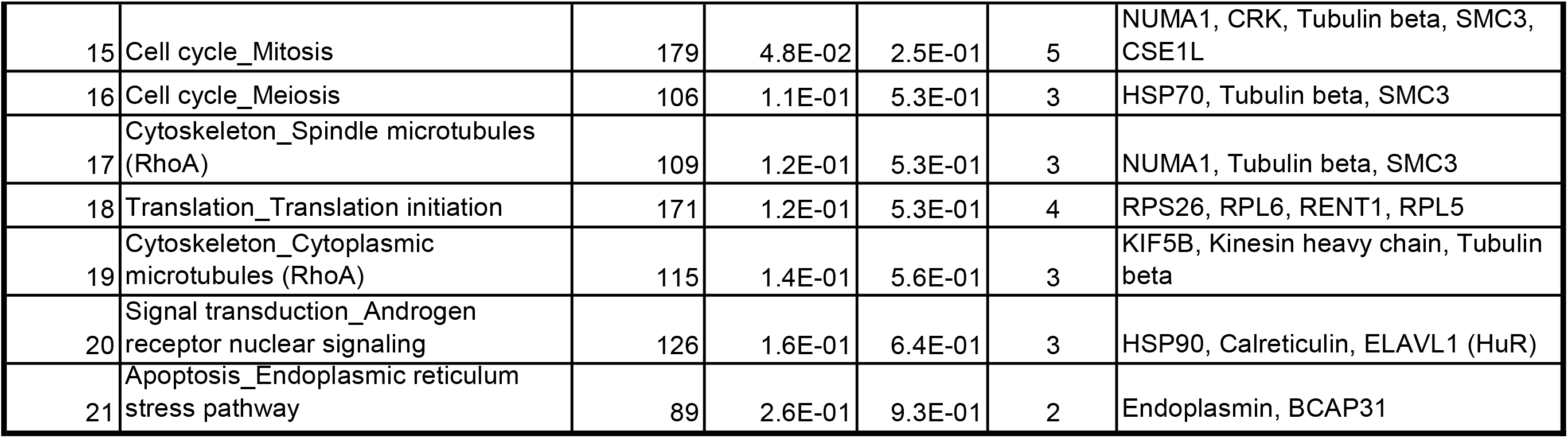
Enrichment by Process Networks (S1 treated vs. Vehicle)

**Supplemental Table 3.** Sixty five (65) Most Significantly Reversed Proteins by pre HMW-HA -Tx in S1 vs S1and C groups.

| UniProtKB | Acc# | Statistics (C vs S1) |  |  | Fold (HMW-HA,S1/S1) | HA Significantly Reversed (yes/ no) |
| --- | --- | --- | --- | --- | --- | --- |
|  |  | SAM | Ttest | Fold (S1/C) |  |  |
| Trifunctional enzyme subunit alpha, mitochondrial | P40939 | 1.180 | 0.0694 | -4.6 | 2.5 | N |
| Basement membrane-specific heparan sulfate proteoglycan core protein | P98160 | 1.538 | 0.0077 | -3.7 | 1.9 | N |
| Signal recognition particle 54 kDa protein | P61011 | 1.261 | 0.0414 | -3.0 | 3.1 | Y |
| Ras-related protein Rab-5B | P61020 | 0.974 | 0.0260 | -2.8 | 2.2 | Y |
| Pyridoxal-dependent decarboxylase domain-containing protein 1 | Q6P996 | 1.416 | 0.0305 | -2.3 | 1.4 | N |
| Heterogeneous nuclear ribonucleoprotein M | P52272 | 1.364 | 0.0084 | -2.3 | 1.9 | Y |
| Prostacyclin synthase | Q16647 | 0.878 | 0.0409 | -2.3 | 2.2 | N |
| Tumor protein D54 | O43399 | 1.158 | 0.0143 | -2.3 | 1.8 | Y |
| Vacuolar protein sorting-associated protein 13A | Q96RL7 | 1.283 | 0.0408 | -2.3 | 2.4 | Y |
| Cleavage and polyadenylation specificity factor subunit 5 | O43809 | 1.021 | 0.0399 | -2.2 | 1.3 | N |
| Pre-mRNA-processing factor 19 | Q9UMS4 | 1.235 | 0.0067 | -2.2 | 2.4 | Y |
| Lanosterol 14-alpha demethylase | Q16850 | 0.787 | 0.0500 | -2.1 | 1.9 | N |
| RNA-binding protein FUS | P35637 | 0.913 | 0.0430 | -2.1 | 1.2 | N |
| 10 kDa heat shock protein, mitochondrial | P61604 | 1.266 | 0.0070 | -2.0 | 1.2 | N |
| Hypoxia up-regulated protein 1 | Q9Y4L1 | 0.910 | 0.0283 | -2.0 | 1.4 | N |
| Staphylococcal nuclease domain-containing protein 1 | Q7KZF4 | 1.799 | 0.0122 | -2.0 | 1.2 | N |
| ELAV-like protein 1 | Q15717 | 0.930 | 0.0288 | -2.0 | 1.1 | N |
| 26S proteasome non-ATPase regulatory subunit 7 | P51665 | 0.785 | 0.0462 | -1.9 | 1.2 | N |
| Prolyl 4-hydroxylase subunit alpha-2 | O15460 | 0.825 | 0.0370 | -1.9 | 1.3 | N |
| Actin-related protein 2/3 complex subunit 1B | O15143 | 0.931 | 0.0194 | -1.7 | 1.3 | N |
| Utrophin | P46939 | 1.480 | 0.0046 | -1.7 | 1.1 | N |
| Vesicle-associated membrane protein-associated protein A | Q9P0L0 | 1.110 | 0.0176 | -1.7 | 2.1 | Y |
| Calnexin | P27824 | 0.788 | 0.0453 | -1.7 | 1.0 | N |
| Alpha-aminoadipic semialdehyde dehydrogenase | P49419 | 1.816 | 0.0374 | -1.7 | 1.4 | N |
| Calreticulin | P27797 | 1.545 | 0.0061 | -1.7 | 1.1 | N |
| Polymerase I and transcript release factor | Q6NZI2 | 1.729 | 0.0121 | -1.7 | 1.4 | N |
| Coatmer subunit alpha | P53621 | 1.058 | 0.0130 | -1.6 | 1.4 | N |
| Serpin H1 | P50454 | 1.248 | 0.0230 | -1.6 | 1.1 | N |
| Neuroblast differentiation-associated protein AHNAK | Q09666 | 1.382 | 0.0059 | -1.6 | 1.1 | N |
| Endoplasmin | P14625 | 1.214 | 0.0226 | -1.5 | 1.0 | N |
| Malate dehydrogenase | Q6FHZ0 | 1.154 | 0.0138 | -1.5 | 1.1 | N |
| 60S ribosomal protein L5 | P46777 | 1.069 | 0.0205 | 1.5 | -1.5 | Y |
| Kinesin-1 heavy chain | P33176 | 0.785 | 0.0343 | 1.5 | -1.2 | N |
| Tubulin beta-2A chain | Q13885 | 1.858 | 0.0022 | 1.5 | -1.1 | N |
| Plectin | Q15149 | 0.942 | 0.0188 | 1.5 | -1.3 | Y |
| Elongation factor 1-alpha 1 | P68104 | 0.864 | 0.0251 | 1.6 | -1.1 | N |
| Transcription elongation factor B polypeptide 2 | Q15370 | 0.842 | 0.0866 | 1.6 | -2.3 | Y |
| Desmoplakin | P15924 | 1.333 | 0.0119 | 1.7 | -1.5 | Y |
| Drug-sensitive protein 1 | Q9NZ23 | 1.768 | 0.0056 | 1.7 | -1.6 | N |
| Heat shock protein 105 kDa | Q92598 | 0.898 | 0.0616 | 1.7 | -1.1 | N |
| 40S ribosomal protein S26 | P62854 | 0.951 | 0.0221 | 1.7 | -1.3 | N |
| Heat shock 70 kDa protein 4 | P34932 | 0.950 | 0.0205 | 1.7 | -1.6 | Y |
| COP9 signalosome complex subunit 4 | Q9BT78 | 0.811 | 0.0424 | 1.8 | -1.1 | N |
| 60S ribosomal protein L6 | Q02878 | 0.981 | 0.0182 | 1.8 | -1.7 | N |
| Pyruvate dehydrogenase E1 component subunit beta, mitochondrial | P11177 | 0.817 | 0.0301 | 1.8 | -1.2 | N |
| Fructose-bisphosphate aldolase C | P09972 | 1.504 | 0.0142 | 1.8 | -1.2 | N |
| 26S proteasome non-ATPase regulatory subunit 12 | O00232 | 1.332 | 0.0112 | 1.9 | -1.9 | Y |
| Regulator of nonsense transcripts 1 | Q92900 | 0.794 | 0.0482 | 1.9 | -2.1 | Y |
| Golgin subfamily A member 3 | Q08378 | 1.110 | 0.0507 | 1.9 | -1.0 | N |
| Ephrin type-A receptor 2 | P29317 | 0.838 | 0.0424 | 2.0 | -1.2 | N |
| Rab GDP dissociation inhibitor beta | P50395 | 0.806 | 0.0517 | 2.0 | -2.0 | Y |
| Heat shock protein beta-1 | P04792 | 0.793 | 0.0792 | 2.0 | -3.2 | Y |
| Similar to 60S ribosomal protein L7;<br>similar to P18124 (PID:d133021) | O95036 | 2.107 | 0.0006 | 2.1 | -1.4 | Y |
| Protein FAM49B | Q9NUQ9 | 0.950 | 0.0718 | 2.1 | -1.3 | N |
| Phosphoglycerate kinase 1 | P00558 | 1.929 | 0.0010 | 2.2 | -1.0 | N |
| Exportin-2 | P55060 | 1.185 | 0.0079 | 2.2 | -2.4 | Y |
| ATP-binding cassette sub-family E<br>member 1 | P61221 | 1.348 | 0.0070 | 2.2 | -1.2 | N |
| Rab GDP dissociation inhibitor alpha | P31150 | 1.645 | 0.0158 | 2.3 | -1.9 | Y |
| Alpha-crystallin B chain | P02511 | 1.324 | 0.0056 | 2.4 | -1.9 | Y |
| Vacuolar protein sorting-associated<br>protein 26A | O75436 | 1.164 | 0.0873 | 2.5 | -2.2 | N |
| Poly(rC)-binding protein 1 | Q15365 | 1.550 | 0.0058 | 2.5 | -1.1 | N |
| Splicing factor 3A subunit 1 | Q15459 | 1.005 | 0.0341 | 2.7 | -1.2 | N |
| ATP-citrate synthase | P53396 | 1.567 | 0.0027 | 3.0 | -1.4 | N |
| Structural maintenance of chromosomes<br>protein 3 | Q9UQE7 | 1.183 | 0.0223 | 3.2 | -1.4 | N |
| Protein diaphanous homolog 1 | O60610 | 2.667 | 0.0055 | 3.4 | -3.3 | Y |

**Supplemental Table 4.** Seventy-five (75) Most Significantly Change Proteins in HMW-HA v Vehicle Groups (37 Increased & 38 Decreased)

| UniProtKB Name | Acc# | Vehicle |  |  |  | HMW-HA Treated |  |  |  | Stats (HMW-HA vs Vehicle) |  |  |
| --- | --- | --- | --- | --- | --- | --- | --- | --- | --- | --- | --- | --- |
|  |  | C1 | C2 | C3 | C4 | HA1 | HA2 | HA3 | HA4 | SAM | Ttest | Fold (HMW-HA vs. Vehicle) |
| Epididymis luminal protein 215 | Q6PUJ7 | 2.0 | 1.1 | 1.1 | 3.8 | 5.3 | 10.7 | 5.6 | 8.2 | 1.44 | 0.00763 | 3.7 |
| 116 kDa U5 small nuclear ribonucleoprotein component | Q15029 | 5.0 | 1.1 | 2.2 | 3.8 | 10.6 | 7.1 | 19.6 | 4.1 | 0.87 | 0.05737 | 3.4 |
| Endothelin-1 | P05305 | 4.0 | 4.3 | 3.4 | 2.8 | 6.3 | 19.6 | 4.7 | 9.2 | 0.86 | 0.07754 | 2.7 |
| Eukaryotic translation initiation factor 2 subunit 1 | P05198 | 3.0 | 1.1 | 4.5 | 1.9 | 5.3 | 4.5 | 9.3 | 9.2 | 1.10 | 0.01564 | 2.7 |
| Coiled-coil domain-containing protein 47 | Q96A33 | 2.0 | 1.1 | 1.1 |  | 6.3 | 1.8 |  | 3.1 | 0.81 | 0.11194 | 2.7 |
| RNA-binding protein 3 | P98179 | 1.0 | 2.1 |  | 1.9 |  | 3.6 | 4.7 | 5.1 | 1.99 | 0.00511 | 2.7 |
| Clathrin interactor 1 | Q14677 | 1.0 | 3.2 | 1.1 | 1.9 | 3.2 | 7.1 | 3.7 | 5.1 | 1.07 | 0.01722 | 2.7 |
| Drug-sensitive protein 1 | Q9NZ23 | 4.0 | 12.9 | 3.4 | 3.8 | 17.9 | 14.3 | 18.7 | 9.2 | 1.01 | 0.01442 | 2.5 |
| Prohibitin-2 | Q99623 | 2.0 | 2.1 |  | 1.9 | 7.4 | 4.5 | 4.7 | 3.1 | 1.49 | 0.02457 | 2.4 |
| Lon protease homolog, mitochondrial | P36776 | 3.0 | 2.1 | 3.4 | 0.9 |  | 5.4 | 6.5 | 5.1 | 1.80 | 0.00256 | 2.4 |
| Rho GDP-dissociation inhibitor 1 | P52565 | 5.0 | 3.2 | 8.9 | 3.8 | 10.6 | 9.8 | 16.8 | 12.3 | 1.24 | 0.00679 | 2.4 |
| Heparin-binding protein HBp15 | Q7Z4W8 | 2.0 | 2.1 | 2.2 | 0.9 | 1.1 | 6.2 | 4.7 | 5.1 | 0.86 | 0.05762 | 2.3 |
| LDLR chaperone MESD | Q14696 |  | 3.2 | 2.2 | 1.9 | 6.3 | 5.4 | 7.5 | 3.1 | 1.21 | 0.01897 | 2.3 |
| Proliferation-associated protein 2G4 | Q9UQ80 | 4.0 | 2.1 | 4.5 | 1.9 | 6.3 | 8.0 | 5.6 | 8.2 | 1.52 | 0.00257 | 2.2 |
| Cysteine and glycine-rich protein 1 | P21291 | 3.0 | 2.1 | 2.2 | 0.9 | 3.2 | 7.1 | 3.7 |  | 0.86 | 0.08043 | 2.2 |
| Glycylpeptide N-tetradecanoyltransferase 1 | P30419 | 4.0 | 1.1 | 2.2 |  | 7.4 |  | 3.7 | 5.1 | 0.89 | 0.04926 | 2.2 |
| 60S ribosomal protein L7 | P18124 | 4.0 | 8.6 | 4.5 | 6.6 | 16.9 | 7.1 | 8.4 | 19.4 | 0.86 | 0.04996 | 2.2 |
| RNA-binding motif protein, X chromosome | P38159 | 1.0 | 1.1 | 3.4 | 4.7 | 5.3 | 6.2 | 6.5 | 4.1 | 1.03 | 0.01872 | 2.2 |
| Proteasome subunit beta type-7 | Q99436 | 2.0 |  | 1.1 | 0.9 |  | 2.7 | 3.7 | 2.0 | 1.02 | 0.03944 | 2.1 |
| Histone H2B type 1-M | Q99879 | 3.0 | 2.1 | 4.5 | 1.9 | 4.2 | 5.4 | 7.5 |  | 1.00 | 0.03805 | 2.0 |
| Calpain small subunit 1 | P04632 | 2.0 | 1.1 |  | 0.9 | 2.1 | 2.7 | 3.7 | 2.0 | 0.95 | 0.02667 | 2.0 |
| Histone H4 | P62805 | 5.0 | 3.2 | 2.2 | 3.8 | 7.4 | 8.9 | 4.7 |  | 1.03 | 0.04549 | 2.0 |
| B-cell receptor-associated protein 31 | P51572 |  | 3.2 | 2.2 | 1.9 | 2.1 | 5.4 | 6.5 | 5.1 | 0.91 | 0.04258 | 2.0 |
| Ubiquitin-conjugating enzyme E2 N | P61088 | 2.0 |  | 1.1 | 1.9 | 3.2 | 2.7 | 4.7 | 2.0 | 0.92 | 0.03730 | 1.9 |
| L-lactate dehydrogenase A chain | P00338 | 6.0 | 5.4 | 4.5 | 2.8 | 8.4 | 6.2 | 12.1 | 7.2 | 0.96 | 0.02624 | 1.8 |
| C-Jun-amino-terminal kinase-interacting protein 4 | O60271 | 1.0 | 1.1 | 1.1 |  | 2.1 | 0.9 | 2.8 |  | 0.85 | 0.12960 | 1.8 |
| Phospholipase D3 | Q8IV08 | 2.0 | 1.1 | 2.2 | 3.8 | 4.2 | 3.6 | 5.6 | 3.1 | 0.83 | 0.02865 | 1.8 |
| Similar to 60S ribosomal protein L7 | O95036 | 6.0 | 13.9 | 7.8 | 5.7 | 16.9 | 11.6 | 11.2 | 20.4 | 0.81 | 0.03202 | 1.8 |
| Adenylyl cyclase-associated protein 1 | Q01518 | 2.0 | 3.2 | 2.2 | 3.8 |  | 5.4 | 4.7 | 5.1 | 1.91 | 0.00363 | 1.8 |
| Src substrate cortactin | Q14247 | 4.0 | 4.3 |  | 2.8 |  | 7.1 | 4.7 | 8.2 | 1.15 | 0.04423 | 1.8 |
| Dihydropyrimidinase-related protein 2 | Q16555 | 9.0 | 5.4 | 3.4 | 4.7 | 10.6 | 8.0 | 11.2 | 9.2 | 1.07 | 0.01695 | 1.7 |
| 40S ribosomal protein S6 | P62753 | 8.0 | 8.6 | 7.8 | 9.4 | 7.4 | 13.4 | 15.0 | 20.4 | 0.92 | 0.06435 | 1.7 |
| Destrin | P60981 | 2.0 | 2.1 | 3.4 | 3.8 | 6.3 | 4.5 | 4.7 | 3.1 | 0.82 | 0.03546 | 1.6 |
| Prolyl 4-hydroxylase subunit alpha-1 | P13674 | 9.0 | 8.6 | 4.5 | 10.4 | 15.8 | 15.2 | 13.1 | 9.2 | 0.94 | 0.01942 | 1.6 |
| Guanine nucleotide-binding protein G(I)/G(S)/G(T) subunit beta-2 | P62879 | 5.0 | 5.4 | 2.2 | 4.7 | 6.3 | 6.2 | 9.3 | 6.1 | 0.90 | 0.02216 | 1.6 |
| Small nuclear ribonucleoprotein Sm D2 | P62316 | 2.0 |  | 1.1 | 1.9 | 3.2 | 2.7 |  | 2.0 | 0.92 | 0.04526 | 1.6 |
| Ribosomal protein L19 | J3QR09 | 9.0 | 11.8 | 6.7 | 5.7 | 12.7 | 12.5 | 10.3 | 15.3 | 0.92 | 0.02256 | 1.5 |
| Poly(rC)-binding protein 1 | Q15365 | 23.1 | 16.1 | 16.8 | 19.8 | 12.7 | 8.9 | 10.3 | 16.4 | 1.06 | 0.01194 | -1.6 |
| Coatomer subunit alpha | P53621 | 23.1 | 21.4 | 20.1 | 25.5 | 20.1 | 14.3 | 9.3 | 13.3 | 1.23 | 0.01213 | -1.6 |
| Insulin-like growth factor 2 mRNA-binding protein 3 | O00425 | 3.0 | 4.3 |  | 3.8 | 2.1 |  | 2.8 | 2.0 | 1.29 | 0.02196 | -1.6 |
| 26S proteasome non-ATPase regulatory subunit 14 | O00487 | 14.1 | 9.6 | 8.9 | 8.5 | 8.4 | 7.1 | 6.5 | 3.1 | 0.82 | 0.03011 | -1.6 |
| 60S ribosomal protein L23a | P62750 | 18.1 | 16.1 | 22.3 | 9.4 | 6.3 | 12.5 | 10.3 | 11.2 | 0.80 | 0.04692 | -1.6 |
| SAP domain-containing ribonucleoprotein | P82979 | 2.0 | 3.2 |  | 2.8 |  | 1.8 | 0.9 | 2.0 | 0.92 | 0.04410 | -1.7 |
| Ran-specific GTPase-activating protein | P43487 | 4.0 | 4.3 |  | 5.7 | 4.2 | 1.8 | 2.8 | 2.0 | 0.98 | 0.02449 | -1.7 |
| DNA replication licensing factor MCM4 | P33991 | 7.0 | 7.5 | 7.8 | 4.7 | 3.2 | 5.4 | 3.7 | 3.1 | 1.19 | 0.00870 | -1.8 |
| Endoplasmic reticulum resident protein 29 | P30040 | 7.0 | 7.5 | 6.7 | 5.7 | 4.2 | 1.8 | 3.7 | 5.1 | 1.38 | 0.00752 | -1.8 |
| Ras-related protein Rab-5C | P51148 | 13.1 | 9.6 | 6.7 | 11.3 | 8.4 | 7.1 | 4.7 | 2.0 | 0.83 | 0.02833 | -1.8 |
| Small ubiquitin-related modifier 1 | P63165 | 2.0 | 3.2 | 2.2 | 1.9 | 1.1 | 1.8 | 0.9 |  | 1.02 | 0.02188 | -1.9 |
| Heme oxygenase 2 | P30519 | 1.0 | 1.1 | 2.2 | 2.8 | 1.1 | 0.9 | 0.9 |  | 0.84 | 0.08138 | -1.9 |
| E3 ubiquitin-protein ligase HUWE1 | Q7Z6Z7 | 14.1 | 7.5 | 6.7 | 14.2 | 6.3 | 5.4 | 3.7 | 7.2 | 0.90 | 0.04352 | -1.9 |
| Synaptic vesicle membrane protein VAT-1 homolog | Q99536 | 12.1 | 10.7 | 8.9 | 16.0 | 2.1 | 5.4 | 7.5 | 10.2 | 0.88 | 0.02442 | -1.9 |
| Serine/arginine-rich splicing factor 5 | Q13243 | 5.0 | 3.2 |  | 5.7 | 3.2 | 2.7 | 1.9 | 2.0 | 1.18 | 0.03957 | -1.9 |
| 40S ribosomal protein S23 | P62266 | 7.0 | 5.4 | 8.9 | 6.6 | 6.3 | 2.7 | 4.7 | 1.0 | 0.87 | 0.03007 | -1.9 |
| U2 small nuclear RNA auxiliary factor 2 isoform b | B5BU25 | 2.0 | 3.2 | 5.6 | 3.8 | 1.1 | 2.7 | 1.9 | 2.0 | 0.80 | 0.04891 | -1.9 |
| Vigilin | Q00341 | 19.1 | 9.6 | 14.5 | 11.3 | 4.2 | 9.8 | 8.4 | 6.1 | 0.98 | 0.02222 | -1.9 |
| Protein disulfide-isomerase A6 | Q15084 | 33.2 | 19.3 | 26.8 | 25.5 | 17.9 | 12.5 | 15.0 | 9.2 | 1.33 | 0.00668 | -1.9 |
| 40S ribosomal protein S8 | P62241 | 24.1 | 22.5 | 32.4 | 31.1 | 19.0 | 12.5 | 11.2 | 13.3 | 1.61 | 0.00274 | -2.0 |
| Small glutamine-rich tetratricopeptide repeat-containing protein alpha | O43765 | 4.0 |  | 3.4 | 2.8 | 1.1 | 1.8 | 0.9 | 3.1 | 1.07 | 0.01887 | -2.0 |
| NADH-cytochrome b5 reductase | Q6ZVI6 | 8.0 | 13.9 | 8.9 | 9.4 | 3.2 | 1.8 | 8.4 | 6.1 | 0.93 | 0.01992 | -2.1 |
| Proteasome subunit beta type-3 | P49720 | 6.0 | 3.2 | 4.5 | 7.5 | 3.2 | 3.6 | 1.9 | 1.0 | 0.95 | 0.02332 | -2.2 |
| Serine-protein kinase ATM | Q13315 | 2.0 | 1.1 | 3.4 |  |  | 0.9 | 0.9 | 1.0 | 0.99 | 0.10617 | -2.3 |
| Endoplasmic reticulum resident protein 44 | Q9BS26 | 3.0 | 5.4 | 2.2 |  |  | 1.8 | 1.9 | 1.0 | 0.95 | 0.08095 | -2.3 |
| 10 kDa heat shock protein, mitochondrial | P61604 | 20.1 | 22.5 | 7.8 | 11.3 | 6.3 | 8.9 | 4.7 | 7.2 | 0.99 | 0.04292 | -2.3 |
| 60S ribosomal protein L38 | P63173 | 3.0 | 5.4 | 6.7 | 7.5 | 3.2 | 1.8 | 2.8 | 2.0 | 1.22 | 0.02097 | -2.3 |
| Atlastin-3 | Q6DD88 | 8.0 | 10.7 | 7.8 | 8.5 | 3.2 | 6.2 | 1.9 | 3.1 | 1.62 | 0.00258 | -2.4 |
| DNA replication licensing factor MCM2 | P49736 | 5.0 | 8.6 | 10.1 | 15.1 | 1.1 | 3.6 | 4.7 | 6.1 | 0.92 | 0.03068 | -2.5 |
| ER membrane protein complex subunit 8 | O43402 |  | 3.2 | 1.1 | 2.8 |  | 0.9 | 0.9 | 1.0 | 1.22 | 0.07751 | -2.5 |
| Alpha-aminoadipic semialdehyde dehydrogenase | P49419 | 2.0 | 4.3 | 2.2 | 3.8 | 1.1 | 1.8 | 0.9 | 1.0 | 1.24 | 0.01905 | -2.6 |
| Far upstream element-binding protein 2 | Q92945 | 15.1 | 8.6 | 11.2 | 11.3 | 1.1 | 8.0 | 3.7 | 5.1 | 1.26 | 0.00594 | -2.6 |
| Proteasome (Prosome, macropain) 26S subunit, non-ATPase, 13, isoform CRA_d | B4DJ66 | 3.0 | 4.3 | 2.2 | 4.7 | 2.1 | 0.9 | 0.9 |  | 1.23 | 0.01199 | -2.7 |
| Dehydrogenase/reductase SDR family member 9 | Q9BPW9 | 4.0 | 2.1 | 2.2 | 2.8 | 1.1 |  | 0.9 | 1.0 | 1.95 | 0.01231 | -2.8 |
| Talin-2 | Q9Y4G6 |  | 27.9 | 13.4 | 18.9 | 10.6 | 3.6 |  | 7.2 | 1.20 | 0.03648 | -2.8 |
| Coatomer subunit gamma-1 | Q9Y678 | 5.0 | 4.3 | 12.3 | 9.4 | 2.1 | 1.8 | 0.9 | 6.1 | 0.82 | 0.03663 | -2.8 |
| Spectrin-like protein of the nuclear envelope and Golgi | Q7RTM4 | 2.0 | 1.1 | 6.7 | 2.8 | 1.1 | 0.9 |  | 1.0 | 0.85 | 0.08919 | -3.2 |
| Serine/arginine repetitive matrix protein 2 | Q9UQ35 | 3.0 | 2.1 | 6.7 | 2.8 | 1.1 | 0.9 |  | 1.0 | 1.25 | 0.03970 | -3.7 |

**Supplemental Table 5.**
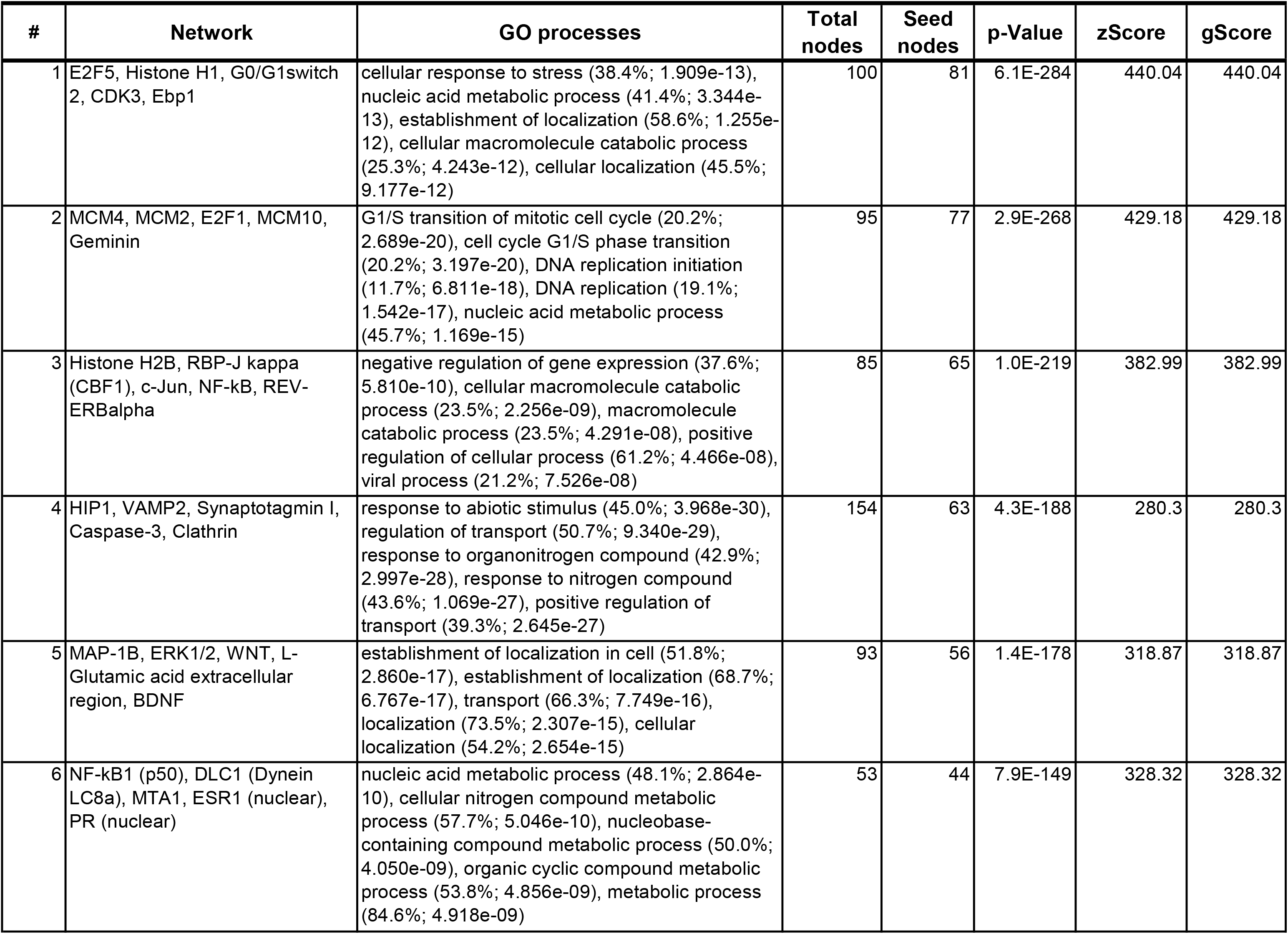
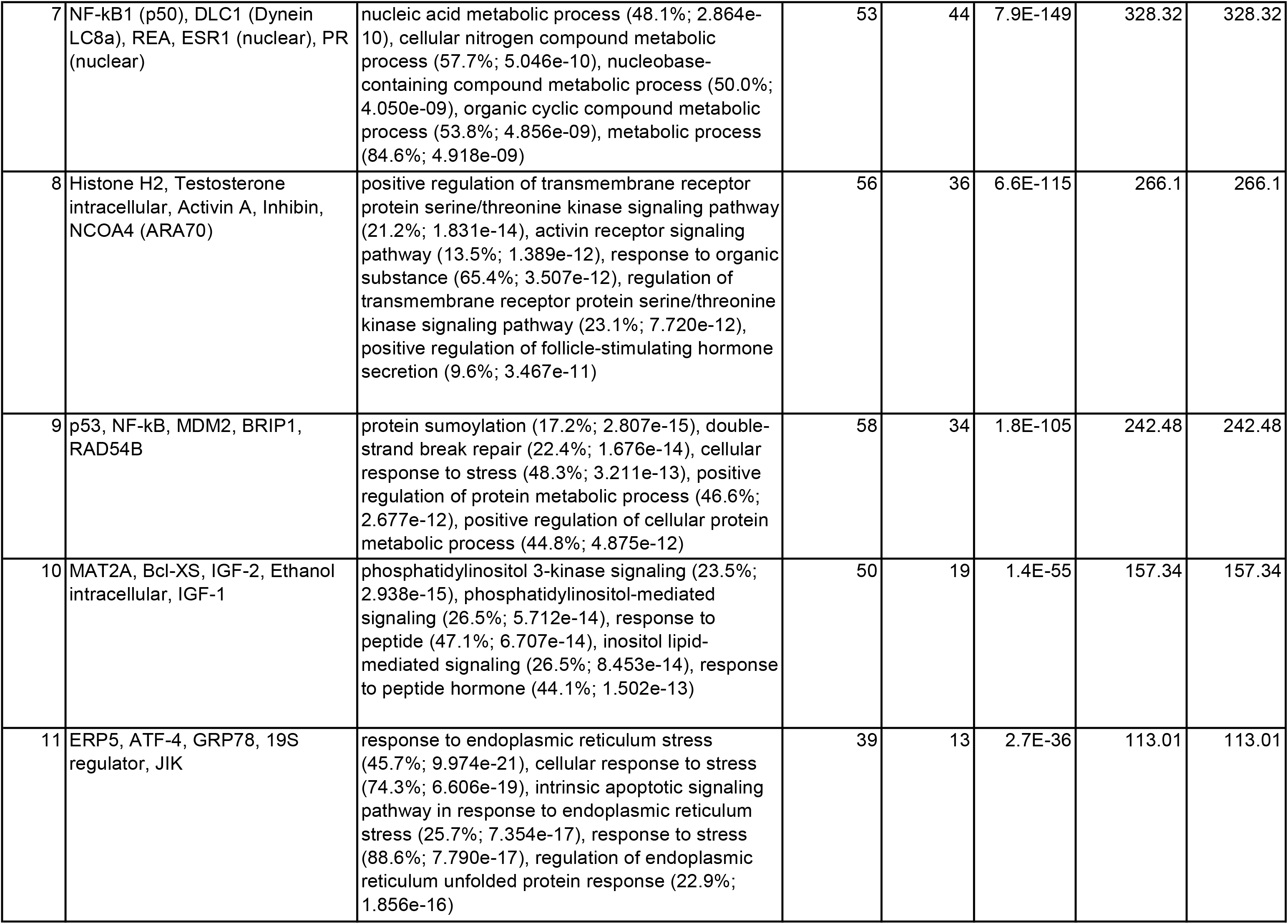
Top Network List (HMW-HA vs. Vehicle)

**Supplementary Table 6:** Human RNA-seq datasets used in the functional analysis..

| **Cell Line** | **GEO**<br>**Accession** | **Note** | **Control** | **Virus**<br>**infected** | **PubMed ID** |
| --- | --- | --- | --- | --- | --- |
| NHBE | GSE147507 | Series1 | Mock | SARS-CoV-2 | 32416070 |
| A549 | GSE147507 | Series5 | Mock | SARS-Cov-2 | 32416070 |
| A549-ACE2 | GSE147507 | Series16 | Mock | SARS-Cov-2 | 32416070 |
| HAE | GSE153970 |  | Mock | SARS-Cov-2 | 32699094 |
| Calu3 | GSE147507 | Series7 | Mock | SARS-Cov-2 | 32416070 |

**Supplemental Figure 1.**
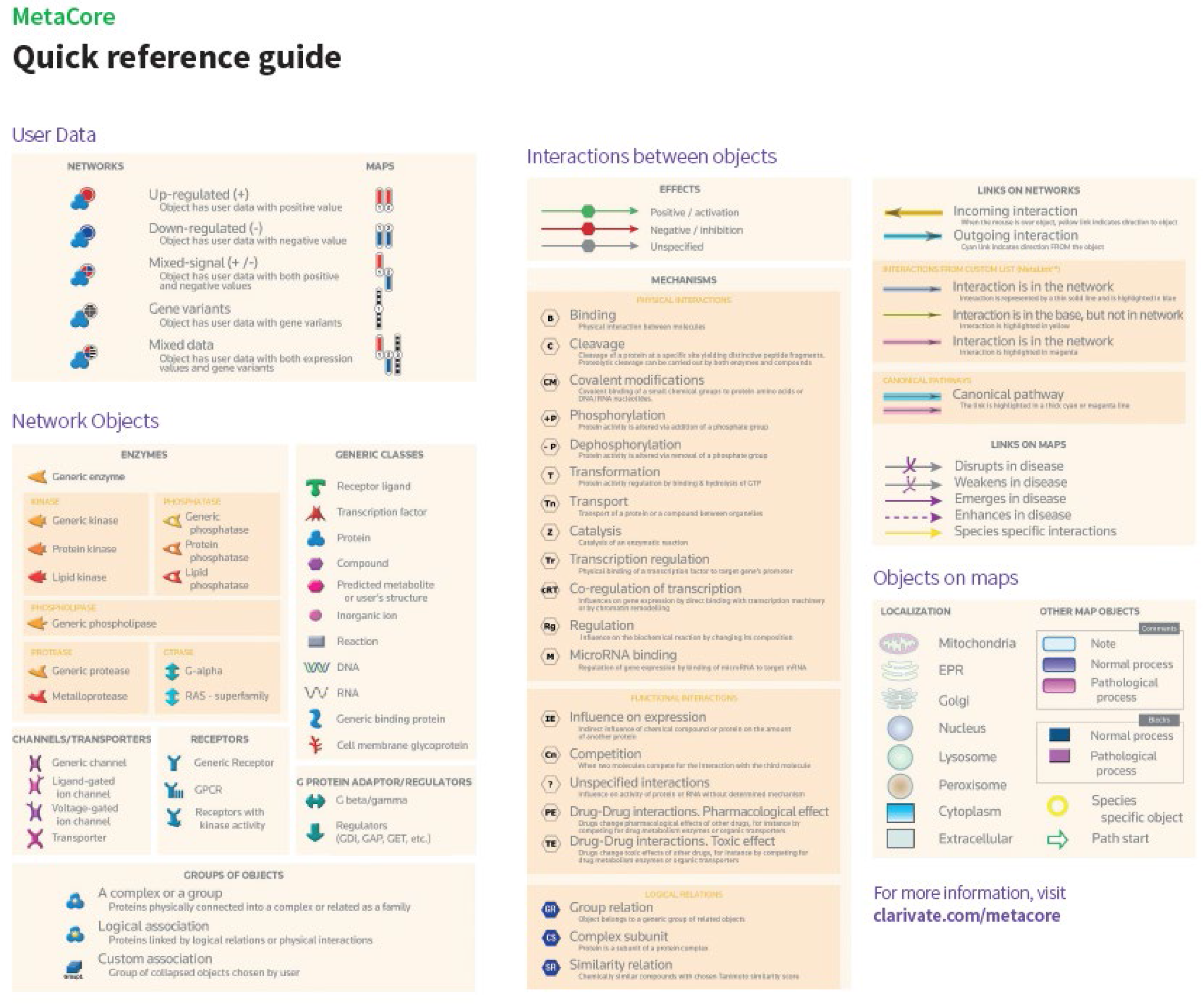
MetaCore Quicke Reference Guide: figure legend used to help follow the Network and Pathway Figures.

**Supplemental Figure 2.**
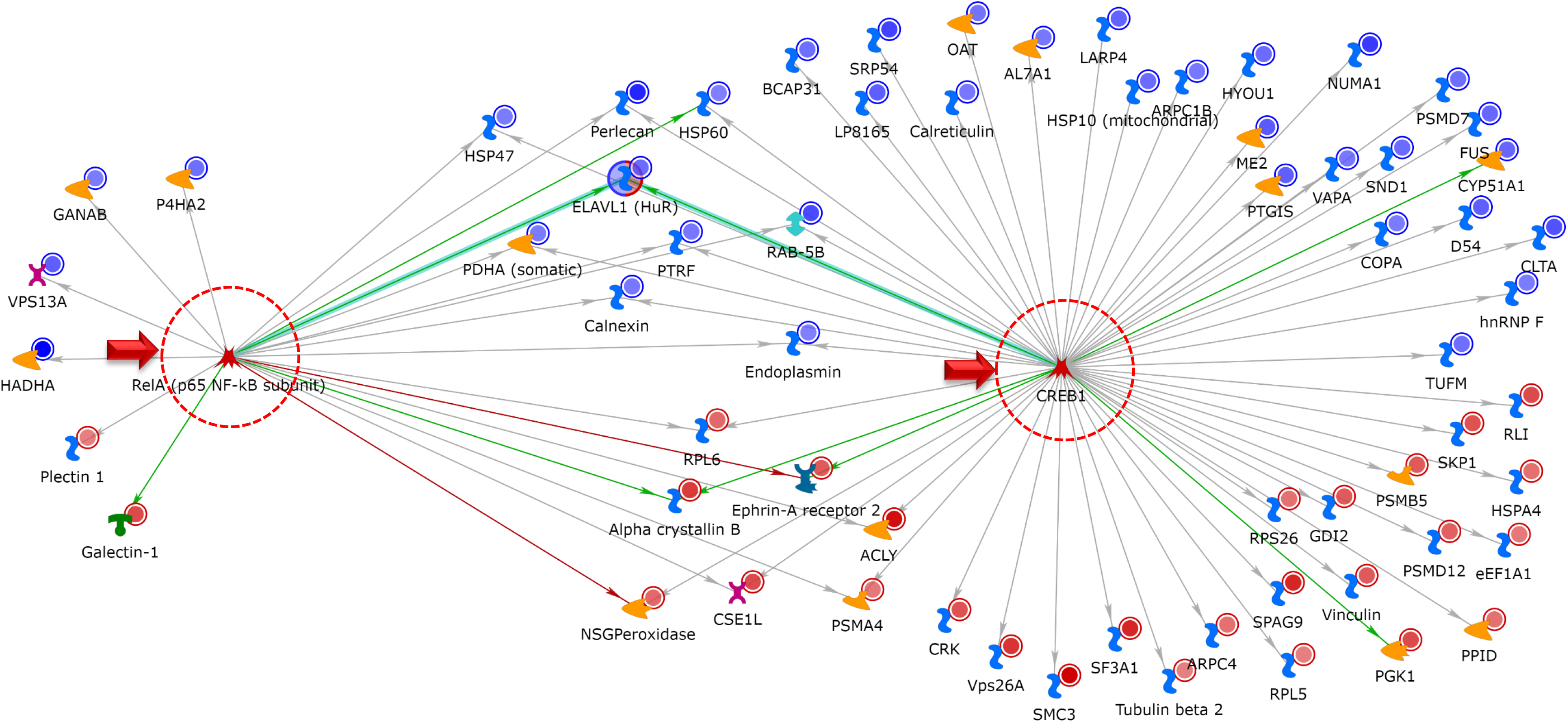
Top Network Map for SPIKE S1 Treatment; RelA and CREB1 presented as minor and major network hubs resepectively; blue circles indicate a decrease, while red circles indicate an increase in protein abundance (Rat L2 Cells; +/− S1 Tx vs. C for 24hrs).

**Supplemental Figure 3.**
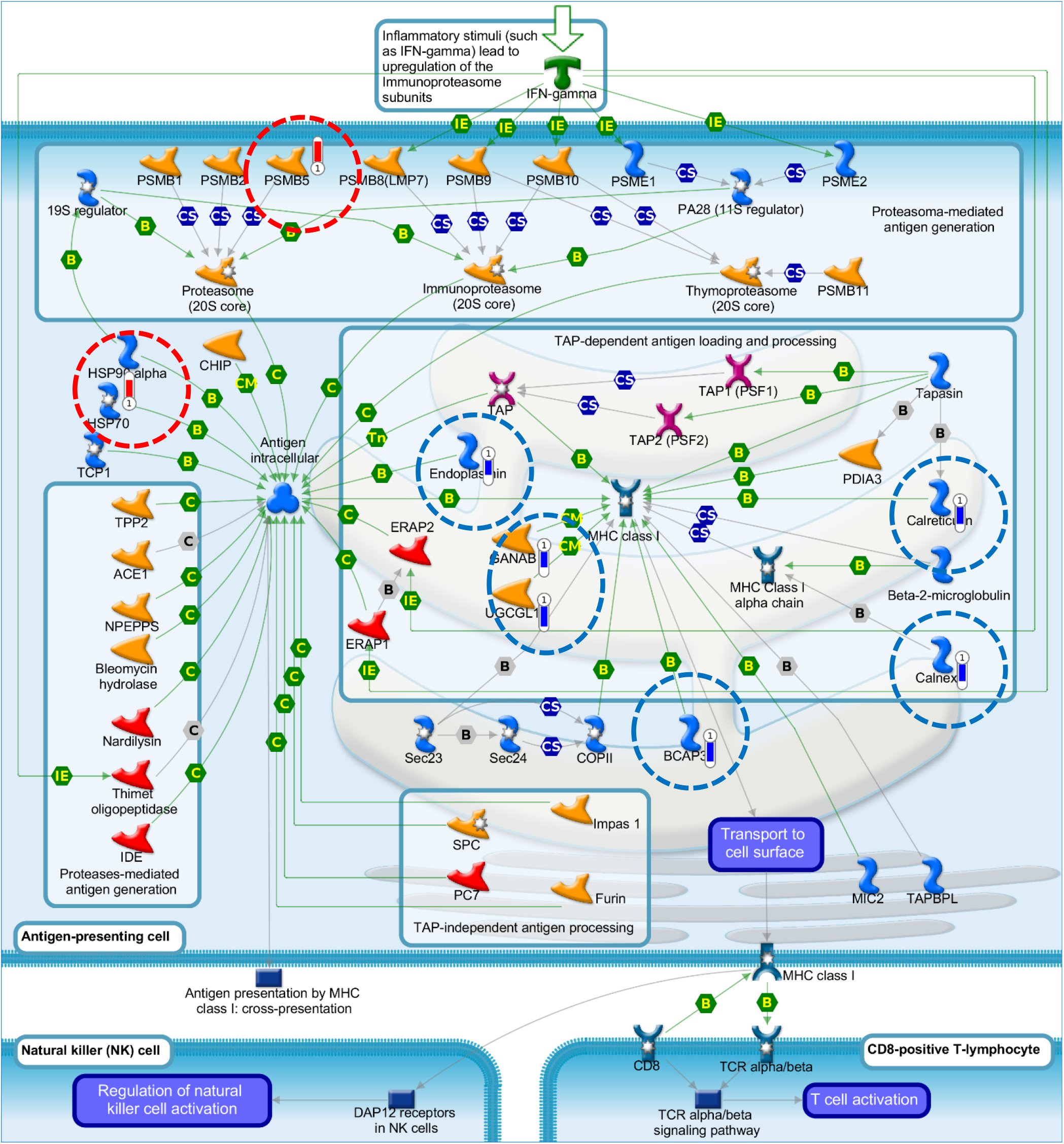
Top Pathway Map for SPIKE S1 Treatment: Immune response_Antigen presentation by MHC class I, classical pathway; blue bars indicate a decrease, while red bars indicate an increase in protein abundance (Rat L2 Cells; +/− S1 Tx vs. C for 24hrs).

**Supplemental Figure 4.**
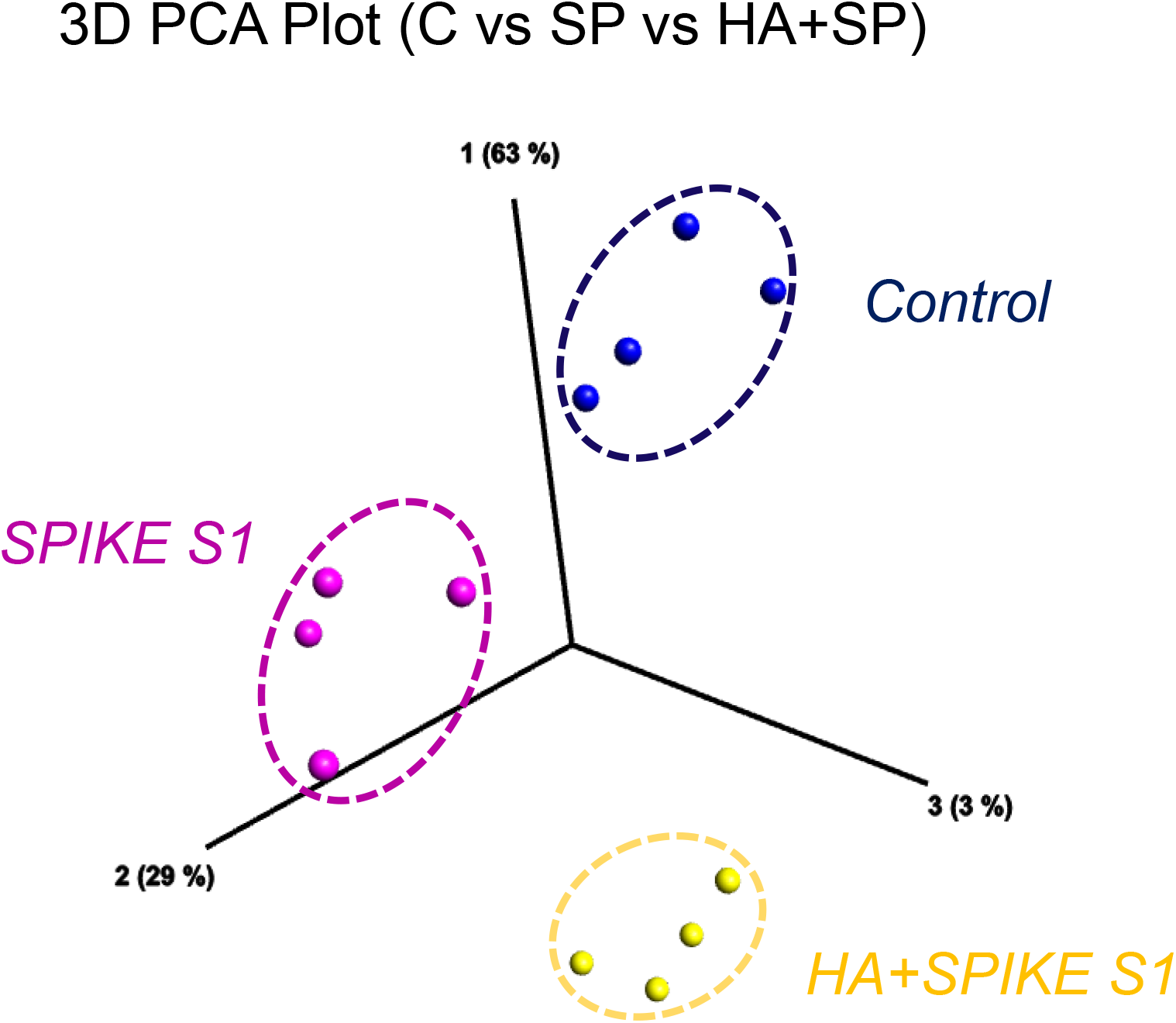
3D PCA Plot of Control (C) vs. S1-Tx (SPIKE S1) vs. Yabro® +S1-Tx (HA+SPIKE S1) in Rat L2 Cells, stemming from the most significant protein differences by ANNOVA (Analysis of variance) using the data matrix from SuppTable.1

**Supplemental Figure 5.**
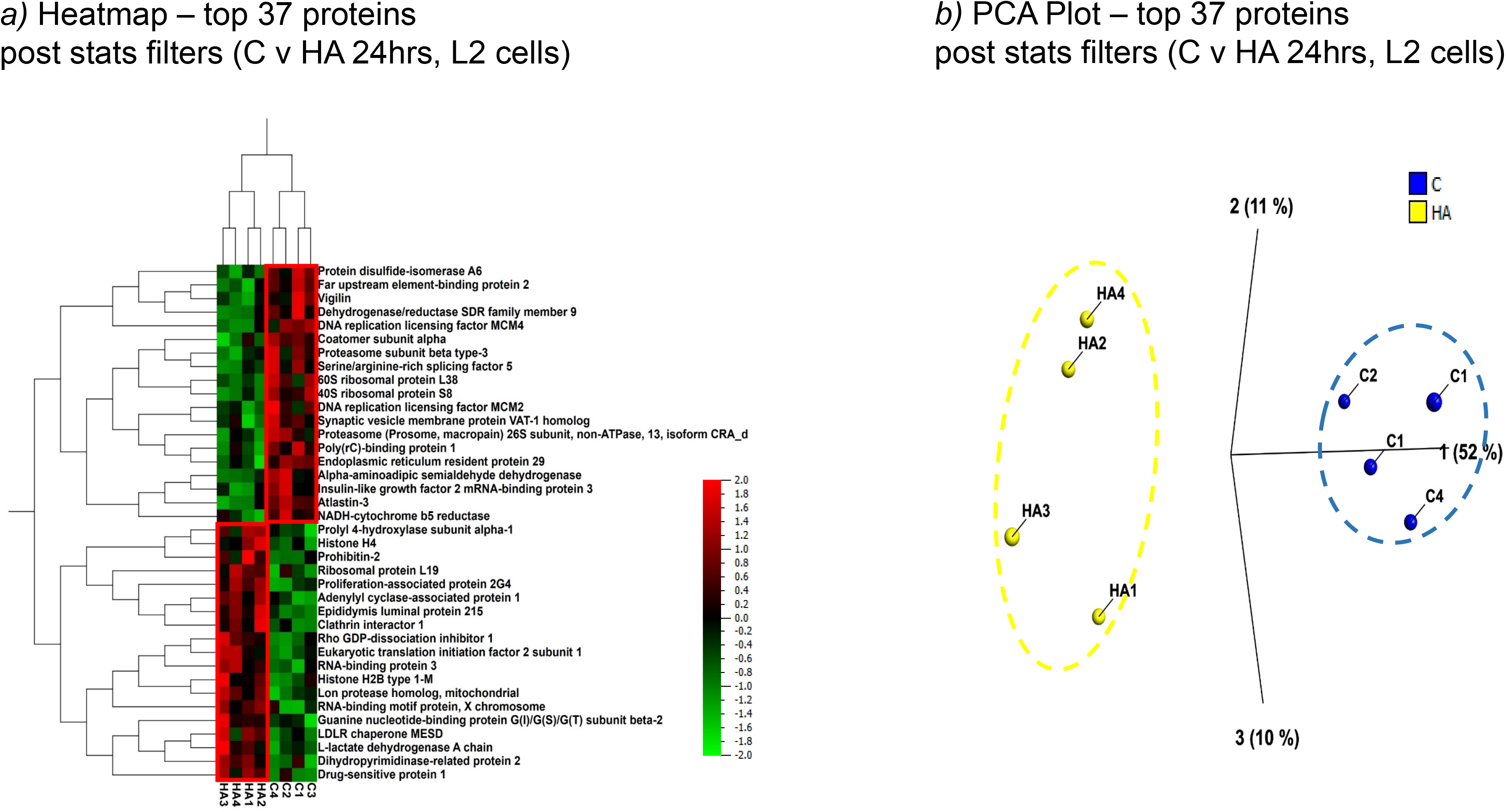
Multivariate analysis carried out on the 37 top most significantly changed proteins between Yabro® (HA) vs. Control (C) using the data from SuppTable.4. a) 2D HCA HeatMap, b) 3D PCA Plot.

